# scPolyA enables efficient, interactive and agent-guided alternative polyadenylation analysis in single-cell and spatial transcriptomes

**DOI:** 10.64898/2026.11.05.764757

**Authors:** Jingsen Yang, Haoyu Yang, Lixinyu Liu, Jiayu Wen

## Abstract

Alternative polyadenylation (APA) changes which regulatory sequences an RNA retains, creating a layer of cellular variation that gene-level expression profiles cannot resolve. Here we present scPolyA, a unified, AI-enabled framework that transforms conventional single-cell and spatial **3′**-end transcriptomes into quantitative maps of RNA-end choice. scPolyA connects *de novo* poly(A)-site discovery to cell- and spot-resolved measurements of terminal-site usage, **3′** untranslated region length and intronic polyadenylation, enabling differential APA analysis across cell states and tissue regions. Its integrated deep-learning module, DeepCPS, improves recognition of sparsely supported poly(A) sites and predicts cleavage positions at nucleotide resolution. An integrated graphical workspace brings parameter exploration, efficient rerunning and genome browsing together, while an APA-specialised AI agent translates natural-language research questions into executed analyses, verified outputs and evidence-guided interpretation. Across diverse single-cell protocols and spatial platforms, scPolyA achieves higher reference-atlas agreement than most evaluated *de novo* methods, while single-cell benchmarks demonstrate competitive runtime and memory use. Applied to cutaneous squamous cell carcinoma, scPolyA reveals recurrent **3′** untranslated region shortening in tumour keratinocytes across all ten patients and corresponding patterns in tumour-enriched regions, with tissue-level contrasts shaped by cellular composition. scPolyA extends conventional transcriptomics from gene abundance to RNA-end regulation, linking transcript choices to cellular states and tissue organisation.

## Introduction

Cells can alter the regulatory potential of an RNA without changing the overall expression of its gene. Alternative cleavage and polyadenylation (APA) makes this possible by selecting different transcript ends and changing which sequences are retained in mature RNAs [1, 2]. Within terminal exons, APA changes which 3′ untranslated region (3′ UTR) elements remain available for interactions with microRNAs and RNA-binding proteins, influencing RNA stability, localisation and translation. Cleavage within introns or alternative exons can also alter protein-coding potential [3–5]. RNA-end choice varies with cellular identity and state: neurons frequently express longer 3′ UTRs, while proliferating and transformed cells often favour proximal poly(A) sites [6–8]. In heterogeneous tissues, understanding this regulation requires knowing which transcript ends different cell populations use and how those choices change, even when gene-level expression is similar.

An extraordinary resource for answering these questions is already being generated, largely for another purpose. Most droplet-based and many plate- and microwell-based single-cell RNA-sequencing assays, together with numerous spatial transcriptomics technologies, use oligo(dT) priming to capture transcripts near their 3′ ends [9–17]. Public resources now encompass tens of millions of cells, yet these libraries are still analysed predominantly as measurements of gene abundance [18–20]. Their potential for APA analysis lies in an informative asymmetry. A small subset of reads extends into an untemplated poly(A) tail, providing comparatively precise evidence of RNA-end positions, whereas the far more abundant neighbouring reads supply the depth needed to quantify usage but locate cleavage less precisely. Oligo(dT) priming at genomic A-rich tracts can further produce apparent ends that resemble cleavage-associated reads [21]. Recovering APA from conventional transcriptomes is consequently not simply a peak-detection problem: it requires reconciling evidence of *where transcripts end* with evidence of *how often each end is used*.

Existing methods have established complementary ways to study APA in conventional 3′-end libraries. Reference-guided approaches, including MAAPER and scUTRquant, quantify established transcript ends [22, 23]. *De novo* methods identify candidates from coverage patterns (scAPA, Sierra and scAPAtrap), fragment distributions (SCAPE), sequence-filtered peaks (SCAPTURE) or poly(A)-tail reads (polyApipe) [24–29]. Other approaches exploit informative first-read sequences, including scTail and scPAISO, the latter also supporting Read2-only analysis [30, 31]. Configurable workflows and spatial approaches further broaden the settings in which APA can be studied [32–34]. Together, these advances show that conventional libraries can support site discovery and usage quantification. However, an important challenge remains in connecting the evidence used to locate RNA ends with the intervals used to quantify their usage. polyApipe, for example, identifies candidate ends from poly(A)-tail reads but assigns unique molecular identifiers (UMIs) within a configurable upstream window (250 nt by default) [29], whereas coverage-based approaches define candidate regions from read accumulation, which may locate cleavage positions less precisely. Because site boundaries determine which UMIs are assigned to each alternative end, these differences can affect downstream APA measurements. Systematic benchmarking has indeed revealed substantial variation in reported poly(A)-site coordinates and usage estimates across methods [35]. This motivates an approach that anchors candidate ends to direct poly(A)-tail evidence while using local coverage to define adaptive UMI-counting intervals.

Defining sites is only part of the challenge. A gene may contain several candidate ends in a pooled catalogue, yet individual cells may have too few terminal UMIs to estimate how frequently each is used. Relaxing discovery thresholds can greatly increase the number of detected sites and multi-site genes without a corresponding gain in measurements with sufficient molecular coverage. The balance between site recovery, direct evidence and quantitative resolution varies with sequencing depth, library chemistry and the cellular or spatial scale of analysis. The number of recovered sites and the number of measurable APA choices are therefore distinct outcomes.

Deep learning offers an opportunity to assess candidate poly(A) sites and refine cleavage positions when direct tail-read evidence is sparse, as often occurs in single-cell and spatial transcriptomes. Existing methods, including APARENT, DeepPASTA and DeepPASS, have demonstrated that local sequence contains predictive information about polyadenylation, while scDeepAPA extends sequence learning to single-cell APA analysis and PolyaID predicts cleavage distributions at nucleotide resolution [28, 36–39]. For candidates supported by very few tail reads, two questions are particularly relevant. Can sequence and local coverage provide additional evidence of a poly(A) site, including when internal priming at genomic A-rich tracts may produce apparent ends? Can sequence information refine cleavage-position estimates when only a few read endpoints are observed? These complementary tasks offer a way to assess candidate-site support and localise cleavage more precisely while keeping predictions distinct from observed RNA-end usage.

Putting these analyses into practice requires more than a single pipeline run. Site-selection choices need to be revisited, their effects on downstream measurements compared and individual candidates examined against their supporting reads. General workflow systems and configurable APA pipelines provide foundations for reproducible computation [32]. Interactive analysis and domain-guided AI assistance offer further ways to connect biological questions with appropriate workflow execution, output checks and evidence-based interpretation [40–42]. Together, these capabilities can make it easier to establish which APA comparisons the available molecules support and how the conclusions depend on analytical settings.

Here we introduce scPolyA (spatial and single-cell poly(A)-site analysis), a unified framework for discovering, quantifying and interpreting RNA-end choice in conventional single-cell and spatial 3′-end transcriptomes (Fig. 1). scPolyA discovers sites *de novo* from pooled poly(A)-tail evidence and uses coverage-adaptive intervals to assign UMIs, retaining cell- and spot-resolved measurements of terminal-site usage, 3′ UTR length and intronic polyadenylation. We developed DeepCPS, an integrated deep-learning module that uses sequence and coverage to assess sparsely supported candidate sites and learns from observed tail-read endpoints to predict cleavage positions at nucleotide resolution. A parameter-aware task graph enables efficient reanalysis when discovery settings change, while a graphical workspace connects parameter exploration with genomic evidence and model scores. An APA-specialised, domain-guided AI assistant translates natural-language research questions into executed analyses, verified outputs and evidence-guided interpretation. Together, these capabilities extend conventional single-cell and spatial transcriptomics beyond gene abundance to reveal how RNA-end regulation varies among cell populations and tissue regions.

**Fig. 1.**
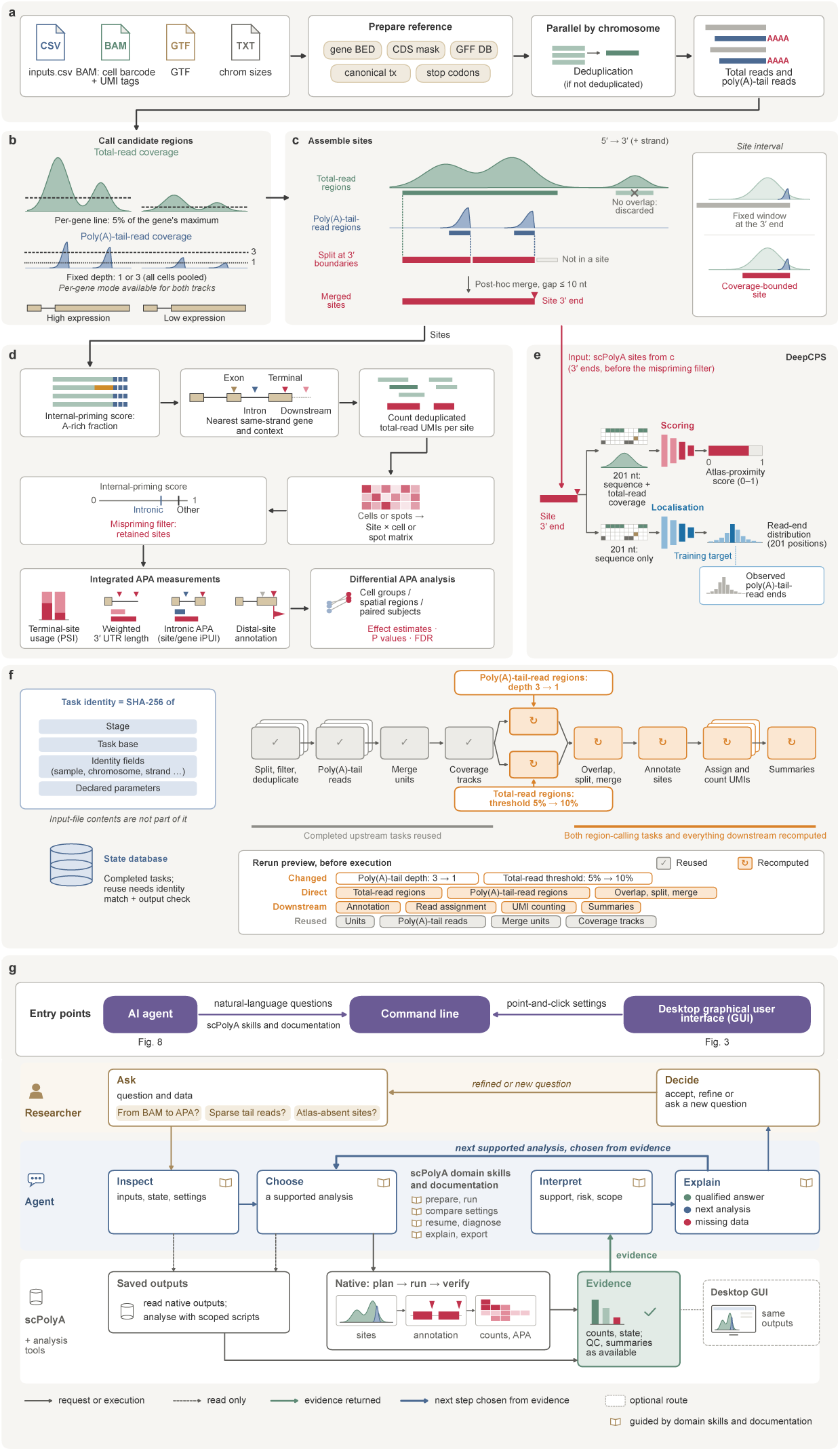
From sparse cleavage evidence to measurable RNA-end choice. scPolyA connects two complementary signals in conventional 3′-tag libraries: poly(A)-tail reads provide evidence of transcript endpoints, while the larger population of neighbouring reads enables their use to be quantified. The same framework links site construction, APA measurements and interpretation across single cells and spatial locations. **a,** Starting with barcoded, UMI-tagged alignments and gene annotations, the pipeline prepares chromosome-specific references and deduplicated reads while retaining cell or spatial-barcode identities. Total reads and reads containing poly(A) tails provide separate coverage tracks. **b,** Strand-specific coverage pooled across barcodes identifies candidate regions on each track. The example uses a total-read threshold relative to the maximum coverage of each gene and tail-read depth thresholds of 1 or 3, illustrating how evidence stringency can be varied. **c,** The two tracks are brought together to define quantifiable sites: a total-read region is retained when it overlaps tail-read evidence, tail-read 3′ boundaries locate candidate endpoints, and total-read coverage determines the interval available for molecule counting. Nearby intervals can be merged. The inset contrasts a coverage-defined interval with a fixed-length upstream window. **d,** The resulting sites are assessed for internal priming, assigned to genes and transcript contexts, and counted using unique molecular identifiers (UMIs) for each cell or spatial unit. The site-by-barcode matrix supports terminal-site usage (PSI), usage-weighted 3′ UTR length, intronic polyadenylation indices (iPUI) and distal-site annotations; cell-group, spatial-region or patient labels then support differential APA analysis. **e,** DeepCPS assesses candidates with limited tail-read support using two models. A sequence-and-coverage scoring model estimates reference-atlas support, while a sequence-only localisation model predicts the distribution of cleavage-associated read endpoints. Its most probable position provides the predicted cleavage coordinate. **f,** As discovery settings change, the task graph distinguishes completed computations that can be reused from site-dependent steps requiring reanalysis. The schematic changes both the poly(A)-tail-read depth (3 to 1) and total-read threshold (5% to 10%); grey check marks (✓) indicate anticipated reuse and orange circular arrows (⟳) recomputation. **g,** Biological questions can be taken through the same analytical outputs using the command line, desktop interface (Fig. 3) or an APA-specialised AI assistant. Guided by scPolyA documentation and domain skills, the assistant can inspect saved evidence or follow a native plan–run–verify pathway, then return an interpretation, a supported next analysis or a description of missing information.

## Results

### scPolyA turns sparse cleavage evidence into cell- and spot-resolved APA measurements

Conventional 3′-end single-cell RNA sequencing and oligo(dT)-based spatial transcriptomics present an unusual imbalance of evidence: reads extending into an untemplated poly(A) tail help pinpoint transcript ends, whereas the much larger read population provides the depth needed to quantify their use. Across the examined libraries, detectable tail reads accounted for approximately 0.03–7.6% of eligible alignments (Supplementary Fig. 2). The challenge is therefore to connect *where* RNA ends occur with *how often* those ends are used. We developed scPolyA (spatial and single-cell poly(A)-site analysis) to unite these complementary signals in site discovery and cell- and spot-resolved quantification (Fig. 1).

Starting from barcoded, UMI-tagged alignments and gene annotations, scPolyA pools read evidence across cells or spatial locations to discover poly(A) sites *de novo*, without requiring a reference poly(A)-site atlas as input (Fig. 1a–c). Poly(A)-tail-read evidence anchors candidate 3′ ends, while total-read coverage defines variable-width intervals for UMI counting rather than fixed windows. After assessing internal priming and assigning sites to genes and transcript contexts, scPolyA quantifies site-specific UMIs in individual cells or spatial units (Fig. 1d; Supplementary Fig. 1a). Pooled evidence thus establishes the candidate ends, while barcode-resolved counting measures their use.

The resulting counts allow the choice between RNA ends to be examined at several levels. Alongside site-by-cell or site-by-spot UMI matrices, scPolyA directly outputs terminal-site usage (PSI), usage-weighted 3′ UTR length, and site- and gene-level intronic polyadenylation indices (iPUI). Distal-site annotations support downstream calculation of distal-site usage (Supplementary Fig. 1c). Together with cell-group, spatial-region and subject labels, these outputs provide the basis for differential APA testing, including paired-subject comparisons, with gene-level effect estimates and false-discovery-rate-adjusted significance (Fig. 1d).

Assessing the sites underlying these measurements is particularly important when tail-read support is limited. To address this challenge, we integrated DeepCPS into scPolyA, providing deep-learning-based assessment of candidate sites and prediction of cleavage positions at nucleotide resolution (Fig. 1e). By incorporating sequence and read-coverage information, DeepCPS complements read-based discovery with additional evidence for site selection and localisation.

Site assessment then leads to a practical question: which alternative ends can be compared within individual cells or spatial units? Lowering a tail-read threshold can reveal additional sites and multi-site genes, while the number of measurable within-gene choices also depends on the number and distribution of site-specific UMIs within each cell or spatial unit. scPolyA supports this exploration through a parameter-aware task graph that parallelises independent operations and reruns only the computational steps affected by changes to site-selection settings (Fig. 1f). Users can therefore revisit discovery thresholds and examine their consequences for both site evidence and measurable APA.

This iterative workflow is accessible through three complementary interfaces to the same pipeline and its outputs. The command line supports scripted execution, while an integrated graphical user interface (GUI) brings flexible parameter selection, rerun previews and comparative genome browsing into one workspace. An APA-specialised, domain-skill-guided AI assistant connects natural-language research questions to analysis planning, execution, output verification and evidence-grounded interpretation (Fig. 1g). In this framework, a poly(A) site is more than a reported coordinate: it is linked to the evidence supporting its placement, the molecules assigned to it and the analytical choices underlying its interpretation.

### DeepCPS extracts cleavage-site information from sparsely supported candidates

Poly(A)-site discovery presents a challenge when candidate sites are supported by very few tail reads. Requiring multiple reads may exclude genuine sites, while retaining sparsely supported candidates calls for additional evidence to assess their validity. To establish how frequently poly(A)-site discovery encounters sparse read support, we examined eight single-cell datasets spanning different sequencing protocols. At a minimum tail-read support threshold of one, a median of 38% of candidate sites (range, 19–59%) were supported by exactly one tail read. Across datasets, a median of 34% of these single-read candidates lay within 25 nt of a reference poly(A) site in PolyA_DB 3 [43] or PolyASite 2.0 [44], compared with 90% of candidates supported by more than ten tail reads (Fig. 2a). Among single-read candidates, reference-atlas agreement ranged from 17–22% in droplet datasets to 46–78% in plate and microwell datasets. Thus, the same tail-read count was associated with markedly different levels of reference-atlas agreement across protocols. We asked whether the sequence and read coverage surrounding each candidate could provide additional evidence of a transcript end, particularly when direct tail-read support was limited.

**Fig. 2.**
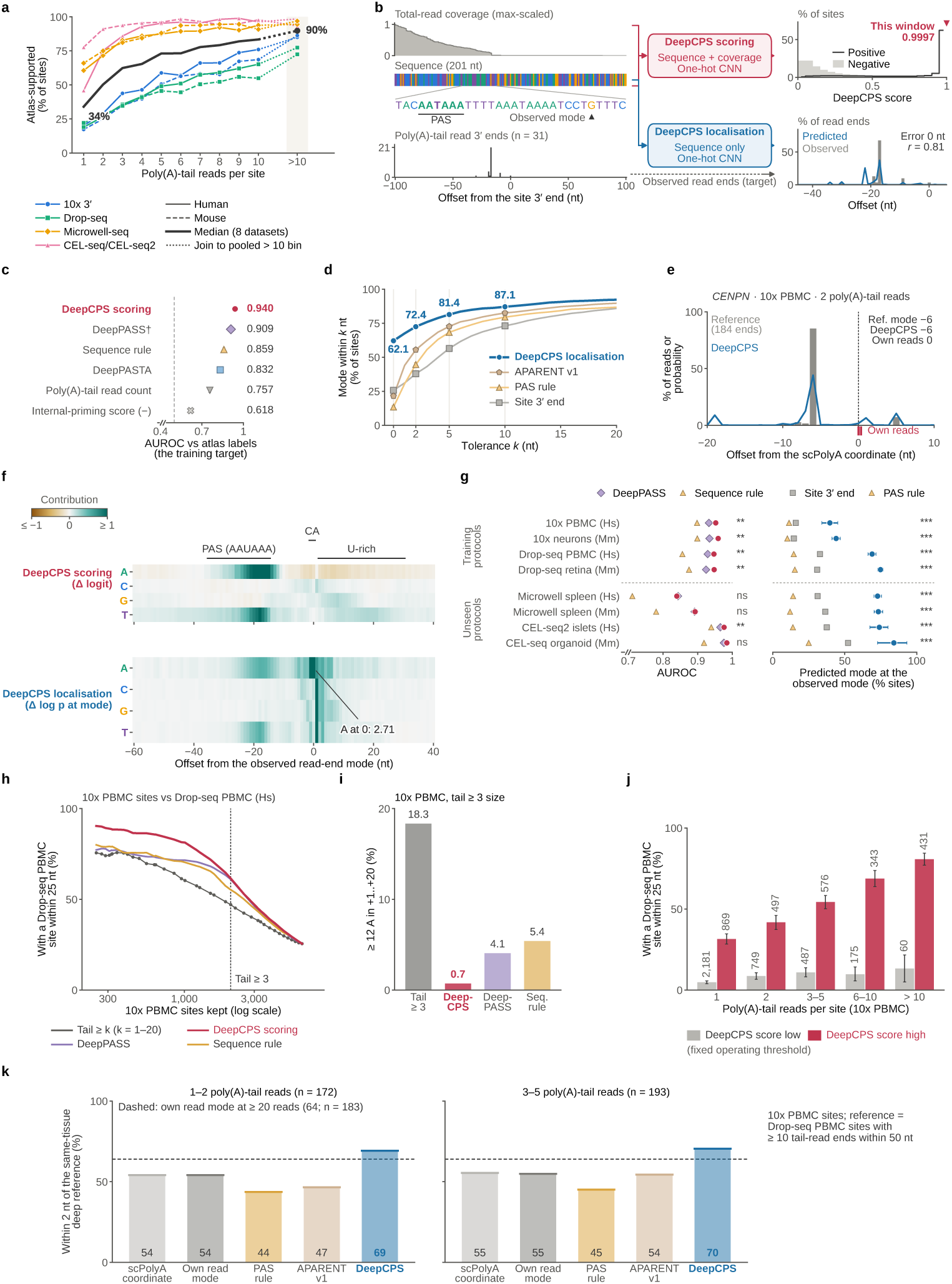
DeepCPS draws additional information from sparse evidence of transcript ends. A candidate supported by only one or two poly(A)-tail reads may be difficult to assess or locate precisely. DeepCPS addresses these uncertainties separately, using candidate-centred sequence and coverage to rank reference-site support and sequence-derived endpoint distributions to estimate cleavage position. Independent datasets test whether these predictions remain informative beyond the original candidates. **a,** Across eight single-cell datasets, the median fraction of candidates within 25 nt of a reference poly(A) site increases from 34% with one tail read to approximately 90% with more than ten. Each line represents a dataset, with the black line showing the median; candidates precede internal-priming filtering. **b,** The held-out *MAZ* example illustrates the separate DeepCPS models. Site scoring uses a 201-nt sequence window together with local total-read coverage (crimson); localisation uses sequence alone to predict the distribution of tail-read endpoints (blue), with its mode providing the predicted coordinate. **c,** On held-out chromosomes, DeepCPS discriminates candidates near reference sites with AUROC 0.940, compared with 0.909 for DeepPASS and 0.832 for DeepPASTA. The plot also includes a fitted sequence rule and simpler read-evidence rankings. Symbols show bootstrap intervals (mostly smaller than the symbols); † marks DeepPASS atlas overlap with the test chromosomes. **d,** Cleavage-site localisation is evaluated against the observed modal tail-read endpoint at 4,115 well-supported sites. DeepCPS predicts this position exactly at 62.1% of sites and within 2 nt at 72.4%; the curves compare increasingly permissive nucleotide tolerances with APARENT, a PAS-offset rule and the original scPolyA coordinate. **e,** At a *CENPN* candidate with two tail reads in 10x PBMC, the DeepCPS prediction coincides with the modal end-point in a deeper Microwell-seq spleen sample, despite a difference from the two PBMC read ends. **f,** Single-nucleotide substitutions show that upstream AAUAAA bases support candidate scoring and bases around the observed cleavage position, including CA, contribute to localisation. Contribution is the output for the original base minus its mean substituted output; positive values indicate a reduction following substitution. Scoring logits and positional log probabilities have different scales. **g,** Frozen models were evaluated on held-out chromosomes from four droplet datasets used in model development and four plate or microwell datasets not used for training. Cleavage localisation transferred well, whereas the scoring advantage over DeepPASS varied by protocol. The left plot reports atlas-proximity AUROC and the right the proportion of exactly predicted modal read endpoints. Significance symbols follow the tests described in Methods. **h,** To assess cross-dataset reproducibility, candidate poly(A) sites on held-out chromosomes in 10x PBMC data were ranked by each method and compared with independently generated Drop-seq PBMC data. A candidate was considered reproduced if a same-strand Drop-seq site supported by at least one tail read was detected within 25 nt. Curves show the proportion of reproduced sites as increasing numbers of candidates are retained. The dashed line marks the size of the tail-read-threshold-3 catalogue (2,072 sites). **i,** At matched catalogue size (2,072 sites), DeepCPS selection retained fewer downstream A-rich candidates than tail-read-count selection (0.7% versus 18.3%). The sequence-rule baseline also penalises downstream adenosines, contributing to its low A-rich fraction. **j,** Among 10x PBMC candidates supported by exactly one tail read, DeepCPS scores distinguished sites with different levels of cross-dataset reproducibility. Candidates were divided into high- and low-scoring groups using a threshold selected on validation chromosomes (chr5 and chr12) to match the PolyA_DB 3 reference-support rate of the catalogue requiring at least three tail reads. Using the reproducibility criterion in panel h, 31.5% of high-scoring candidates were detected in Drop-seq PBMC, compared with 4.9% of low-scoring candidates. Bars show 95% bootstrap intervals, and numbers indicate candidate counts. **k,** At sites supported by one or two tail reads, DeepCPS places 69% of endpoints within 2 nt of a deeper same-tissue read-end mode, compared with 54% for the original scPolyA coordinate. The second group contains sites with three to five reads; the dashed line shows the performance of the site’s own read-end mode at well-covered sites.

To address this question, we developed DeepCPS (Deep learning of Cleavage and Polyadenylation Sites) to learn two complementary properties of candidate RNA ends: *their resemblance to established poly(A) sites and the nucleotide at which cleavage is most likely to occur* (Fig. 2b). Its scoring model combines one-hot-encoded nucleotide sequence with total-read coverage in a candidate-centred 201-nt window. Training on scPolyA-generated sites exposes the model to the sparse, intronic and A-rich candidates encountered during *de novo* discovery. For model training, we assembled 217,102 candidates from four droplet datasets, assigning positive scoring labels to sites within 25 nt of an atlas entry and negative labels to those more than 100 nt away. To pinpoint the cleavage nucleotide, we trained a separate localisation model to predict the distribution of poly(A)-tail-read endpoints from the surrounding sequence, using well-supported sites as training examples. The most probable position provides a sequence-based estimate of the cleavage site, including for candidates with sparse tail-read support. Chromosomes 2, 9 and 16 were held out for testing, with chromosomes 5 and 12 reserved for model selection and calibration (Supplementary Fig. 3a; Methods). The training strategy therefore links each prediction to the evidence it is intended to recover.

We first tested whether candidate context could distinguish sites beyond their observed read support. On held-out chromosomes, DeepCPS achieved a scoring area under the receiver operating characteristic curve (AUROC) of 0.940 (95% bootstrap interval 0.937–0.942), compared with 0.909 for DeepPASS [28] and 0.832 for DeepPASTA [37]. Simple rankings by tail-read count or lower internal-priming risk reached 0.757 and 0.618, respectively (Fig. 2c). The advantage over DeepPASS persisted under locus-level resampling (ΔAUROC = 0.030; *P*_BH_ ≤ 0.001) and across all four test datasets and examined genomic contexts (Supplementary Fig. 3b). The single-read candidates made the added information particularly clear: despite identical tail-read counts, DeepCPS distinguished atlas-supported candidates with an AUROC of 0.898, compared with 0.863 for DeepPASS. To make DeepCPS scores interpretable as probabilities of reference-atlas support, we calibrated them using isotonic regression, which maps model scores to observed frequencies of atlas matches. Calibration was fitted on validation chromosomes and improved agreement between predicted probabilities and observed atlas support on held-out chromosomes, reducing expected calibration error from 0.034 to 0.008 (Supplementary Fig. 3c).

To understand what drove this discrimination, we separated the contributions of sequence and coverage through matched ablations (Supplementary Fig. 4a,b). Sequence alone achieved an AUROC of 0.932 and coverage alone 0.825. Combining sequence and coverage increased AUROC to 0.940, with the greatest improvement for candidates in non-terminal exons (0.716 to 0.741; Supplementary Fig. 4b). Sequence therefore supplied most of the recognition signal, while coverage added information about how the candidate appeared in the sequenced library. The one-hot sequence-and-coverage model also outperformed variants incorporating frozen embeddings from the pretrained genomic language model Nucleotide Transformer v2 [45] (AUROC 0.932 with coverage and 0.927 without). We therefore selected this compact sequence-and-coverage architecture, which achieved the best scoring performance in these comparisons while avoiding the additional computational time required to generate pretrained embeddings.

Having established how well DeepCPS recognised candidate sites, we next asked whether it could predict cleavage positions at nucleotide resolution. We first evaluated localisation at 4,115 atlas-supported test sites with at least ten tail reads, where the most frequently observed tail-read endpoint provided a better-supported reference for assessing positional accuracy. DeepCPS matched this position exactly at 62.1% of sites, compared with 21.5% for APARENT [36], 13.4% for a rule based on the offset from a canonical polyadenylation signal (PAS) and 25.7% for the original scPolyA site boundary (Fig. 2d). Accuracy reached 72.4% within 2 nt, with similar results on the plus and minus strands (72.0% and 73.0%; exact McNemar *p <* 10^−16^ against APARENT). The *CENPN* example brings this positional inference into focus: at a 10x PBMC (peripheral blood mononuclear cell) candidate supported by two tail reads, DeepCPS predicted the endpoint observed in a deeper Microwell-seq spleen reference, at a different position from the candidate’s own read ends (Fig. 2e; further examples in Supplementary Fig. 5). This illustrates how sequence can supplement a sparsely sampled cleavage position.

To investigate which sequence features DeepCPS uses to recognise poly(A) sites and locate cleavage positions, we performed *in silico* mutagenesis by systematically substituting individual nucleotides and measuring how each change affected model predictions. For each substitution, we kept the read-coverage input unchanged and compared the model output with that obtained from the original sequence. In the site-scoring model, substitutions within the upstream AAUAAA polyadenylation signal reduced the scoring logit by an average of 1.81 per base (95% interval, 1.77–1.86), indicating that the intact motif strongly supported candidate-site recognition. In contrast, replacing downstream adenosines increased the scoring logit by an average of 0.16, suggesting that these bases contributed negatively to site scores in their original sequence context. In the cleavage-localisation model, substitutions within the CA dinucleotide reduced the log probability assigned to the observed cleavage position by an average of 0.87, consistent with the known preference for cleavage immediately after CA [39] (Fig. 2f). Together, these results show that DeepCPS captures established polyadenylation sequence features, using the upstream polyadenylation signal to assess candidate sites and local sequence preferences to identify cleavage positions.

We next evaluated how well DeepCPS generalises across species and sequencing platforms. In cross-species and cross-protocol evaluations, DeepCPS achieved scoring AUROCs of 0.896–0.940 (Supplementary Fig. 4c). Applied without refitting to four plate and microwell datasets, DeepCPS localised 72.8–84.1% of observed endpoints to the exact nucleotide, compared with 31.2–52.3% for the original site boundary (exact McNemar *p ≤* 5.2 × 10^−4^ in each dataset; Fig. 2g). Scoring AUROCs ranged from 0.840 to 0.984, with differences from DeepPASS of −0.003 to +0.011, compared with +0.018 to +0.025 on the training protocol families. The contrast between the two tasks is informative: sequence-based cleavage localisation transferred particularly well, whereas the relative scoring advantage varied with protocol. This suggests that positional sequence information is broadly portable, while candidate ranking also reflects how each assay generates coverage and samples RNA ends.

We next asked whether DeepCPS could improve poly(A)-site selection for downstream APA analysis. A common approach requires a minimum number of supporting tail reads, which may exclude genuine sites with sparse read support. We compared DeepCPS-based selection with a threshold requiring at least three tail reads, choosing the DeepCPS score threshold on validation chromosomes to achieve similar reference-atlas agreement. Across eight test datasets, DeepCPS retained more candidate sites in every dataset (median 3,081 versus 1,945; *P*_BH_ = 0.023), while maintaining comparable atlas-support rates (median 76.9% versus 79.9%; *P*_BH_ = 0.945). At matched catalogue size, DeepCPS selected fewer intronic and downstream A-rich candidates, reducing the median fraction of downstream A-rich sites from 0.4% to 0.1% (Supplementary Fig. 6a,b). We then asked whether improved site selection also increased the precision of APA quantification by computationally reducing UMI depth and comparing distal-site usage estimates with their full-depth values. Sampling errors remained similar across filtering strategies, indicating that usage precision primarily depends on the number of observed molecules (Supplementary Fig. 6c). These results show that DeepCPS enables the retention of more candidate poly(A) sites while maintaining comparable reference-atlas agreement, with the precision of their usage measurements continuing to depend on molecular coverage.

We next asked whether poly(A) sites prioritised by DeepCPS could also be detected in an independently generated sequencing dataset, providing additional evidence of site reproducibility beyond reference-atlas agreement. To test this, we identified candidate poly(A) sites in 10x PBMC data and examined whether corresponding sites were detected in a separate Drop-seq PBMC dataset. Starting from the same candidates on held-out test chromosomes, we ranked sites using DeepCPS, DeepPASS, a sequence-based rule or tail-read count. To ensure a fair comparison, we selected the same number of sites (2,072) from each ranking and measured the fraction with a corresponding site within 25 nt in the Drop-seq data. DeepCPS-selected sites showed 61.5% cross-dataset agreement, compared with 47.1% for tail-read selection, 55.3% for the sequence-based rule and 61.0% for DeepPASS (Fig. 2h; Supplementary Fig. 6d). DeepCPS also selected substantially fewer downstream A-rich sites than tail-read selection (0.7% versus 18.3%; Fig. 2i), consistent with reduced selection of potential internal-priming artefacts. These results show that DeepCPS prioritises candidate poly(A) sites with stronger independent experimental support than tail-read count alone.

We further examined whether DeepCPS could improve the assessment of poly(A) sites supported by very few tail reads. We first asked whether DeepCPS could distinguish reproducible poly(A) sites among candidates supported by exactly one tail read. Using a score threshold fixed on validation chromosomes, 31.5% of high-scoring candidates were detected in the independent Drop-seq PBMC dataset, compared with only 4.9% of low-scoring candidates (Fig. 2j). We next examined whether DeepCPS could improve cleavage-position estimates for sparsely supported sites. Among 172 sites supported by one or two tail reads and evaluated against a deeply sequenced reference from the same tissue, DeepCPS predicted cleavage positions within 2 nt of the reference in 69.2% of cases, compared with 54.1% using the original scPolyA coordinates (Fig. 2k). Together, these findings show that DeepCPS extracts useful information beyond tail-read counts, improving both the prioritisation of sparsely supported poly(A) sites and the precision of their predicted cleavage positions.

### An integrated GUI brings flexible APA analysis and genome browsing together

Read evidence and DeepCPS predictions provide complementary views of candidate RNA ends. To bring these together in a unified workflow, we developed an integrated graphical workspace for running, reviewing and exploring scPolyA analyses (Fig. 3a). Users can configure and execute analyses, access site counts and APA measurements, and inspect RNA-end evidence in an embedded genome browser. The GUI uses the same analysis engine and outputs as the command line, connecting interactive exploration with scripted processing.

**Fig. 3.**
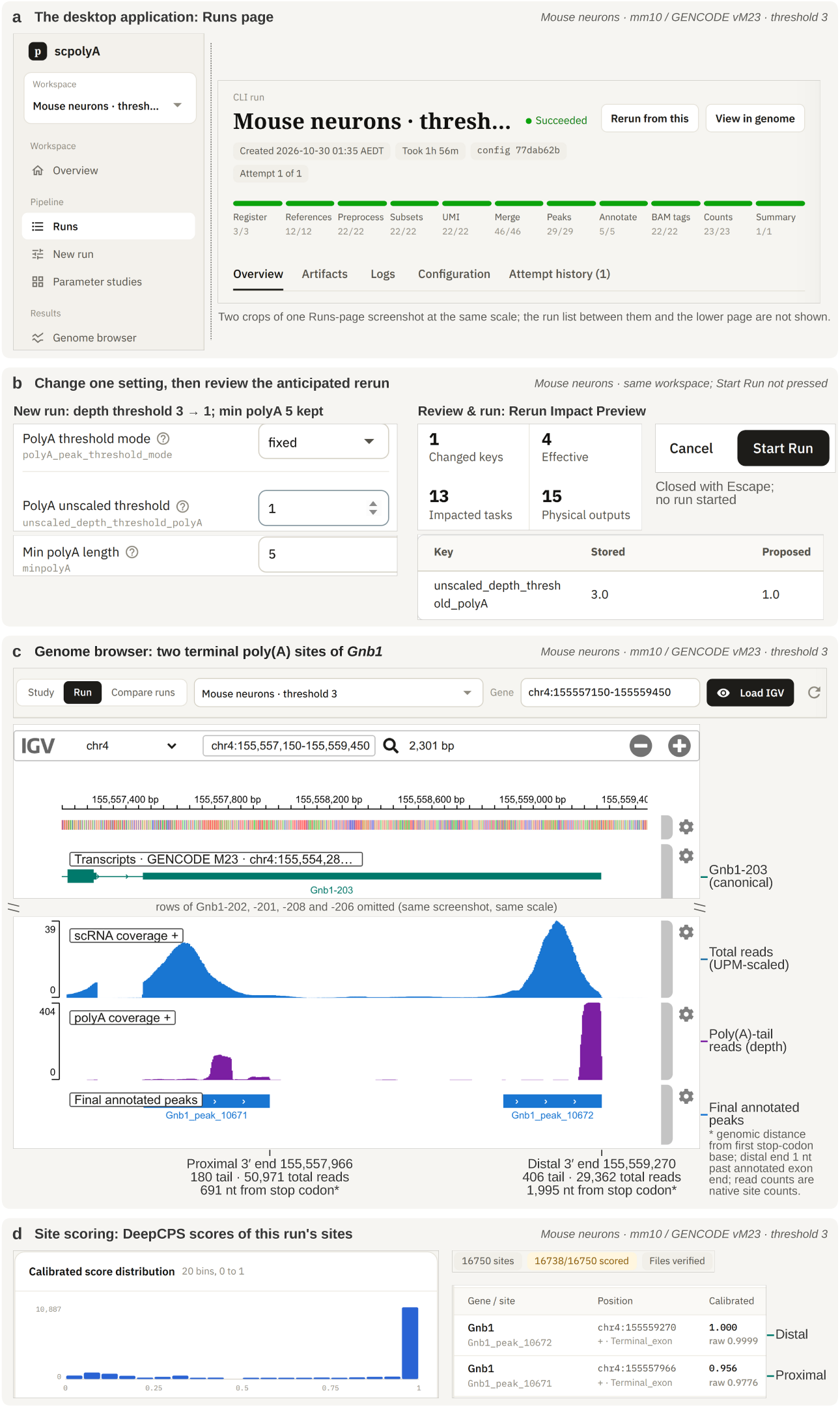
An integrated workspace connects APA results to the evidence and settings that produced them. A site catalogue is often the starting point for further decisions about threshold stringency and individual transcript ends. The scPolyA desktop interface allows these decisions to be explored within the same analysis environment, from an existing run to genomic inspection and DeepCPS assessment. **a,** The Runs page shows a completed threshold-3 analysis of mouse neurons, including execution stages and access to its outputs. The two crops are taken from the same screen. **b,** Lowering the tail-read depth threshold from 3 to 1 produces a preview of tasks and outputs that would be recomputed or reused. The panel shows a preview, not an executed rerun. **c,** At *Gnb1*, the embedded genome browser aligns two terminal poly(A) sites with the canonical transcript, total-read coverage and poly(A)-tail-read peaks. The proximal and distal endpoints are 691 and 1,995 nt downstream of the first stop-codon base and are supported by 180 and 406 tail reads, respectively. **d,** The same mouse-neuron run can be examined on the Site scores page. The proximal and distal *Gnb1* sites have calibrated atlas-proximity scores of 0.956 and 1.000, providing sequence-informed assessments alongside the read coverage.

After an initial analysis, users can revisit poly(A)-site discovery settings to examine how different thresholds affect site identification and downstream APA measurements. Before launching a rerun, the GUI previews which computational steps and outputs will change, allowing results from unaffected steps to be reused (Fig. 3b). Parameter studies support systematic comparisons across multiple settings, while side-by-side genome browsing enables inspection of differences at individual loci. Users can therefore evaluate thresholds by both the number of candidate sites recovered and the read evidence supporting them.

The GUI also integrates a genome browser that allows users to inspect identified poly(A) sites alongside their supporting reads and transcript annotations. In the mouse-neuron dataset, *Gnb1* illustrates how this enables direct examination of alternative RNA ends (Fig. 3c). Its proximal and distal sites lie 691 and 1,995 nt downstream of the stop codon, respectively. The two sites are supported by distinct poly(A)-tail-read peaks containing 180 and 406 reads, together with corresponding peaks in total-read coverage. Viewed alongside transcript annotations, these signals provide evidence for two alternative RNA ends within the same gene.

DeepCPS adds sequence-informed assessment of candidate sites to this workflow. Integrated into the scPolyA pipeline, its calibrated site scores are available through the GUI’s “Site scores” view (Fig. 3d), including for the two *Gnb1* sites illustrated in the genome browser. Users can therefore examine model assessments alongside poly(A)-tail-read support, total-read coverage and transcript annotations, connecting DeepCPS predictions to the underlying experimental evidence.

### Tail-anchored site maps recover molecular hallmarks and cell-type APA across protocols

We next evaluated scPolyA across sequencing protocols and compared its site discovery and APA measurements with seven established tools. The benchmark included eight single-cell datasets spanning four protocol families, two species and six tissues (Fig. 4a; Supplementary Fig. 2c). Five tools performed *de novo* site discovery (polyApipe, scAPAtrap, Sierra, SCAPTURE and SCAPE), while MAAPER and scUTRquant quantified reference-defined sites. Analyses used matched alignments and cell whitelists, and scPolyA was evaluated using read-based discovery without DeepCPS filtering (Methods).

**Fig. 4.**
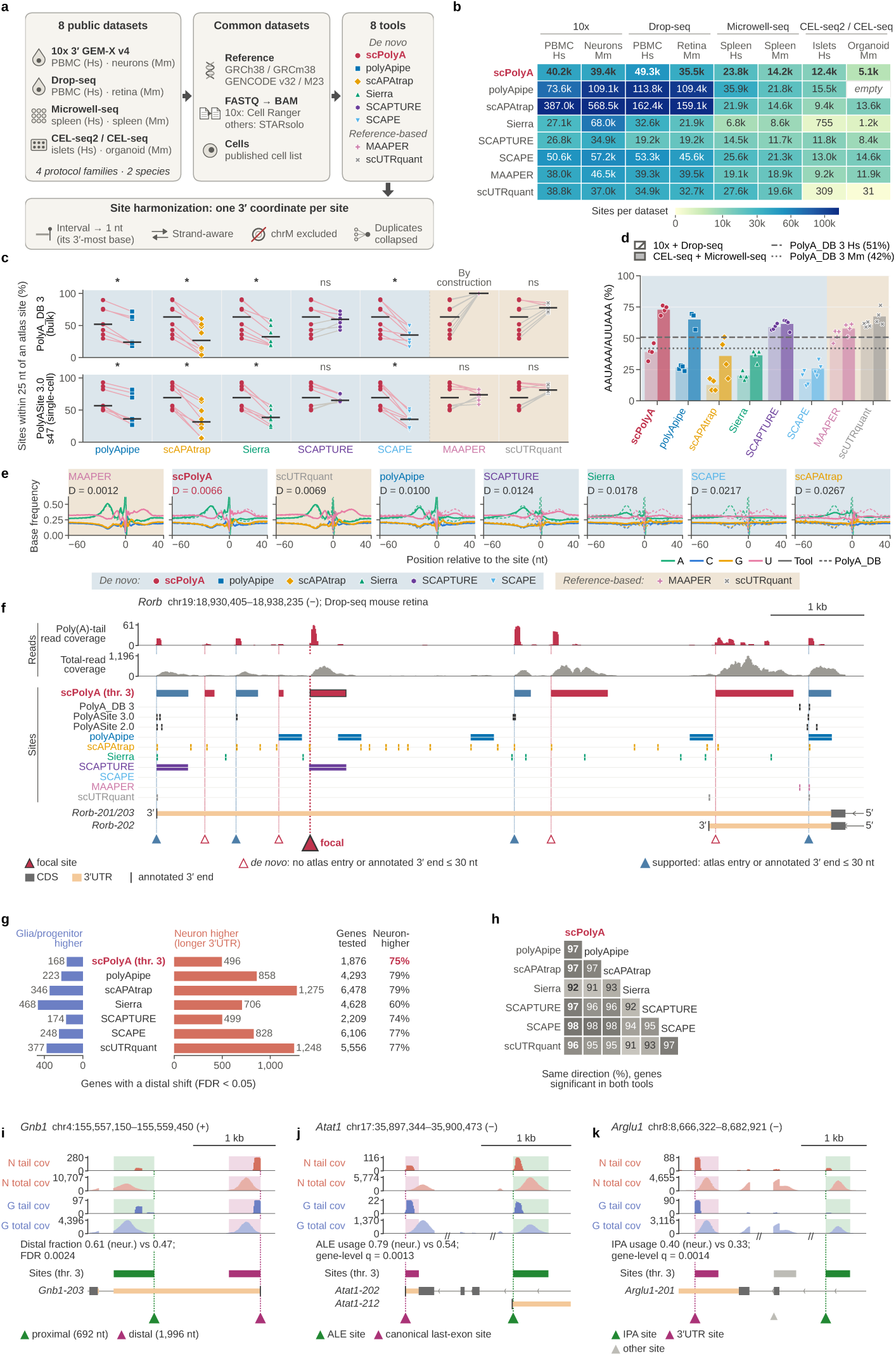
Read-supported poly(A)-site discovery recovers molecular features of transcript ends and reveals cell-type differences in APA. A site catalogue is biologically useful when its coordinates are supported by RNA-end evidence and its counts resolve alternative-end selection. Across four single-cell protocol families, scPolyA is assessed first against reference and sequence evidence, then through differences between neurons and glia or progenitors. **a,** Eight human and mouse datasets spanning 10x, Drop-seq, Microwell-seq and CEL-seq protocols were evaluated using matched inputs. Seven other methods cover *de novo* discovery and reference-guided quantification; reported sites were harmonised to strand-aware 3′ coordinates. **b,** Site counts vary considerably among methods and libraries, illustrating differences in catalogue breadth. Values are shown in thousands; “empty” denotes a run with no reported sites. **c,** The fraction of reported coordinates within 25 nt of PolyA_DB 3 or PolyASite 3.0 is compared within each dataset. scPolyA has median PolyA_DB 3 support of 63.3%, above four evaluated *de novo* methods and similar to DeepPASS-filtered SCAPTURE. Lines connect paired datasets, bars indicate medians and stars adjusted paired tests; MAAPER uses PolyA_DB coordinates by construction. **d,** Upstream AAUAAA or AUUAAA motifs provide an independent sequence-based view of the reported coordinates. Points represent datasets, bars the means and horizontal lines the human and mouse reference-atlas frequencies. **e,** Strand-oriented nucleotide frequencies from −60 to +40 nt around each reported site are compared with species-matched PolyA_DB reference profiles. Solid curves show each method and dashed curves the reference. The profile-distance metric *D* summarises positional differences in nucleotide frequencies, with smaller values indicating closer agreement. **f,** In mouse retina, a *Rorb* last-exon site is supported by 43 concentrated tail reads and an upstream AAUAAA motif despite being distant from atlas entries and annotated transcript ends. Coverage, transcript models and calls from other methods are aligned at the locus; the filled crimson triangle marks this candidate and blue triangles mark reference-supported sites. **g,** Having established site-level support, the analysis examines alternative terminal-site usage between neurons and glia or progenitors. Of 664 genes showing significant differences with scPolyA, 75% have higher distal-site usage in neurons (FDR < 0.05). Bars show the direction of each gene-level change. **h,** Methods agree on the direction of 91–98% of shifts among genes significant in both members of a pair. **i,** At *Gnb1*, distal tandem-site usage is higher in neurons than in glia or progenitors (0.61 versus 0.47), as shown alongside cell-class-specific coverage and the alternative terminal sites. **j,** At *Atat1*, an alternative last-exon site is used more strongly in neurons (0.79 versus 0.54), demonstrating an APA-associated change in transcript structure. **k,** At *Arglu1*, usage of an intronic site is higher in neurons (0.40 versus 0.33). In i–k, N and G denote neuron and glia/progenitor coverage; coloured shading and triangles identify the focal ends, and // denotes shortened introns.

Across the eight datasets, scPolyA identified a median of 29,651 poly(A) sites per dataset (range, 5,138–49,339), with an average of 51.7% in 3′ UTRs and 29.4% in introns (Fig. 4b; Supplementary Fig. 7a,c). We first compared the positional agreement of the *de novo* site catalogues with established poly(A)-site annotations. Using PolyA_DB 3 [43] as the reference, a median of 63.3% of scPolyA sites lay within 25 nt of a reference site, compared with 23.9% for polyApipe, 26.6% for scAPAtrap, 32.4% for Sierra and 35.0% for SCAPE (Fig. 4c; Supplementary Fig. 7b). scPolyA exceeded these four methods in every evaluable dataset (*P*_BH_ ≤ 0.025), while shifted-site controls showed only 2.9–3.7% agreement. Its reference-atlas agreement was also comparable to SCAPTURE, which incorporates DeepPASS-based sequence filtering (63.3% versus 59.4%). Evaluation against PolyASite 3.0 [46] showed a similar pattern, with median agreement of 69.5% for scPolyA and 65.1% for SCAPTURE, compared with 31.6–38.6% for the other *de novo* methods. Reference-guided methods require a different interpretation: scUTRquant achieved 77.6% agreement, while MAAPER achieved 100% agreement with PolyA_DB 3 by construction, as it uses this atlas to define the sites being quantified. The partial overlap of sites and multi-site genes between tools further showed that different discovery strategies recovered overlapping but distinct catalogues (Supplementary Figs 9 and 10a,b).

The number and reference-atlas agreement of detected sites also depend on the minimum read support required for site discovery. To examine this, we varied scPolyA’s tail-read threshold from three to one, which increased the number of detected sites by 1.4–4.5-fold (Supplementary Fig. 7a) and the number of genes with at least two sites by 1.7–10.4-fold (Supplementary Fig. 11a,d). Sites supported by fewer reads generally showed lower atlas agreement, which increased from 21.8% among single-read sites to 85.6% among sites supported by more than 30 tail reads (Source Data; Supplementary Table 1). However, the effect of lowering the threshold differed markedly across protocols. Atlas agreement decreased from 71.8% to 29.7% in 10x PBMC but only from 94.1% to 89.6% in mouse Microwell-seq spleen. At the one-read threshold, the two 10x datasets also showed higher fractions of downstream A-rich candidates (28.3% and 22.1%, versus at most 2.2% in the other datasets; Supplementary Fig. 11b). These findings highlight how the same read-support threshold can yield different site catalogues across sequencing protocols.

Beyond reference-atlas agreement, we examined whether scPolyA sites exhibited the characteristic sequence features of cleavage and polyadenylation. A median of 57.1% of scPolyA sites contained an upstream AAUAAA or AUUAAA polyadenylation signal (PAS), similar to SCAPTURE (60.7%) and higher than the 17.7–28.3% observed for the other *de novo* methods (Fig. 4d). We also compared nucleotide frequencies at positions surrounding the identified cleavage sites and found that scPolyA produced the sequence profile most closely resembling that of reference-atlas sites among the *de novo* methods (Fig. 4e). Further motif analysis revealed additional upstream PAS variants and UGUA elements, together with downstream U- and GU-rich sequences (Supplementary Fig. 8). These findings show that scPolyA recovers the characteristic sequence architecture of cleavage and polyadenylation sites, complementing their positional agreement with reference atlases.

An important advantage of *de novo* poly(A)-site discovery is the potential to identify RNA ends missing from existing reference catalogues. Across eight single-cell datasets, a median of 36.7% of scPolyA-identified sites lay more than 25 nt from PolyA_DB 3 annotations and 30.5% from PolyASite 3.0 annotations. One illustrative example is a site in the last exon of *Rorb* in Drop-seq mouse retina (Fig. 4f). Located 1,656 nt upstream of the annotated transcript end, this site could represent an alternative proximal polyadenylation event generating a shorter 3′ UTR. A prominent cluster of 43 poly(A)-tail reads within 25 nt of the site, together with local total-read coverage and a canonical AAUAAA polyadenylation signal 23 nt upstream, provides supporting evidence for this candidate RNA end.

We next tested whether the resulting site-resolved APA measurements captured differences in RNA-end choice between cell types. In 10x mouse-brain data, 664 of 1,876 tested genes differed in non-proximal terminal-site usage between neurons and glia or progenitors (false discovery rate (FDR) < 0.05). Of these, 75% favoured more distal sites in neurons (Fig. 4g), consistent with the established tendency towards longer neuronal 3′ UTRs [8]. All seven tools evaluated for this comparison recovered a neuron-higher majority (60–79%). Among genes significant in both tools of a pair, directional agreement was 91–98%, while an abundance-preserving null yielded no significant genes (Fig. 4h; Supplementary Fig. 10c). Thus, the same broad cell-type-associated pattern emerged across methods despite differences in their site catalogues.

Finally, we inspected individual loci to illustrate how cell-type differences in RNA-end usage reflect distinct forms of APA. In *Gnb1*, distal tandem-site usage rose from 0.47 in glia and progenitors to 0.61 in neurons (Fig. 4i). In *Atat1*, alternative-last-exon usage increased from 54% to 79% (Fig. 4j), while intronic-site usage in *Arglu1* increased from 33% to 40% at a site in canonical intron 2 (Fig. 4k). Together, these examples show how scPolyA links read-supported RNA-end maps to distinct forms of cell-type-associated APA, including tandem 3′ UTR choice, alternative last-exon use and intronic polyadenylation.

### scPolyA extends read-supported APA analysis across spatial transcriptomics platforms

The single-cell analyses established how scPolyA connects RNA-end discovery to differences in APA between cell populations. Spatial transcriptomics adds a further dimension by revealing where alternative RNA ends are used within tissues. Because scPolyA operates on barcoded alignments, its discovery and quantification strategy can also be applied when barcodes identify spots, bins or pixels. We evaluated this approach across eight tissue sections from seven spatial protocols: Visium, legacy Spatial Transcriptomics (ST), Visium HD 3′, Stereo-seq, Slide-seqV2, DBiT-seq and HDST (Fig. 5a; Supplementary Table 2).

**Fig. 5.**
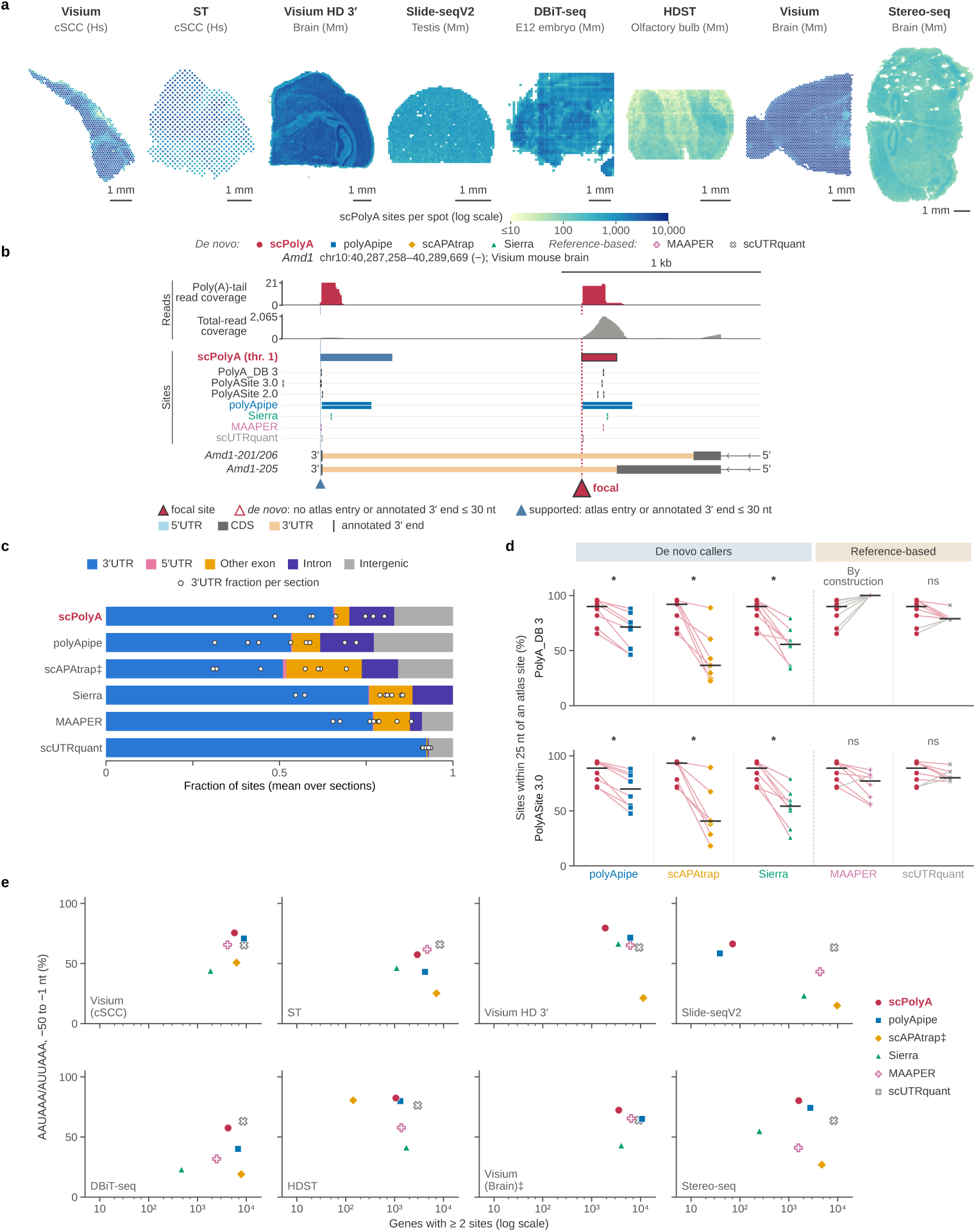
Cleavage-evidence-guided site discovery extends to spatial transcriptomics across different read structures and resolutions. Spatial barcodes change the unit in which RNA-end usage is measured, but the underlying distinction between tail-read evidence and total-read coverage remains. Across seven spatial platforms, scPolyA recovers site catalogues enriched for established RNA ends while retaining candidates outside existing annotations. **a,** Eight sections from Visium, legacy ST, Visium HD, Slide-seqV2, DBiT-seq, HDST and Stereo-seq are shown in their tissue geometries. Colour denotes the number of scPolyA sites detected with at least one UMI per spot, bin or pixel, on a common logarithmic scale. **b,** In mouse-brain Visium data, a last-exon site in *Amd1* lies outside nearby reference and annotated transcript ends but is supported by 18 tail reads and an upstream AUUAAA signal. The focal candidate is marked in crimson along-side coverage, atlas and transcript tracks. **c,** The reported sites span 3′ UTR, 5′ UTR, other exonic, intronic and intergenic contexts. Stacked bars show the mean genomic-context distribution across sections and points the 3′ UTR fraction in individual sections. **d,** Across spatial sections, scPolyA reaches median PolyA_DB 3 proximity of 89.9%, compared with 71.3% for polyApipe, 55.6% for Sierra and 36.5% for scAPAtrap. Lines pair methods within the same section, bars show medians and stars adjusted paired tests; MAAPER uses reference-defined coordinates by construction. PolyASite 3.0 provides the second atlas comparison. **e,** For each section and method, the number of genes with at least two reported sites is plotted against the fraction of sites carrying an upstream AAUAAA or AUUAAA signal, showing catalogue breadth and motif support together. Filled symbols denote *de novo* methods and open symbols reference-based methods. ‡ marks the uncompleted scAPAtrap brain-Visium run.

To illustrate how poly(A)-tail reads support *de novo* site discovery in spatial transcriptomes, we first examined mouse-brain Visium data. scPolyA identified a candidate poly(A) site in the *Amd1* 3′ UTR, 555 nt from the nearest annotated transcript end (Fig. 5b). The site was supported by 18 poly(A)-tail reads and an upstream AUUAAA polyadenylation signal, despite being absent from PolyA_DB 3 and PolyASite. We further assessed the performance of scPolyA across all eight spatial sections, comparing site recovery and reference-atlas agreement with other *de novo* methods. On average, 65.5% of scPolyA sites were located in 3′ UTRs (Fig. 5c). Median PolyA_DB 3 agreement reached 89.9%, compared with 71.3% for polyApipe, 55.6% for Sierra and 36.5% for scAPAtrap (Fig. 5d). scPolyA exceeded each of these *de novo* methods in every section where that method completed its analysis (*P*_BH_ ≤ 0.024), while shifted-site controls showed at most 4.7% agreement. The comparison of multi-site gene recovery and upstream AAUAAA or AUUAAA polyadenylation signals further highlighted differences in catalogue breadth and sequence support between methods (Fig. 5e).

Slide-seqV2 provided a demanding test because recognisable poly(A)-tail reads accounted for only approximately 0.03% of its alignments, compared with 0.47–5.6% in the other spatial sections. Even with such sparse direct cleavage evidence, scPolyA identified 1,853 sites, compared with 1,217 for polyApipe. Sierra and scAPAtrap reported much larger catalogues of 7,751 and 59,423 candidates, respectively, but with lower reference-atlas agreement. These results demonstrate that scPolyA can recover a focused catalogue enriched for reference-supported poly(A) sites even when detectable tail reads are rare, providing a foundation for quantifying RNA-end usage across spatial locations.

### Efficient processing and parameter-aware reuse support iterative APA analysis

We next evaluated the computational cost of scPolyA, examining both the performance of an initial analysis and the savings achieved through parameter-aware rerunning with different poly(A)-site discovery thresholds. We therefore evaluated both the cost of an initial scPolyA run and how much computation could be reused after a threshold change. Across eight single-cell datasets, six *de novo* tools were benchmarked on the same server under a 16-CPU quota and a 256-GiB memory limit (Methods). scPolyA had the second-shortest median wall time and the lowest median peak memory (Fig. 6a,b). Its median method-stage runtime was 25 min (range, 5.7–136 min; 17.1 s per million aligned reads), with median peak memory of 34.2 GiB (range, 11.0–103.2 GiB).

**Fig. 6.**
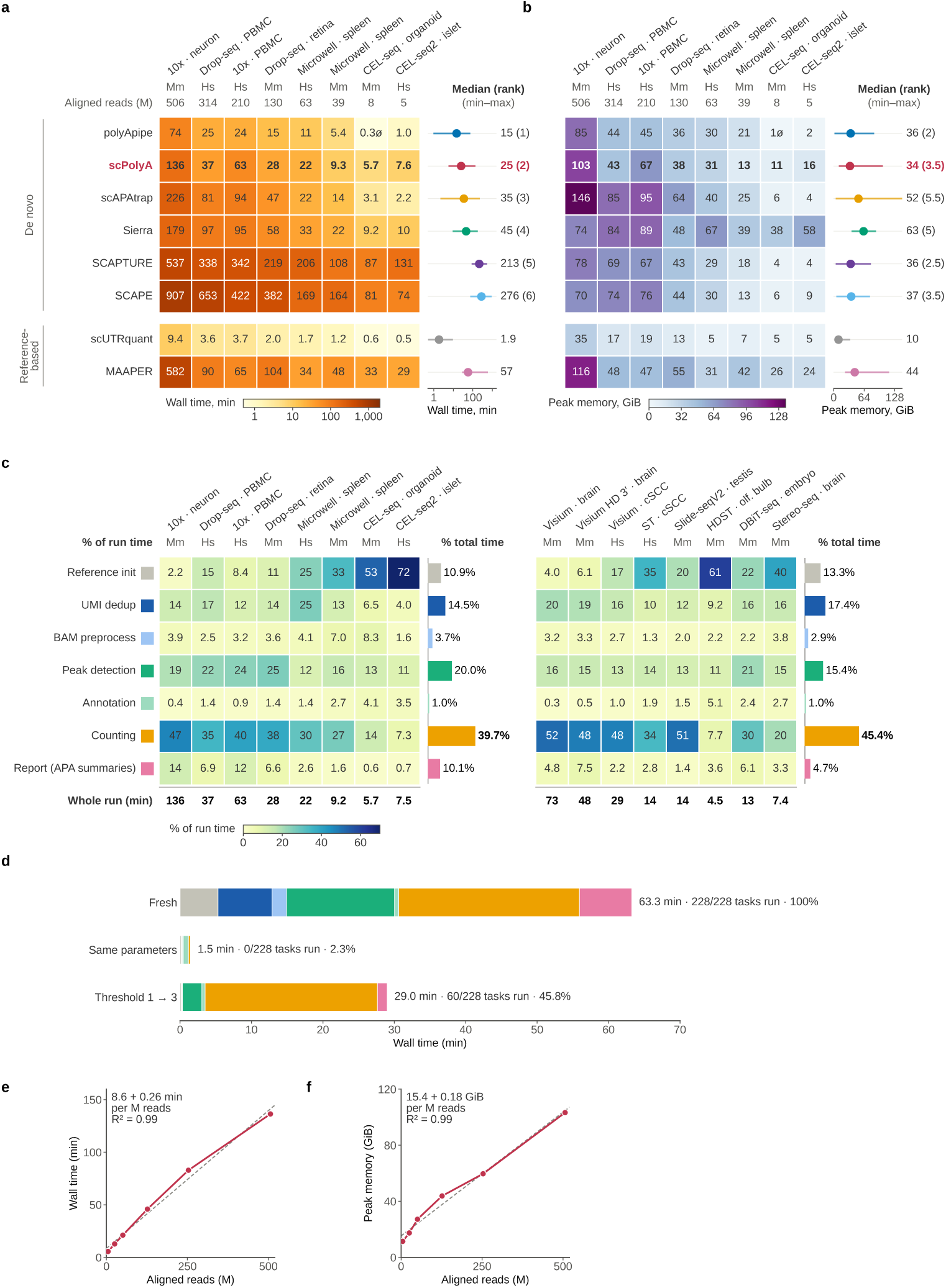
Selective recomputation makes evidence thresholds practical to revisit. Changing a site-selection threshold can alter the candidate catalogue and every downstream APA measurement, while much of the initial read preparation remains applicable. The figure assesses initial computational cost and then shows how scPolyA updates the affected work without repeating an entire analysis. **a,** Native method-stage wall time is compared across eight single-cell libraries ordered by aligned-read count. scPolyA has a median duration of 25 min; heat-map colours use a logarithmic scale. **b,** Peak memory for the same runs has a median of 34.2 GiB for scPolyA and includes page cache. Side summaries in a and b show medians, ranges and median within-dataset ranks among the six *de novo* methods; reference-based methods are displayed but not ranked with them. **c,** Stage-level profiles across eight single-cell and eight spatial analyses show that UMI counting takes the largest pooled fraction of scPolyA runtime (39.7% for single-cell libraries). This work depends on the selected site intervals and must be updated when assignments change. **d,** In 10x PBMC, an initial tail-threshold-1 analysis requires 63.3 min and executes 228 tasks, whereas an unchanged rerun completes in 1.5 min without executing tasks. Raising the threshold to 3 executes 60 tasks in 29.0 min, including updated molecule counting and APA summaries. Colours identify stage groups and percentages are relative to the initial run. **e,** On downsampled 10x mouse-neuron data, scPolyA runtime increases approximately linearly with aligned-read count (*R*^2^ = 0.99); the dashed line shows the fit. **f,** Peak memory also increases approximately linearly over the same depth series (*R*^2^ = 0.99). The analyses used a 16-CPU quota; peak memory includes page cache.

We next examined where computation was spent and which steps needed to be repeated when site-selection settings changed. UMI counting accounted for 39.7% of pooled runtime, peak detection for 20.0%, and reference preparation, deduplication and read preprocessing together for 29.1% (Fig. 6c). Overall, the stages through count-matrix generation accounted for 89.9% of runtime. scPolyA’s parameter-aware task graph can reuse upstream preparation while updating the site-dependent computations needed to generate revised UMI counts and APA measurements.

We measured the benefit of this reuse by revisiting a completed 10x PBMC analysis. A fresh analysis at a tail-read threshold of one executed 228 tasks in 63.3 min. Repeating the same configuration required no tasks to be rerun and completed in 1.5 min. Raising the threshold from one to three executed only 60 of the 228 tasks and completed in 29.0 min, or 45.8% of the fresh-run time (Fig. 6d; Supplementary Table 4). Upstream preparation was reused, and peak-detection time fell from 15.1 to 2.7 min. Site-specific UMI counting was recomputed for the revised catalogue (24.1 min, compared with 25.3 min in the fresh run), ensuring that the updated APA measurements reflected the new molecule assignments. The time saving therefore came from retaining valid computations while repeating the steps affected by the threshold change.

Finally, we examined how the cost of an initial analysis scaled with sequencing depth by downsampling the 10x mouse-neuron dataset. As the number of aligned reads increased from approximately 5 million to 506 million, runtime rose from 5.6 to 136 min, closely following a linear relationship (*R*^2^ = 0.99). Peak memory increased from 11 to 103 GiB over the same range (Fig. 6e,f). Together, efficient initial processing and selective recomputation allow scPolyA users to explore poly(A)-site discovery thresholds without repeating the entire pipeline for each setting.

### Patient-matched and spatial data reveal keratinocyte-associated 3′UTR shortening in cSCC

Increased proximal poly(A)-site usage has been associated with proliferation and transformation [6, 7]. In heterogeneous tumours, however, interpreting 3′ UTR shortening requires distinguishing changes within cell populations from differences in the cellular composition of tissue regions. We therefore applied scPolyA to patient-matched cutaneous squamous cell carcinoma (cSCC), asking whether 3′ UTR shortening recurs across patients, which cell populations exhibit it and how it is reflected in tumour tissue. We analysed 47,061 single cells from tumour and matched normal skin of ten patients, together with Visium or legacy Spatial Transcriptomics (ST) sections from six of these patients [47] (Fig. 7a; Supplementary Fig. 12).

**Fig. 7.**
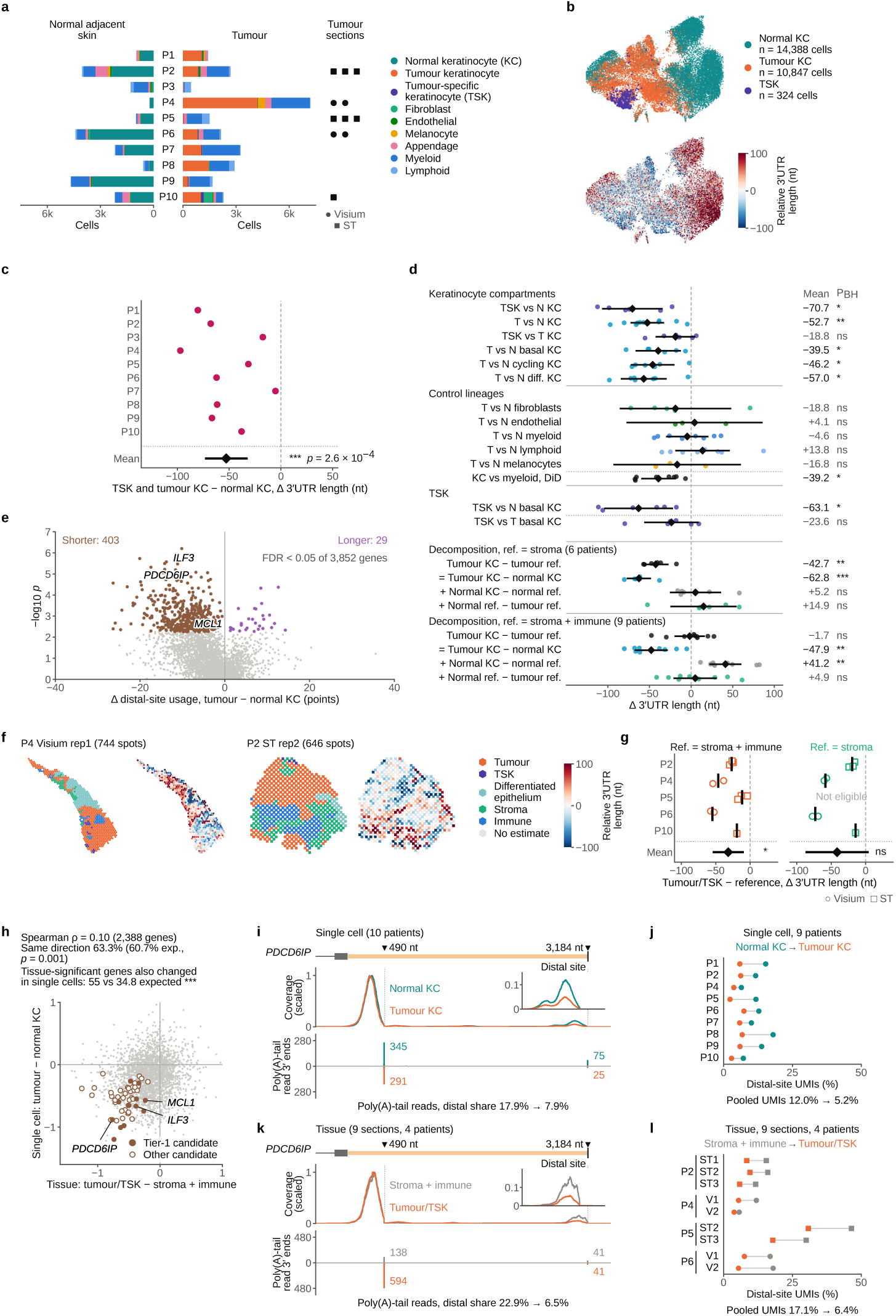
Patient-matched and spatial measurements identify a recurrent keratinocyte-associated pattern of 3′ UTR shortening in cutaneous squamous cell carcinoma (cSCC). A tumour-region APA difference can reflect altered RNA-end choice within cells, changes in cellular composition or both. The analysis first establishes the patient- and lineage-level pattern, then examines its spatial context and the individual ends contributing to it. **a,** The study combines normal-skin and tumour single-cell data from ten patients, with their cell-type composition and available Visium or legacy spatial sections indicated. **b,** In a Harmony-corrected representation of keratinocytes, normal, tumour and tumour-specific keratinocyte (TSK) populations are shown alongside cell-level relative 3′ UTR length. The reported contrasts are calculated at the patient level. **c,** Tumour-associated keratinocytes have shorter 3′ UTRs in all ten patients, with a mean patient-paired difference of −52.8 nt (*p* = 2.6 × 10*^−^*^4^). Points represent patients and the diamond the mean and 95% confidence interval. **d,** The shortening extends across basal, cycling and differentiating keratinocytes, with no consistent corresponding change in the other lineages examined. The lower rows use an analogous single-cell contrast to separate the tumour-keratinocyte state effect from cellular-composition and reference-state contributions. Points represent patients; diamonds show means with 95% intervals, and adjusted *P* values are indicated. **e,** Of 3,852 eligible genes, 432 differ in distal-site usage between tumour and normal keratinocytes (FDR < 0.05), with 403 favouring more proximal ends in tumour cells. Each point represents a gene, positioned by effect size and significance. **f,** Spatial sections display marker-defined tumour, TSK-enriched, differentiated-epithelial, stromal and immune regions beside the relative 3′ UTR length of individual spots. **g,** Tumour/TSK-enriched regions also show shorter 3′ UTRs than stromal-plus-immune regions (11 sections from five patients; mean of patient means −31.8 nt). Stroma-only comparisons provide a second reference. Symbols distinguish Visium and legacy ST sections; inference uses patients as replicates after averaging sections within patient. **h,** Gene-level effects show modest overall concordance between spatial and single-cell analyses (Spearman *ρ* = 0.10). Among 121 genes meeting the spatial section-level significance threshold, 55 also change in the same direction in single cells at nominal *p <* 0.05, exceeding the permutation expectation. **i,** Read coverage and tail-read endpoints around two terminal sites of *PDCD6IP* are shown for pooled normal and tumour keratinocytes. The sites correspond to approximately 490- and 3,184-nt 3′ UTRs and provide direct read-end context for the usage comparison. **j,** Distal-site UMI usage at *PDCD6IP* decreases from 12.0% in pooled normal keratinocytes to 5.2% in pooled tumour keratinocytes, with the same direction in nine evaluable patients. **k,** The corresponding spatial-region tracks show the proximal and distal *PDCD6IP* sites in stroma-plus-immune and tumour/TSK-enriched locations. The distal share of tail-read endpoints decreases from 22.9% to 6.5%. **l,** Distal-site UMI usage decreases from 17.1% to 6.4% in pooled spatial counts, with the same direction across nine evaluable sections. The distal tail-read fraction in panel k is a separate read-based measurement from the UMI-based usage shown in panels j and l.

We first compared usage-weighted 3′ UTR lengths between tumour and normal keratinocytes within each patient. Among 25,559 keratinocytes (Fig. 7b), tumour keratinocytes had shorter 3′ UTRs in all ten patients, with a mean paired difference of −52.8 nt (95% CI −73.5 to −32.2 nt; two-sided paired *t*-test *p* = 2.6 × 10^−4^; Fig. 7c). The direction persisted after controlling for gene composition and sequencing depth (Supplementary Fig. 13a). In contrast, intronic-site usage showed no consistent directional change across patients (lower in five of ten; *p* = 0.47). The most consistent tumour-associated APA change therefore involved terminal 3′ UTR choice.

Was this shortening confined to a particular keratinocyte state, or did it extend across the lineage? Basal, cycling and differentiating keratinocytes each showed shorter 3′ UTRs in tumour than in matched normal skin, with mean differences of −39.5 to −57.0 nt and the same direction in 6/6, 6/6 and 7/7 eligible patients, respectively (*P*_BH_ ≤ 0.027; Fig. 7d). Tumour-specific keratinocytes (TSK) also had shorter 3′ UTRs than normal keratinocytes (−70.7 nt; 6/6 patients; *P*_BH_ = 0.015), although their difference from other tumour keratinocytes was smaller and not significant (−18.8 nt; 4/6 patients; *P*_BH_ = 0.17). Myeloid, lymphoid, fibroblast, endothelial and melanocyte populations showed no corresponding consistent shift (*P*_BH_ ≥ 0.49). Among eight patients eligible for a direct comparison, the tumour–normal change in keratinocytes was 39.2 nt more negative than in myeloid cells (*P*_BH_ = 0.013). Thus, shortening extended across keratinocyte states and was most consistently associated with this lineage.

To identify the RNA-end choices underlying the length difference, we examined distal-site usage gene by gene. Of 3,852 genes tested in tumour and normal keratinocytes, 432 differed at FDR < 0.05, and 403 of these shifted towards proximal sites (Fig. 7e). In 50 within-patient cell-label permutations, at most one gene passed the same significance threshold. These results connect the recurrent reduction in 3′ UTR length to widespread changes in terminal-site usage across individual genes.

We next asked whether the keratinocyte-associated pattern could also be detected in tumour-enriched tissue regions. Expression-marker scores were used to identify tumour/TSK-enriched, stromal and immune regions in the spatial sections (Fig. 7f). Across 11 eligible sections from five patients, tumour/TSK-enriched regions had shorter 3′ UTRs than stromal and immune regions in every patient (mean of patient means −31.8 nt; 95% CI −54.5 to −9.1 nt; *p* = 0.018; Fig. 7g). The direction was also negative when stroma alone was used as the reference (−41.5 nt across four patients) and persisted in analyses restricted to shared genes and adjusted for sequencing depth (Supplementary Fig. 14). In a separate assessment of spatial organisation, relative 3′ UTR length showed structure beyond an abundance-preserving null in 15 of 16 sections, with 12 remaining significant after adjusting for depth and region identity (Supplementary Fig. 14b; all section maps in Supplementary Fig. 15). The tissue-level pattern was consistent with the single-cell keratinocyte findings, although spatial contrasts also depend on the cellular composition of the regions being compared.

To examine the contribution of cellular composition, we decomposed analogous contrasts in the single-cell data into tumour-state, cellular-composition and reference-state terms (Fig. 7d; Methods). With stroma as the reference, the total difference was −42.7 nt across six patients, comprising a −62.8-nt tumour-state term, a +5.2-nt composition term and a +14.9-nt reference-state term. This was close to the spatial stroma-only estimate of −41.5 nt. Including immune cells in the reference increased the composition term to +41.2 nt, offsetting much of the −47.9-nt tumour-state term and yielding a total of −1.7 nt across nine patients. The corresponding spatial estimate was −31.8 nt, but the single-cell and spatial contrasts involved different samples and cellular mixtures. Across reference definitions and common-gene analyses, the tumour-state term remained negative (Supplementary Fig. 13c). These analyses show why the magnitude of a tissue-level APA contrast depends on its reference populations, even when a keratinocyte-associated shortening component is recurrent.

The single-cell and spatial analyses pointed to related shifts in RNA-end usage, but did the same genes change in both? Across 2,388 jointly assessed genes, effect sizes were modestly correlated (Spearman *ρ* = 0.10; 95% bootstrap interval 0.06–0.14). Directional agreement was 63.3%, compared with 60.7% expected from the marginal distributions (permutation *p* = 0.001; Fig. 7h). Agreement was more apparent among the 121 genes meeting the spatial section-level significance threshold: 55 also changed in the same direction in single cells at nominal *p <* 0.05, compared with 34.8 ± 4.8 expected under permutation (*p* = 5 × 10^−5^). Thirteen additionally passed the single-cell FDR criterion and the specified composition and myeloid screens (Supplementary Fig. 16; Supplementary Table 5). This identified a focused set of genes with APA changes supported by both cellular and tissue-level measurements.

*PDCD6IP* illustrates this cross-modality pattern at an individual locus. The gene encodes ALIX, an ESCRT-associated protein involved in cytokinetic abscission and exosome biogenesis [48, 49], and its proximal and distal poly(A) sites define 3′ UTRs of 490 and 3,184 nt (Fig. 7i,k). In single-cell data, distal-site usage was lower in tumour keratinocytes in all nine evaluable patients, with pooled usage decreasing from 12.0% in normal keratinocytes to 5.2% in tumour keratinocytes (Fig. 7j). The same direction was observed in all nine evaluable spatial sections from four patients: distal usage decreased from 17.1% in stromal/immune regions to 6.4% in tumour/TSK-enriched regions (Fig. 7l). The fraction of spatial poly(A)-tail reads at the distal site likewise decreased from 22.9% to 6.5% (Fig. 7k), providing direct read-end evidence alongside the UMI-based measurements. This recurrent preference for the shorter *PDCD6IP* 3′ UTR nominates the gene for functional follow-up and illustrates how scPolyA connects patient-level APA changes to cell populations, tissue regions and individual transcript ends.

### Agent-guided analyses connect research questions to the evidence needed for APA interpretation

To support question-driven APA analysis, we developed an APA-specialised, domain-guided AI assistant integrated with the scPolyA pipeline (Fig. 1g). The assistant translates biological research questions into executable analysis workflows, verifies the resulting outputs and interprets findings using the underlying poly(A)-site evidence and APA measurements. We evaluated these capabilities through three newly executed case studies (Fig. 8a), progressing from generating APA measurements in human peripheral blood mononuclear cells (PBMCs), to examining how site-discovery thresholds affect the results, and finally to assessing candidate RNA ends absent from reference atlases in mouse neurons. Each question was investigated by two agents using the same source data. One execution per case is shown in Fig. 8, with all six runs presented in Supplementary Fig. 17.

**Fig. 8.**
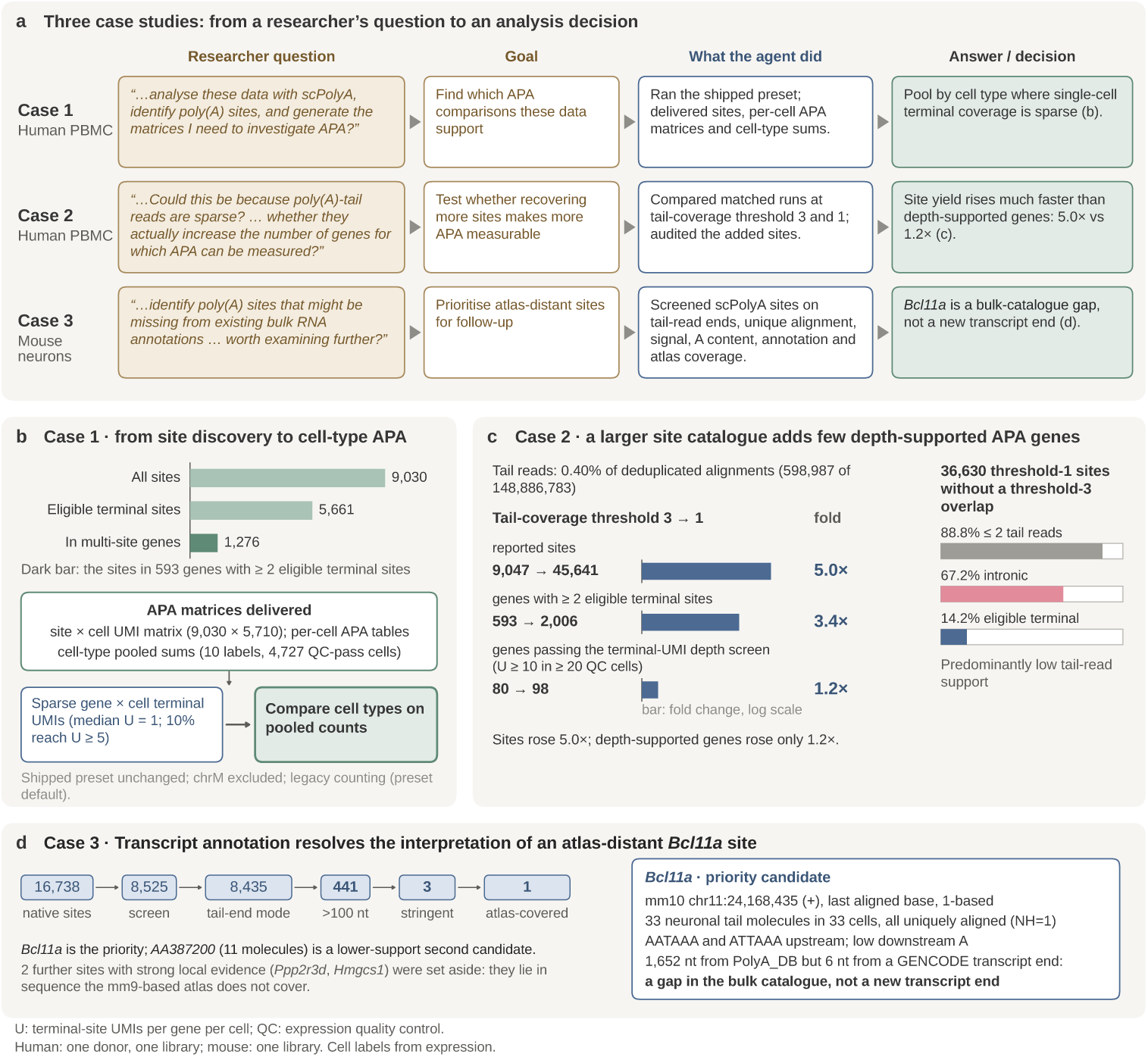
APA-specialised AI agents execute question-driven scPolyA analyses and assess the supporting evidence. Three researcher questions examine whether the agents can generate APA measurements, compare discovery settings against molecular coverage, and interpret atlas-distant RNA ends in their genomic context. The cases progress from single-cell quantification to threshold sensitivity and candidate review. **a,** Each question leads to an analysis plan, scPolyA execution, output review and an evidence-based interpretation. Two agents independently addressed each question using the same source data. One execution per case is shown here, with all six in Supplementary Fig. 17. **b,** The first PBMC analysis yields 9,030 sites across 5,710 barcodes. Of 5,661 eligible terminal sites, 1,276 belong to 593 genes with at least two terminal ends. Sparse terminal UMI coverage in individual gene–cell observations (median one; 10% with at least five) motivates pooled counts for ten groups comprising 4,727 quality-controlled cells (nine named immune-cell groups and one unassigned group). **c,** Lowering the pooled tail-read depth threshold from 3 to 1 increases sites from 9,047 to 45,641 (5.0-fold) and multi-site genes from 593 to 2,006 (3.4-fold), while genes meeting the terminal-UMI depth screen rise from 80 to 98 (1.2-fold). The screen requires at least ten terminal UMIs in at least 20 quality-controlled cells. Among 36,630 threshold-1 candidates without overlap with a threshold-3 site, 88.8% have no more than two tail reads, 67.2% are intronic and 14.2% have eligible terminal context. Case 2 used a separate native execution from the 9,030-site Case 1 analysis in panel b. **d,** The mouse-neuron analysis narrows 16,738 native sites to 8,525 after the initial evidence screen, 8,435 with an observed tail-end mode and 441 more than 100 nt from PolyA_DB. Three sites meet the stringent evidence criteria; only *Bcl11a* lies in atlas-covered sequence. Its RNA end is supported by 33 tail-read molecules in 33 neurons and lies 1,652 nt from a PolyA_DB entry but 6 nt from a GENCODE transcript end, identifying a gap in the bulk poly(A)-site catalogue at an annotated transcript end.

The first question was whether a routine PBMC library could provide the measurements needed to investigate APA across immune-cell populations. The agent executed scPolyA using its predefined configuration and delivered an annotated catalogue of 9,030 sites and a site-by-cell unique molecular identifier (UMI) matrix covering 5,710 cell barcodes (Fig. 8b). Among 5,661 eligible terminal sites, 1,276 belonged to 593 genes with at least two alternative terminal ends. These genes offered a starting point for comparing RNA-end choice, but molecular coverage in individual cells was sparse: among observed gene–cell terminal-APA measurements, the median was one terminal UMI and only 10% had at least five. The agent therefore also produced site-specific UMI sums for nine annotated immune-cell groups and an unassigned group, comprising 4,727 quality-controlled cells. The analysis connected the size of the discovered catalogue to the cellular resolution at which its alternative ends could be meaningfully compared.

This led to a second question: would a more permissive discovery threshold produce a comparable increase in measurable APA? Poly(A)-tail reads comprised 0.40% of deduplicated alignments in the PBMC library. In a separate execution from Case 1, the agent compared matched scPolyA runs at pooled tail-read depth thresholds of 3 and 1, increasing reported sites from 9,047 to 45,641 (5.0-fold) and genes with at least two eligible terminal sites from 593 to 2,006 (3.4-fold; Fig. 8c). The gain was smaller when measured by cell-level molecular coverage. With the agent’s screen requiring at least ten terminal UMIs in at least 20 quality-controlled cells, the number of qualifying multi-site genes increased from 80 to 98 (1.2-fold). Of 36,630 threshold-1 candidates without overlap with a threshold-3 site, 88.8% had at most two supporting tail reads, 67.2% were intronic and 14.2% had eligible terminal context. These findings distinguish the substantial expansion of candidate RNA ends from the more modest gain in genes with sufficient terminal-UMI coverage for subsequent APA analysis.

The third case shifted from measurement to interpretation: which RNA ends absent from the bulk poly(A)-site atlas merit closer investigation? In one mouse-neuron library, the agent screened 16,738 native scPolyA sites using cell-associated read support, endpoint localisation, sequence and alignment evidence, and reference-atlas coverage (Fig. 8d). The successive initial and endpoint-mode screens retained 8,525 and 8,435 sites, respectively; 441 were more than 100 nt from the nearest eligible PolyA_DB entry. Three passed the stringent evidence criteria, but only one lay in sequence represented by the mm9-derived atlas. This candidate, in *Bcl11a*, was supported by 33 tail-read molecules from 33 neurons, with uniquely aligned reads, upstream polyadenylation signals and little downstream A-rich sequence. Although its endpoint lay 1,652 nt from the nearest PolyA_DB site, it was only 6 nt from an annotated GENCODE transcript end. The analysis therefore identified a gap in the bulk site catalogue at an already annotated RNA end. The two other strongly supported candidates, in *Ppp2r3d* and *Hmgcs1*, lay outside the atlas’s mapped sequence coverage and were retained as distinct reference-coverage cases.

Across these cases, the assistant connected researcher questions to scPolyA execution and then to evidence-based analytical decisions. It identified when sparse individual-cell measurements favoured cell-type pooling, showed how threshold stringency changes the relationship between candidate discovery and molecular coverage, and interpreted an atlas-distant site using its supporting reads and transcript annotation. Together, the examples illustrate how the framework can support decisions about what RNA-end measurements reveal and which findings warrant further examination.

## Discussion

The value of a poly(A)-site map lies in the regulatory questions it makes possible: which RNA ends different cells use, how those choices change with cell state, and where the changes appear in tissue. Earlier studies have shown that conventional single-cell transcriptomes can reveal poly(A) sites and their usage. An important insight from our study is that these are not independent steps. How a site is defined shapes which molecules are assigned to it, which alternative ends can be compared and, ultimately, which APA differences become measurable. scPolyA connects atlas-independent site discovery, DeepCPS-based assessment, barcode-resolved APA measurements and iterative evidence review in a unified framework for investigating RNA-end choice across cells and tissues.

The connection begins with two kinds of reads that describe RNA ends differently. Reads carrying an untemplated poly(A) tail are relatively rare but provide comparatively precise positional evidence, whereas abundant neighbouring reads provide the depth needed to measure usage. polyApipe also uses tail reads for discovery, assigning molecules within a configurable upstream window (250 nt by default) [29]. In scPolyA, tail-read evidence anchors candidate ends and total-read coverage defines variable-width intervals for UMI counting. These intervals follow the observed read distribution around each candidate, accommodating differences in coverage between loci and library chemistries. Because an interval determines which UMIs are counted for each alternative end, the definition of a poly(A) site also shapes how its usage is measured. This connection between locating an end and counting its molecules is central to quantitative APA analysis.

Finding additional RNA ends, however, does not necessarily mean measuring more APA. A gene may have several candidate sites in a pooled catalogue while most individual gene–cell observations contain too few terminal UMIs to estimate their relative use. This explains why relaxing tail-read thresholds can greatly expand site catalogues and the number of multi-site genes while yielding a much smaller increase in genes with sufficient molecular coverage for downstream comparisons. The relevant sensitivity depends on the scale of the biological question. A site can be well supported across a library even when its usage becomes informative only after combining cells from a defined population. The same site-by-cell matrix can support both individual-cell inspection and cell-group comparisons, provided the available molecular depth is taken into account. Site yield, evidence for a candidate end and the precision of its usage measurement should therefore be considered together.

Reference-atlas agreement offers another form of evidence, but its interpretation depends on what the atlas contains. Reference catalogues reflect the tissues and experimental settings in which RNA ends have been sampled, so an unmatched candidate is not necessarily an unsupported one. The *Rorb* and *Amd1* examples illustrate how concentrated tail reads and recognisable polyadenylation signals can support candidate ends absent from these catalogues. Sequence-based site recognition and cleavage prediction have important precedents [28, 39]. DeepCPS brings these capabilities to candidates generated through read-based discovery, using sequence and coverage for site scoring and sequence to refine cleavage positions. The greater cross-dataset detection of high-scoring single-read candidates, together with improved localisation under sparse support, shows why tail-read count alone does not capture all available evidence. At the same time, atlas proximity, a score calibrated to atlas support, a predicted cleavage position and observed UMI usage answer different questions. Interpreting them together, while preserving these distinctions, can prioritise sites for further investigation without treating a model prediction as proof of a biological cleavage event.

The cSCC analysis illustrates the biological perspective gained when RNA-end measurements are linked to cell identity. Shorter 3′ UTRs have long been associated with proliferation and transformation [6, 7], but heterogeneous tumours contain populations whose relative abundance can differ between samples. In patient-matched tumour and normal skin, keratinocytes showed shorter 3′ UTRs in all ten patients. The pattern extended across basal, cycling and differentiating states, while other examined lineages showed no corresponding consistent shift. Its presence across these keratinocyte states suggests that the finding cannot be explained solely by changes in the proportion of cycling cells. Terminal-site changes across hundreds of genes further connected the aggregate shortening to particular RNA-end choices. The resulting insight is more specific than an overall tumour–normal difference: it identifies a recurrent, keratinocyte-associated change within a heterogeneous tissue.

Spatial measurements add an important dimension to this interpretation. A tumourenriched region contains mixtures of cells, so its difference from stromal or immune regions reflects both RNA-end use within populations and the cellular composition of the regions being compared. In the cSCC data, changing the reference population substantially altered the aggregate contrast, even though the tumour-keratinocyte-associated component remained negative in analogous single-cell decompositions. The magnitude of a regional APA difference therefore cannot be read directly as the magnitude of a change within one cell population. Single-cell measurements help explain what contributes to a spatial contrast, while spatial maps reveal where the combined signal occurs. Gene-level concordance between modalities was modest overall, but a subset of changes recurred in both, including the proximal shift in *PDCD6IP*. These convergent loci offer focused starting points for investigating the regulatory consequences of altered 3′ UTR choice in tumours.

The dependence of these interpretations on analytical choices makes iterative analysis part of the scientific process. A fixed tail-read threshold does not represent the same strength of evidence across libraries, because chemistry, coverage and downstream A-rich candidates vary. Changing that threshold can alter both the candidate site map and which APA comparisons have enough molecules to be informative. scPolyA’s parameter-aware execution makes these choices practical to revisit by reusing valid upstream computations, while its GUI brings the resulting site maps, read coverage, transcript annotations and DeepCPS scores into a common workspace. The APA-specialised AI assistant extends this approach from interactive inspection to question-driven execution and evidence review. In its case studies, it recognised when sparse cell-level counts favoured cell-group summaries, distinguished an expansion of candidate sites from a gain in measurable APA, and interpreted atlas-distant candidates in the context of sequence and reference coverage. The value of this integrated environment lies in keeping the biological question, analytical decisions and supporting molecular evidence connected as an analysis develops.

This creates an opportunity for conventional cell and tissue atlases. Gene abundance and transcript-end choice are different dimensions of RNA regulation, and a stable gene count need not imply stable use of its alternative ends. Measuring both from the same libraries can reveal cell populations or tissue regions in which transcript-end composition changes, potentially altering the regulatory sequences retained in RNA without a corresponding change in total gene expression. Such observations nominate specific post-transcriptional mechanisms for experimental investigation. By linking read-supported end positions to UMI-based usage and biological context, scPolyA makes this layer of regulation accessible in routinely generated single-cell and spatial transcriptomes, without requiring a poly(A)-site reference atlas as input or a dedicated new sequencing assay. Its broader contribution is to bring RNA-end choice into the quantitative analysis of cells and tissues.

## Methods

### scPolyA pipeline

scPolyA converts barcoded, unique molecular identifier (UMI)-tagged 3′-end alignments into strand-specific poly(A)-site intervals, site-by-cell or site-by-spot UMI counts, and within-gene APA measurements (Fig. 1). It combines two complementary sources of evidence: reads with untemplated poly(A) tails locate candidate RNA ends, and coverage from all deduplicated reads defines the intervals over which those ends can be quantified. Candidate sites are discovered from reads pooled across a cohort, while molecule counting retains the original cell or spatial-barcode identities. No poly(A)-site reference atlas is required for discovery. The main single-cell, spatial and cutaneous squamous cell carcinoma (cSCC) analyses used the `terminal-aware` mode with the merged-site representation; the separately described agent Case 1 used a shipped `legacy` preset. Supplementary Methods detail alternative analysis modes, command-line options, file formats and software settings (Supplementary Tables 3 and 6).

### Input preparation and genomic annotation

Inputs comprise coordinate-sorted BAM alignments, a sample table, a gene annotation in GTF format and chromosome sizes. Barcodes and UMIs are read from configurable BAM tags (`CB` and `UB` in the evaluated datasets). Cell selection, barcode correction and alignment are completed upstream. Samples are processed by chromosome, with UMI-tools used to remove PCR duplicates within barcode, strand and alignment-position groups. Its directional-network algorithm joins UMIs separated by one nucleotide according to their relative read counts. Libraries previously deduplicated may be explicitly identified as such. Sample-specific barcode suffixes preserve the identity of cells or spatial units across input libraries.

The annotation supplies gene, exon and coding-sequence intervals, canonical-transcript contexts and stop-codon-relative positions. For a protein-coding gene, scPolyA selects the transcript with the largest summed coding-sequence length to establish the canonical terminal exon and the reference stop coordinate for measuring 3′ UTR distance. The evaluated reference construction excluded genes whose names begin with `Gm` or contain `Rik`. The selection rules for overlapping genes, transcripts without a coding sequence, optional bulk-informed annotations and nonstandard contigs are described in Supplementary Methods. The same preparation steps are used by the command line and graphical application.

### Poly(A)-tail extraction and coverage-constrained site construction

Candidate tail reads are identified by an A-rich soft clip at the aligned 3′ end of a forward-strand read or a T-rich clip at the corresponding end of a reverse-strand read. The evaluated analyses required at least five clipped A/T nucleotides, no more than one other clipped nucleotide and mapping quality at least 10 for tail evidence. These thresholds describe the observed soft clip and do not estimate biological poly(A)-tail length. Strand assignment assumes that the alignment orientation corresponds to transcript sense, as in the evaluated forward-stranded 3′-tag protocols. The mapping-quality requirement applies specifically to tail-read extraction; total-read coverage and UMI counting retain their separately defined criteria.

scPolyA generates separate, strand-specific coverage tracks from all deduplicated reads and from the extracted tail reads, pooling alignments across barcodes. Total-read coverage intervals that overlap annotated coding sequences are removed before calling total-read regions; the tail-read track is not coding-sequence-masked. The study used total-read coverage scaled per million alignments and a gene-specific detection threshold equal to 5% of the gene’s maximum coverage. Tail-read coverage remained unscaled, with fixed inclusive depth thresholds of one or three deduplicated tail reads per genomic position, as specified for each analysis. Thus “threshold 1” and “threshold 3” refer to pooled discovery evidence, not to cell-level UMI counts. Above-threshold regions were joined across gaps of at most 10 nt in each track.

A total-read region becomes eligible for site construction when it overlaps a tail-read region on the same strand. The 3′ boundaries of the tail-read regions establish candidate endpoints, partitioning the supported total-read region into potential counting intervals (Fig. 1b,c). Their opposite boundaries follow total-read coverage; coverage extending beyond the outermost tail-supported endpoint is not included. Two interval representations are available. The `resolved` representation retains the individual split intervals, while `merged` joins nearby intervals on the same strand. The evaluated merged mode (10-nt merge distance) therefore yields one final site per contiguous total-read region or merged group, anchored at its most 3′ tail-supported boundary. Closely spaced ends within one continuous total-read region are consequently not separately quantified in these analyses. The strand-specific half-open coordinate rules and support aggregation are provided in Supplementary Methods.

### Internal-priming assessment, site annotation and UMI counting

To identify candidates potentially created by oligo(dT) priming within A-rich genomic sequence, scPolyA examines the genome-aligned sequence immediately adjacent to each tail read’s endpoint. A read is flagged when at least eight of the ten strand-oriented aligned bases are A (or T on the reverse strand). For site *i*, the internal-priming score *m_i_*is the fraction of overlapping tail reads flagged by this rule; soft-clipped tail sequence does not enter the calculation. Sites are assigned to same-strand genes and classified by the canonical transcript, prioritising terminal exon, other exon, intron and the region up to 2,000 nt downstream of the canonical terminal exon (Supplementary Fig. 1a). In `terminal-aware` mode, non-intronic sites with *m_i_ >* 0.9 were removed before counting. Intronic candidates were counted, but only those with *m_i_ ≤* 0.7 were eligible for intronic APA summaries. These cutoffs preserve the distinction between recording an intronic candidate and accepting it as an intronic APA measurement.

Deduplicated total-read alignments are assigned to the final site intervals using sense-strand featureCounts with a minimum 1-nt overlap. Reads assigned ambiguously to multiple sites or carrying multimapping assignments do not contribute to the site counts. A subsequent directional UMI correction groups assigned molecules by barcode and site, across read positions. The resulting sparse count matrix contains *n_ic_*, the number of distinct UMIs assigned to site *i* in cell or spatial unit *c*. Counting uses the final site intervals, so changing the discovery threshold or merging strategy may change the molecules attributed to an end. Site annotations, count matrices and downstream summaries are generated as native scPolyA outputs (Supplementary Table 6).

### Terminal and intronic APA measurements

Let *T_g_* contain the eligible terminal-exon and downstream sites of gene *g*, and *J_g_* its eligible intronic sites. For barcode *c*, terminal and intronic UMI support are 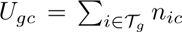 and 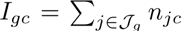. For genes with at least two eligible terminal sites and sufficient terminal support (*U_gc_ ≥* 1 in the evaluated configuration), scPolyA reports terminal-site usage and usage-weighted 3′ UTR distance:

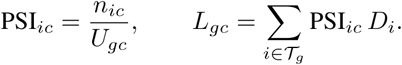

Here *D_i_* is the nonnegative genomic distance, in transcript orientation, from the inferred canonical stop position to the site endpoint. This quantity is an unspliced genomic distance and is not a reconstructed transcript length. Signed distances are separately used to identify the most distal terminal site; distal flags are provided in the native annotation for downstream usage analyses.

Site-level and gene-level intronic polyadenylation indices measure intronic versus terminal molecule support using a pseudocount:

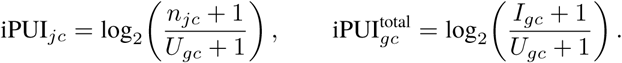

Terminal support for iPUI includes all retained terminal sites, including those of single-terminal-site genes. Native tables also support gene-level terminal counts, distal-terminal usage and gene- or site-resolved comparisons across barcodes. An unobserved or ineligible gene–barcode pair is distinguished from an eligible site with zero observed usage; missing APA values are not converted to zero. The detailed eligibility, sparse-output and coordinate conventions, together with the alternative `legacy` calculation, are documented in Supplementary Methods (Supplementary Fig. 1c).

### Differential APA and statistical analysis

Differential APA tests in this study were performed using study-specific analysis scripts applied to the native site-count matrices and annotations. The separate released `scpolya-diff-apa` edgeR/limma module [50, 51] was not used for the reported biological tests. Unpaired cell-type comparisons used site-UMI ratios pooled within the groups and studentised cell-label permutation tests, retaining fixed eligibility criteria across permutations. Patient-matched analyses pooled counts by patient and condition and tested within-patient differences in transformed distal usage or gene-centred 3′ UTR distance. Benjamini–Hochberg correction was applied within each specified family of gene-level tests or contrasts. The relevant sample definitions, permutation counts, eligibility criteria and sensitivity analyses are specified below and in Supplementary Methods.

### Graphical user interface and parameter-aware execution

#### Shared execution engine and selective reanalysis

scPolyA provides command-line, Python and NiceGUI-based desktop interfaces to the same configuration parser, execution engine, task-state database and native output schema. Computation is organised into tasks whose identities depend on stage, input unit and declared parameters; the resulting state records support reuse of completed work. When the pooled tail-read depth threshold changes, upstream preprocessing, deduplication and coverage generation can be retained while tail-region selection, site construction, annotation, counting and APA summaries are updated (Fig. 1f). The GUI’s “Review & run” view and the Python API report anticipated reuse and recomputation before a rerun begins. Reuse is based on recorded task identities and stage-specific completion checks. Because BAM contents, input registrations and some annotation or barcode-tag changes are not comprehensively fingerprinted, those changes require a new run identifier or output directory. This restriction was respected in the reported analyses; task hashing, output checks and path-repair controls are specified in Supplementary Methods.

#### Analysis configuration, genomic inspection and DeepCPS scores

The graphical workspace brings together run configuration, execution status, parameter studies and comparison of native outputs with an embedded igv.js genome browser [52] (Fig. 3). Genome tracks display annotated transcripts, total-read coverage, poly(A)-tail evidence and alternative site intervals, allowing analysis settings to be assessed at individual loci and across parameter choices. The mouse-neuron *Gnb1* example in Fig. 3c shows two terminal sites with their respective endpoint and coverage evidence. The integrated DeepCPS “Site scores” view displays model assessments of discovered candidates in the same run, alongside the underlying coverage and site context (Fig. 3d). This functionality is an implemented part of the pipeline and GUI; users may inspect model scores without applying them as site filters.

#### GUI execution and numerical concordance

Figure 3a,b illustrates a completed threshold-3 mouse-neuron run and a preview of the work that a threshold change would affect; the previewed change was not executed for that figure. A separate experiment initiated an actual threshold-3-to-1 rerun through the GUI on the 10x Genomics GEM-X 3′ v4 human PBMC library (5,710 barcodes; Supplementary Fig. 18). An independent command-line run at the same threshold-1 settings provided a numerical check of the completed GUI execution. Rows were matched by primary key across eight native output tables—sites, site–gene annotations, site UMIs, terminal gene–cell counts, PSI, weighted 3′ UTR distance and site- and gene-level iPUI—with identical categorical values and missingness and absolute-plus-relative numerical tolerance of 10^−12^. All rows agreed (Supplementary Fig. 19). This separates the interface’s ability to preview a rerun from evidence that it executed one and produced the same analytical outputs as the command line.

#### DeepCPS architecture, training and extended evaluation

DeepCPS comprises two neural networks addressing complementary properties of RNA ends (Fig. 2): a candidate-scoring model estimates proximity to established poly(A) sites from sequence and local coverage, and a localisation model predicts the positional distribution of cleavage-associated read endpoints from sequence. The networks are integrated into scPolyA for site assessment and localisation. DeepCPS filtering was *not* used for the main read-based benchmarking or cSCC catalogues; the historical analyses therefore assess the discovery pipeline without an additional neural-network site-selection step.

#### Candidate labels and chromosome-partitioned datasets

We assembled 217,102 scPolyA threshold-1 candidates, before internal-priming filtering, from four droplet datasets: human 10x PBMC, mouse 10x neurons, human Drop-seq PBMC and mouse Drop-seq retina (Supplementary Fig. 3a). Each strand-oriented candidate endpoint defined the centre of a 201-nt sequence window. For candidate scoring, positives lay within 25 nt of a same-strand site in PolyA_DB 3 or PolyASite 2.0 [43, 44]; negatives lay more than 100 nt from either atlas or had no reference match. Candidates 26–100 nt from reference sites were excluded from the labelled scoring task. These labels describe atlas proximity, not independent biochemical confirmation of cleavage.

To limit genomic-position overlap across model development and evaluation, chromosomes 2, 9 and 16 were held out for testing in both species; chromosomes 5 and 12 were reserved for validation, hyperparameter selection and calibration. The remaining chromosomes were used for training. The scoring test partition contained 33,336 labelled candidates (13,440 positive and 19,896 negative). The chromosome partitions were shared by both Deep-CPS tasks, the ablation experiments and baselines fitted in this study. Released DeepPASS and DeepPASTA models were evaluated on the same held-out candidate sets without retraining.

#### Network architectures and training

The scoring model encodes the transcript-oriented 201-nt window as four one-hot nucleotide channels and one strand-specific total-read-coverage channel. Coverage is measured in alignments per million and divided by the maximum within the window, retaining local coverage shape while removing the absolute-depth scale. An 11-nt convolution with 64 output channels feeds six residual dilated convolutional blocks (kernel width 5; dilations 1, 2, 4, 8, 16 and 1; dropout 0.2). Global mean and maximum pooling are combined before a 128-unit hidden layer and binary output logit. The model was trained with class-weighted binary cross-entropy and AdamW (learning rate 10^−3^, weight decay 10^−2^, batch size 128), up to 20 epochs with early stopping based on validation area under the precision–recall curve (AUPRC). Predictions from three independently trained seeds were averaged. Isotonic regression on validation chromosomes calibrated the ensemble’s predicted atlas-proximity probabilities; operating-point filters were selected from the uncalibrated scores.

The separate localisation model uses one-hot sequence only, with the same convolutional stem and residual trunk followed by a 1 × 1 convolution yielding 201 positional logits. Its training targets were counts of deduplicated same-strand poly(A)-tail-read endpoints at reference-proximal sites with at least ten supporting reads. Training minimised multinomial negative log-likelihood of the observed endpoint distribution, with endpoint counts capped at 200 reads per site and random window shifts from −40 to +40 nt. AdamW optimisation used learning rate 10^−3^, weight decay 0.1 and batch size 128, with at most 20 epochs and early stopping on validation loss. Three model seeds were averaged, and the modal position of their predicted distribution was reported as the localisation coordinate. The observed read-end mode provides a practical proxy for cleavage position; it is not an assay-independent measurement of the biochemical cleavage nucleotide.

#### Held-out performance and evaluation against other methods

On held-out chromosomes, candidate scoring was measured by area under the receiver operating characteristic and precision–recall curves (AUROC and AUPRC), with 95% intervals from 1,000 class-stratified bootstrap resamples. DeepCPS was evaluated on the same candidates as the released DeepPASS and DeepPASTA models [28, 37], rankings by tail-read count or internal-priming score and a fitted logistic-regression sequence baseline using upstream PAS presence and downstream A-richness. Paired scoring comparisons used 2,000 bootstrap resamples clustered by the 5,656 species–gene loci, with two-sided empirical *P* values and Benjamini–Hochberg adjustment. Candidate subsets were also stratified by tail-read support to assess model behaviour when a site had only one or a few supporting reads.

Localisation accuracy was the fraction of atlas-supported test sites with at least ten poly(A)-tail reads for which the predicted coordinate equalled, or lay within *k* nucleotides of, the observed modal endpoint. Evaluation included the original scPolyA site boundary, a canonical PAS-offset rule and APARENT v1 [36], with anchoring selected on validation data. Locus-level bootstrap resampling provided uncertainty intervals, and paired exact-coordinate success rates were compared by exact McNemar tests. Scoring against atlas entries, matching a modal read endpoint and quantifying actual site UMIs were treated as distinct outcomes.

#### Input and embedding ablations

To determine which information supported candidate scoring, matched models used the same chromosome splits, architecture, training procedure and seeds while varying input features. Ablations included sequence alone, total-read coverage alone, the combined sequence-and-coverage input, and frozen Nucleotide Transformer v2-50m embeddings [45] projected to 32 dimensions with or without coverage. Additional models used frozen DeepPASS or DeepPASTA embeddings followed by a trained prediction head. Separate cross-species and cross-protocol experiments trained on one species or droplet-protocol group and evaluated on another. These analyses distinguish the contributions of sequence and observed coverage and test whether pretrained sequence representations improve the same candidate-scoring task. Exact sequence-window adaptation for the released models and the baseline feature definitions are provided in Supplementary Methods.

#### In silico mutagenesis

We measured model sensitivity to local sequence by substituting each position in 2,000 positive and 1,000 negative held-out candidate windows with each of the three alternative nucleotides, leaving the coverage channel unchanged. For nucleotide *j*, its contribution was the prediction on the original sequence minus the mean prediction across its three substitutions:

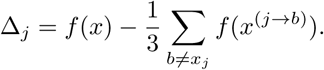

For candidate scoring, *f* was the pre-sigmoid logit; for localisation, it was the log probability assigned to the observed modal read-end position. A positive contribution means that changing the original base lowers the relevant model output. The resulting profiles were used to examine upstream polyadenylation signals, downstream A-rich sequence and nucleotide preferences around cleavage-associated endpoints (Fig. 2f). These computational perturbations characterise learned sequence associations; they do not measure experimental cleavage efficiencies.

#### Protocol transfer, catalogue selection and independent PBMC replication

To test transfer beyond the original droplet datasets, frozen DeepCPS ensembles were applied without retraining or threshold tuning to held-out chromosomes from Microwell-seq, CEL-seq and CEL-seq2 datasets. Candidate scoring was assessed on threshold-1 sites after the native internal-priming filter; localisation was evaluated separately on reference-proximal sites with at least ten read endpoints (Fig. 2g). Score-based site-selection thresholds were chosen on validation chromosomes of the four training datasets. One uncalibrated threshold (*τ* = 0.707) matched the number of threshold-1 candidates overlapping threshold-3 sites; another (*τ* = 0.292) matched their PolyA_DB 3 support. Both were fixed before application to the eight held-out dataset partitions. Candidate retention, atlas proximity, genomic context and read-thinning effects on distal usage were evaluated separately, with matched-dataset tests and adjustment families described in Supplementary Methods.

We further assessed cross-dataset reproducibility using independently generated 10x and Drop-seq PBMC datasets, with observed poly(A) sites serving as the evaluation outcome instead of reference-atlas annotations (Fig. 2h–k). Although both datasets contributed training examples on other chromosomes, this evaluation was restricted to held-out test chromosomes. Methods ranked the same threshold-1 candidates, and a candidate was considered reproduced if the other PBMC dataset contained a same-strand threshold-1 site within 25 nt. Matched catalogue sizes and tail-read-support strata, including single-read sites, tested whether scores distinguished candidates with the same observed tail-read count. A separate sparse-site localisation test used the modal endpoint of at least ten deduplicated tail reads at a nearby site in the independent PBMC dataset as a read-based positional reference. The original site boundary, a candidate’s own read-end mode, the PAS rule and APARENT were evaluated against that same reference. These independent analyses examine reproducibility and position while retaining the distinction between a read-derived reference and a directly validated cleavage event.

### Datasets and comparative evaluation

The main single-cell benchmark analysed eight public 3′-tag datasets from four protocol families (10x Genomics, Drop-seq, Microwell-seq and CEL-seq/CEL-seq2), spanning two species and six tissues. The spatial benchmark comprised eight sections representing seven platforms: Visium, legacy Spatial Transcriptomics (ST), Visium HD 3′, Stereo-seq, Slide-seqV2, DBiT-seq and HDST. Corrected-barcode alignments, input libraries, read structures, accession identifiers and preprocessing differences are detailed in Supplementary Fig. 2 and Supplementary Table 2. The main comparisons used scPolyA’s terminal-aware, merged-site, read-based discovery with DeepCPS filtering disabled.

The method panel comprised five other *de novo* callers (polyApipe, scAPAtrap, Sierra, SCAPTURE and SCAPE) and two reference-guided quantification methods (MAAPER and scUTRquant). Comparisons used matched inputs and cell selections where supported, common gene annotations and strand-aware harmonisation of reported site endpoints. Evaluated implementations and settings appear in Supplementary Table 3 and Supplementary Methods. The methods differ in their supported inputs and locus definitions; notably, MAAPER derives positions from a reference atlas, SCAPTURE applies a separate DeepPASS sequence filter, and polyApipe uses fixed upstream counting windows (250 nt by default). A method was evaluated only where its native analysis completed; incomplete or empty outputs were documented.

Site reference agreement was the fraction of reported, strand-specific endpoints within 25 nt of a same-strand PolyA_DB 3 entry; shifted-reference controls and PolyASite 2.0/3.0 provided additional context. We compared genomic-context distributions, canonical polyadenylation signals, strand-oriented base composition, overlaps between callers and the number of genes with multiple reported sites (Figs 4 and 5; Supplementary Figs 7–11). Atlas proximity was treated as evidence of reference agreement, not a direct measure of cleavage accuracy. For atlas-distant examples, candidates were further assessed for concentrated tail-read endpoints, upstream PAS motifs, downstream A-richness, transcript annotation and tissue representation in the reference catalogues. Comparisons across completed datasets used paired Wilcoxon signed-rank tests with the stated Benjamini–Hochberg adjustment families; locus and atlas-audit rules are provided in Supplementary Methods.

For the mouse-neuron analysis (Fig. 4g–k), expression-based assignments defined neurons (6,315) and glial or progenitor cells (3,184). Differential non-proximal terminal-site usage was tested separately for each method on its own site-by-cell count matrix (threshold 3 for scPolyA), in genes with at least 20 terminal UMIs and ten expressing cells in each class. Each gene was tested with 2,000 studentised cell-label permutations, and false discovery rate was controlled at 0.05 within each method. Neuron- and glia-specific site-usage profiles and a gene-abundance-preserving null supported the cell-type comparison. Illustrative locus analyses distinguished tandem 3′ UTR, alternative last-exon (ALE) and intronic polyadenylation (IPA) choices using canonical transcript context (Supplementary Fig. 1b). These examples were selected for positional and read-level evidence; the *Arglu1* locus met the annotation criterion but fell below two otherwise applied illustrative support thresholds, as specified in Supplementary Methods.

Runtime and peak memory were evaluated serially on a shared server under a 16-CPU quota and 256-GiB cgroup memory limit for the single-cell comparison (Fig. 6a,b). Memory denotes the cgroup peak, including page cache. Timed stage boundaries differ between tools, so the reported comparative figures describe the stated native workloads, not identical operations (Supplementary Table 3). Parameter-reuse measurements compared a fresh threshold-1 PBMC analysis, an unchanged rerun and a threshold-1-to-3 rerun, each with the completed count matrix and APA summaries (Fig. 6d; Supplementary Table 4). Scaling was evaluated by subsampling the 10x mouse-neuron reads to five fractions plus the complete dataset and fitting runtime and memory against retained read number (Fig. 6e,f). Full resource accounting and timing definitions appear in Supplementary Methods.

### Patient-matched and spatial cSCC analyses

The cutaneous squamous cell carcinoma (cSCC) case study used public matched tumour and normal skin single-cell data from ten patients (GSE144236, 47,061 retained cells) and 16 Visium or legacy ST sections from six overlapping patients (GSE144239/GSE144240) [47]. Published cell identities and keratinocyte subtype labels were retained; no APA-derived cell identities were introduced. Three cohorts (single-cell, Visium and ST) were processed separately with scPolyA’s terminal-aware, merged mode and pooled tail-read threshold 3, without DeepCPS site filtering. The tissue compartments were defined using expression-marker scores as tumour/TSK-enriched, differentiated epithelium, stroma or immune regions; they represent enriched regions, not purified cell types (Supplementary Figs 12–15). Input processing, region scoring and inclusion criteria are specified in Supplementary Methods.

The primary length readout was a gene-centred, terminal-UMI-weighted genomic 3′ UTR distance. For barcode or pseudobulk unit *x*,

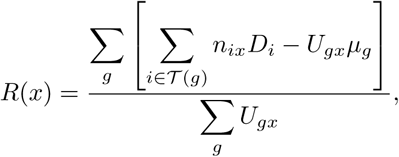

where *T* (*g*) denotes eligible terminal sites of gene *g*, *U_gx_* their total UMI count, *D_i_* their genomic stop-relative distances and *µ_g_*the chosen gene-specific reference mean. Genes required at least two terminal sites. The gene-specific reference mean was calculated separately at the cohort, spatial-section or matched-patient level, depending on the analysis. This gene-centering reduces the influence of differences in the genes represented across comparisons. Distal-usage and intronic-usage contrasts were evaluated separately; the latter uses an intronic fraction and is distinct from the native log-ratio iPUI output. Per-cell and per-spot visualisations required at least 20 eligible terminal UMIs per unit.

Patient-matched tests pooled eligible UMIs within patient, condition and cell class. Each contrast required at least ten cells per group, and patient-specific tumour-minus-normal differences were tested with two-sided paired *t*-tests and 95% confidence intervals. The primary keratinocyte comparison involved ten patients; subtype and control-lineage contrasts used their separately eligible patients and were adjusted within the reported families. Gene-level distal-usage tests required at least four evaluable patients with ten terminal UMIs per condition; transformed patient-level usage differences were tested by two-sided one-sample *t*-tests with Benjamini–Hochberg correction. Fixed-gene weighting, depth thinning, matched-cell analyses and within-patient label permutations assessed robustness (Supplementary Fig. 13).

For spatial comparisons, eligible terminal counts were pooled within marker-defined regions of each section, requiring at least ten spots per compared region. The primary tumour/TSK-enriched versus stroma-plus-immune contrast involved 11 sections from five patients; the stroma-only comparison included eight sections from four patients. Region contrasts averaged sections within patients and used patients as replicates; gene-level spatial tests used sections and are exploratory. Spatial structure was assessed using Moran’s *I* with abundance-preserving and depth/region-adjusted nulls (Supplementary Fig. 14). To examine the contribution of cellular composition to spatial contrasts, we decomposed the analogous single-cell contrast as

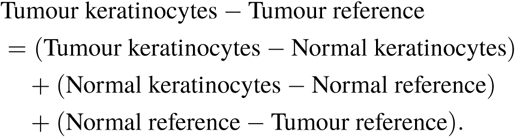

The decomposition was evaluated at the patient level with stromal or stromal-plus-immune references and common-gene sensitivity analyses (Supplementary Fig. 13c). Cross-modal agreement used genes evaluable in both data types, Spearman correlation, sign-agreement permutations and staged gene-level criteria, with candidate-selection and myeloid/composition controls detailed in Supplementary Methods and Supplementary Fig. 16.

### APA-specialised AI agents and case studies

Two APA-specialised agents, Claude Opus 5.5 in Claude Desktop and GPT-6-Astra in the GPT desktop application, each received the same three researcher questions and source datasets, producing six recorded executions (Fig. 8; Supplementary Fig. 17). Versioned scPolyA-specific skills and documentation supplied the available operations and evidence-review procedures, separately from the questions. The agents could inspect inputs and native outputs, execute scPolyA through its command line and interpret resulting site annotations, UMI counts and genomic evidence. Both used medium reasoning effort and scPolyA v2.1.0 in the recorded runs. Figure 8 presents the Claude execution for Case 1 and the GPT executions for Cases 2 and 3. Each displayed execution was selected after independent numerical review; detailed settings and the paired agent findings appear in Supplementary Methods and Supplementary Fig. 17. The case studies used one human PBMC donor library for the two PBMC questions and one mouse-neuron library for the neuronal question; their native catalogues and analysis criteria are reported separately.

#### Researcher questions

The following questions were submitted individually, separately from scPolyA skills, dataset paths and operational instructions.

**Case 1.** I have 10x single-cell RNA-seq data from human PBMCs. I’m interested in studying alternative polyadenylation across immune cells. Can you analyse these data with scPolyA, identify poly(A) sites, and generate the matrices I need to investigate APA?

**Case 2.** I want to investigate alternative 3′ UTR usage in immune cells, but my scPolyA results contain relatively few genes with multiple poly(A) sites. Could this be because poly(A)-tail reads are sparse? Can you investigate whether we can recover more sites, assess their reliability, and determine whether they actually increase the number of genes for which APA can be measured?

**Case 3.** I’m studying RNA regulation in neurons and want to identify poly(A) sites that might be missing from existing bulk RNA annotations. Can you investigate our single-cell neuronal data and find candidate sites with convincing evidence that would be worth examining further?

#### Case 1: PBMC matrix generation

The displayed agent used the shipped predefined PBMC configuration without modification. This preset excluded chrM and used `legacy` counting, unlike the terminal-aware mode in the core benchmarks. It produced 9,030 native sites (8,974 entering counting), a matrix of 5,710 barcodes and site-specific UMI totals by cell group. Post-run terminal eligibility assessment yielded 5,661 terminal sites, including 1,276 in 593 multi-terminal genes. Among observed gene–cell terminal measurements the median terminal-UMI count was one and 10% had at least five. The output also included pooled site UMI counts for ten groups: nine named immune-cell groups comprising 4,620 quality-controlled cells and 107 unassigned cells, for 4,727 QC-passing cells in total. The eligibility analysis of this native legacy run did not turn it into a terminal-aware pipeline execution.

#### Case 2: discovery thresholds and terminal-molecule depth

With matched settings within this case, the agent compared pooled tail-read depth thresholds 3 and 1 in the PBMC library. Native site counts increased from 9,047 to 45,641 and genes with at least two eligible terminal sites from 593 to 2,006. A separate terminal-UMI depth screen required at least ten total eligible terminal UMIs in each of at least 20 quality-controlled cells of a gene, identifying 80 and 98 multi-site genes, respectively. This screen measures terminal-molecule depth; it does not require two different alternative sites to have positive counts in the same cell. There were 36,630 threshold-1 candidates with no interval overlap with a threshold-3 site, distinct from the difference in catalogue totals. The analysis kept the catalogue, eligibility and depth-supported populations separate (Fig. 8c).

#### Case 3: atlas-distant neuronal ends

The mouse-neuron analysis evaluated 16,738 native sites with site, barcode, tail-end, internal-priming, sequence and annotation evidence. An initial screen retained 8,525; a read-end-mode check retained 8,435; and 441 were more than 100 nt from a same-strand PolyA_DB entry. The reference atlas had been mapped from mm9 to mm10, and sequence coverage of that mapping was evaluated separately from distance to mapped atlas entries. Three candidates passed the stringent read and sequence screen, of which only the *Bcl11a* candidate lay in atlas-covered sequence. Its endpoint was supported by 33 distinct tail-read molecules in 33 neurons, 1,652 nt from a PolyA_DB site but 6 nt from an annotated GENCODE transcript end, indicating a gap in the bulk site catalogue at an already annotated RNA end. The full evidence screen, the two candidates outside mapped atlas coverage and the separate lower-support *AA387200* finding are documented in Supplementary Methods and Supplementary Fig. 17.

#### Independent numerical verification

For all six executions, numerical results were recomputed from native scPolyA outputs and checked against the reported analyses, including site inventories, UMI counts, threshold-comparison denominators and neuronal candidate evidence. One execution per question appears in Fig. 8; Supplementary Fig. 17 presents both agents’ analyses and the independent review findings. Numerical verification records are provided in Source Data; agent session transcripts are not included in that deposit. The 9,030-site Case 1 catalogue and the 9,047-site Case 2 catalogue arose from different native executions and must not be combined.

### Statistical analysis and reproducibility

Unless otherwise specified, intervals are 95%, tests are two-sided and *P* values are adjusted with the Benjamini–Hochberg procedure within the stated analysis family. Exact tests, resampling units, eligibility criteria, permutation counts and sensitivity analyses are specified above and in Supplementary Methods. Dataset-level replication was used for comparative benchmarking (*n* = 7–8), test-chromosome dataset-level replication for DeepCPS operating-point comparisons (*n* = 8), and patient-level replication for the principal cSCC analyses (ten paired single-cell patients and five for the primary spatial comparison). Multiple tissue sections from one patient were averaged before patient-level inference. Public datasets were selected for compatible 3′-tag sequencing, barcode/UMI information and established cell or spatial-unit lists; no formal power calculation was performed for these resource datasets. All analyses were computational, and experiment-assignment randomisation and blinding did not apply. Full software revisions, numerical tests and execution conditions are documented in Supplementary Methods and Supplementary Tables 2–6.

## Declarations

### Data availability

All analysed datasets are public (Supplementary Table 2). Poly(A) sites, count matrices, DeepCPS scores and figure source data will be deposited at Zenodo (https://doi.org/10.5281/zenodo.23771068). Panel-level values and extended diagnostic and exploratory tables are provided as Source Data; output schemas are in Supplementary Table 6, cSCC candidates in Supplementary Table 5 and tail-stratified support in Supplementary Table 1.

### Code availability

scPolyA v2.1.0, including the desktop application and the DeepCPS models (scoring and localisation ensembles with the calibration table), is available at GitHub (https://github.com/JeanWen-Lab/scPolyA) under the MIT licence, and the analysis scripts for every figure will be archived at Zenodo (https://doi.org/10.5281/zenodo.23771068). The same Zenodo record contains the source code of scPolyA revision 34c2f85 (tag scpolya-paper-freeze-20261021), used for the benchmark analyses. Revisions ce004fc and 0b0d766, used for some analyses, are its direct ancestors and differ from it only by fixes that do not change sites or counts. The revision used for each analysis is given in Methods and in the file manifest of the Zenodo record.

### Author contributions

J.Y. led the software architecture and development of scPolyA, including pipeline modularisation, performance optimisation, and the graphical user interface. J.Y. conceived and implemented the parameter-aware task execution and reuse framework and collaborated with J.W., H.Y., and L.L. on methodological refinements. J.Y. and L.L. jointly conceived the agent-assisted analysis workflow, which J.Y. designed and implemented. J.Y. contributed to figure preparation, manuscript drafting and revision.

H.Y. conceived and developed DeepCPS, including poly(A)-site scoring and cleavage-site localisation. H.Y. led the systematic evaluation of scPolyA and DeepCPS, including comparative benchmarking, model validation, and single-cell and spatial transcriptomic analyses. H.Y. led the biological applications, prepared most of the figures, and took the lead in interpreting the results, organising the findings, and drafting the manuscript.

L.L. collected and processed datasets for pipeline development and evaluation, contributed to spatial benchmarking, and jointly conceived the agent-assisted analysis workflow with J.Y.

J.W. conceived and designed the study and developed the original computational pipeline that formed the foundation of scPolyA. J.W. contributed to subsequent methodological development and interpreted the results together with H.Y. and J.Y. J.W. secured funding, supervised the research, and reviewed and revised the manuscript.

J.Y. and H.Y. contributed equally to this work. All authors revised the manuscript and approved the final version.

### Funding

This work was supported by the Australian Research Council (ARC) Centre of Excellence for the Mathematical Analysis of Cellular Systems (CE230100001), an ARC Discovery Project grant (DP220101388), a National Health and Medical Research Council (NHMRC) Ideas Grant (GNT2027770), and the Talo Scholarship and Talo Innovative Grant funded by Taiyang Zhang and Loong Wang.

### Competing interests

The authors declare no competing interests.

### Ethics approval and consent

Not applicable.

## Acknowledgements

We acknowledge the National Computational Infrastructure (NCI), Australia, for providing high-performance computing resources.

## Notes

https://github.com/JeanWen-Lab/scPolyA

https://doi.org/10.5281/zenodo.23771068

### Competing Interest Statement

The authors have declared no competing interest.

