## Supplementary Information (Supplementary Methods, Figures 1-19 and Tables 1-6) for "scPolyA enables efficient, interactive and agent-guided alternative polyadenylation analysis in single-cell and spatial transcriptomes"

**Supplementary Table legends** **63**

**1 Supplementary Table 1** **64**

**2 Supplementary Table 2** **65**

**3 Supplementary Table 3** **67**

**4 Supplementary Table 4** **68**

**5 Supplementary Table 5** **69**

**6 Supplementary Table 6** **70**

entries outside the species-defined standard chromosome set. A "sample" usually denotes one library or spatial section; it is not necessarily equivalent to one donor or biological replicate. Cells from different donors within one BAM are distinguished by their barcodes and downstream metadata. The same BAM must not be registered under multiple sample identifiers.

Three optional sample-table columns control preprocessing. `is_deduped` (default 0) skips UMI deduplication for inputs already processed; `--force-dedup-bams` overrides this declaration. `requires_split` (default 0) writes per-chromosome BAM files before deduplication, and `--force-split-bams` applies this to whole-sample BAM inputs. Whole-sample BAMs that skip deduplication are split regardless of `requires_split`; this option has no effect on inputs already supplied as per-chromosome BAM directories. `barcode_suffix` (default empty) distinguishes otherwise identical barcodes across samples. Corrected barcodes and UMIs are read from configurable BAM tags (`--cb-tag`, alias `--cell-barcode-tag`, default `CB`; `--umi-tag`, default `UB`). Cell calling, cell selection and barcode correction are completed upstream; scPolyA does not generate cell whitelists. Supported species are human, mouse, fly and worm; `--contigs` restricts the standard contig set and `--remove-mt` excludes mitochondrial contigs.

From the input GTF, the pipeline constructs gene, exon and coding-sequence (CDS) intervals, an annotation database and canonical-transcript records. Chromosome naming is adapted to the first input BAM (adding or removing `chr`), followed by filtering to the configured standard contigs. `MT` and `chrMT` are treated as mitochondrial names. Before reference construction, the evaluated implementation removes genes whose names begin with `Gm` or contain `Rik`, including their associated gene features. This case-sensitive rule applies to all species and uses `gene_id` when `gene_name` is unavailable. Such genes cannot subsequently receive site assignments.

For each protein-coding gene, the canonical transcript maximises the sum of CDS feature lengths (`end-start`); ties are resolved by the first transcript in genomic-start order. A gene without a CDS-bearing transcript has no inferred canonical stop codon. The stop coordinate is inferred immediately $3'$ of the annotated CDS (CDS end $+1$ on the plus strand or CDS start $-1$ on the minus strand), corresponding to the first stop-codon base where the GTF excludes the stop codon, as in GENCODE. The canonical terminal exon and this stop coordinate determine transcript-context annotation and the reported $3'$ UTR distance.

The optional `prepare-bulk` command derives strand-specific bulk RNA-seq-supported gene-locus and $3'$ UTR-window annotations from bigWig coverage (`--plus-bw` and `--minus-bw`). With `--bulk-gene-gtf`, a gene assignment is retained only when the site overlaps the corresponding bulk-supported locus; otherwise a placeholder is returned. With `--bulk-window-gtf`, overlap with a bulk-supported $3'$ UTR window determines assignment priority. These annotations constrain assignment but do not replace GTF-derived genes, CDS intervals, stop coordinates or canonical transcripts. Neither bulk annotation option was used for the analyses reported here.

#### Read preprocessing and UMI deduplication

Read deduplication is performed before site discovery so that PCR copies do not inflate either the evidence for a tail-read endpoint or the neighbouring coverage track. The rules below also establish how molecule identities are carried forward to the later, site-specific UMI correction.

Input BAMs are processed as sample-by-chromosome units in parallel. Unless a sample is declared pre-deduplicated, PCR duplicates are removed with UMI-tools `dedup --per-cell --extract-umi-method tag`, which groups reads by cell barcode, strand and $5'$ alignment position and collapses UMI networks within each group. The UMI-tools position convention accounts for soft-clipped bases. The default directional-network rule joins UMIs separated by one nucleotide when their read counts satisfy $n_a \geq 2n_b - 1$; the higher-count UMI represents the lower-count neighbour. The implementation also permits `adjacency`, `cluster` and `unique` methods (`--dedup-method`). The representative-read random seed is fixed by `--tools-seed` (default 2026); optional deduplication statistics are enabled by `--dedup-stats`.

Deduplication precedes extraction of tail reads and generation of either coverage track. Sample-specific suffixes, if requested, are appended to the configured barcode tag and the `CR` tag after deduplication and before tail extraction. Splitting, barcode rewriting and deduplication have independent worker limits (`--max-split-workers`, `--max-filter-workers` and `--max-dedup-workers`). Deduplication units are scheduled in descending BAM-index mapped-read count; other tasks follow the workflow's native ordering. All downstream computations use the deduplicated files or files explicitly declared as pre-deduplicated.

### Poly(A)-tail-read extraction

Tail-read extraction must recognise the observed end of an alignment without imposing the same filters on the much larger set of reads used for total coverage. The following options control which soft clips contribute direct endpoint evidence; they do not estimate the length of the underlying biological poly(A) tail.

Tail-read extraction exposes three independent criteria: the minimum observed soft-clip length and A/T count $a_{\min}$ (`--minpolyA`, default 10), the allowed number of non-A/T clipped bases $n_{\max}$ (`--nonA-allowed`, default 1) and minimum alignment mapping quality $Q_{\min}$ (`--minMAPQ`, default 0). All comparisons are inclusive. The study-specific values, strand orientation and distinction from biological tail length are given in the main Methods. Unmapped reads are excluded, but the extraction routine does not otherwise exclude secondary or supplementary alignments by flag. The mapping-quality threshold is specific to this extraction stage and is not inherited by total-read coverage or site UMI counting.

Tail extraction and merging have separate worker controls (`--max-extract-workers` and `--max-merge-workers`). Deduplicated input BAMs and their extracted tail-read BAMs are merged by chromosome across samples with `samtools merge`; chromosome chunks are concatenated with `samtools cat`. These cohort-level BAMs generate the pooled discovery tracks, while the deduplicated sample-by-chromosome files are retained for barcode-resolved counting.

### Coverage peak detection

The rare tail-read track and the total-read track are thresholded separately because they support different steps of site construction. The exact coverage scale, CDS-masking rule and threshold mode determine which segments of each track are available when the two are combined.

Strand-specific coverage is computed independently from all deduplicated reads and extracted tail reads using `bedtools genomecov -bg -split -strand`, pooling cells or spatial units for site discovery. Coverage $C(x)$ is per-base read depth, with spliced alignments contributing only aligned blocks. Before peak calling, total-read bedGraph intervals are removed if they overlap any annotated CDS by at least 1 nt on either strand (`bedtools intersect -v`); this removes the entire constant-depth interval, including any bases beyond the CDS. The tail-read track is not CDS-masked.

The `--coverage-scaling` option (default `none`) leaves both tracks unscaled or scales the total-read track (`expr`), the tail-read track (`polyA`) or both tracks (`both`) to reads per million. For track $r$, the scaling factor is $f_r = 10^6/N_r$, where $N_r$ is the unfiltered BAM-record count in the corresponding deduplicated or extracted-tail input (`samtools view -c`); the scaled coverage is $f_rC(x)$. Fixed thresholds are then supplied on the corresponding scale (`--scaled-depth-threshold`, `--scaled-depth-threshold-polyA`, or the respective `--unscaled-depth-threshold` options).

Regions are called separately on total-read and tail-read tracks. Fixed-threshold mode retains positions with $C(x) \geq T$. Dynamic mode derives a gene-specific threshold using coverage within each same-strand annotated gene $g$ of length $L_g$. In dynamic mode, positions outside gene intervals are excluded; positions belonging to overlapping genes are retained if they meet any eligible gene's threshold. Total-read statistics use the CDS-masked track. Available summary statistics are the mean, median, median of positive-depth positions and maximum:

$$\bar{C}_g = \frac{1}{L_g} \sum_{x \in g} C(x), \qquad \tilde{C}_g = \mathrm{median}_{x \in g}\, C(x),$$
$$\tilde{C}_g^{+} = \mathrm{median}_{x \in g,\, C(x) > 0}\, C(x), \qquad M_g = \max_{x \in g} C(x).$$

The ordinary median includes zero-depth positions, whereas $\tilde{C}_g^{+}$ excludes them (and is set to zero when none are covered). For the first three statistics, $S_g \in \{\bar{C}_g, \tilde{C}_g, \tilde{C}_g^{+}\}$ and $T_g = \max(F, \alpha S_g)$, where $F$ and $\alpha$ are the configured floor and multiplier (defaults 0 and 1). For `max_norm_fixed`, $T_g = \beta M_g$, where $\beta \in [0, 1]$ is the fraction of the within-gene maximum (default 0.3). The two tracks are configured separately using `--exp-peak-threshold-mode`, `--polyA-peak-threshold-mode` and their associated statistic, floor, multiplier and normalised-threshold options. The selected mode must have the corresponding threshold or dynamic statistic explicitly supplied; incompatible fixed and dynamic settings are rejected.

Above-threshold intervals are merged across configurable gaps. The total-read option `--merge-distance` is required and nonnegative; the tail-read option `--merge-distance-polyA` defaults to the same value. The study's scaling, 5%-of-maximum total-read peak threshold and track-specific gap settings are specified in the main Methods; tail-depth thresholds were varied as indicated for each analysis.

**Coverage-constrained site construction**

The interval-splitting step translates endpoint evidence into regions suitable for counting molecules. The one-base gaps and merging rules below determine whether neighbouring ends remain distinguishable or become a single reported site, which can change the effective resolution of downstream APA measurements.

The following rules specify the interval-splitting operation described in the main Methods. For a plus-strand total-read region beginning at $E_s$ and sorted overlapping tail-region ends $p_1 < \cdots < p_K$, the initial BED intervals are $[E_s, p_1)$, $[p_1 + 1, p_2)$, …, $[p_{K-1} + 1, p_K)$. On the minus strand, for sorted overlapping tail-region starts $s_1 < \cdots < s_K$ and a total-read region ending at $E_e$, they are $[s_1, s_2 - 1)$, …, $[s_K, E_e)$. Consecutive split intervals therefore have a one-nucleotide gap. When a tail region overlaps two total-read regions, it contributes a boundary to both without first being clipped to either total-read region. The selected $3'$ endpoint follows tail evidence; counting eligibility follows the full site interval.

The `--annotate-peak-source` option selects `resolved` (the package default) or `merged`. Merging occurs if the difference between the following BED start and previous BED end is no greater than `--merge-distance`. Thus the default 1-nt gap between split intervals permits them to be rejoined when the merge distance is at least 1 nt; the scientific implications of the evaluated merged mode are described in the main Methods. Tail-read and total-read support for a merged interval are sums over constituents, and its internal-priming fraction is their tail-read-weighted mean, $m = (\sum_j m_j n_j)/(\sum_j n_j)$. Read-to-site assignment is then performed against the final intervals; constituent UMI counts are not summed. In `legacy` mode, merging follows internal-priming filtering.

**Internal priming and site annotation**

An apparent A-rich RNA end may arise from priming within genomic sequence. The read-level score below quantifies that risk, while annotation determines which candidates contribute to terminal or intronic APA summaries. Keeping these decisions separate preserves intronic read evidence even when a candidate does not qualify for an intronic APA index.

Implementation parameters are the aligned-window length $k$ (`--misprime-in`, default 10) and required A/T count $a$ (`--misprime-A-count`, default 8). On the plus strand, the final $k$ aligned bases are examined; on the minus strand, the first $k$ aligned bases are examined for T. Shorter aligned segments contribute all available bases. Soft-clipped bases are excluded, and overlapping reads are not additionally screened by strand. For tail reads $R_i$ overlapping site $i$, the internal-priming fraction is

$$m_i = |R_i|^{-1} \sum_{r \in R_i} \mathbf{1}\{c_k(r) \geq a\},$$

where $c_k(r)$ counts the specified strand-oriented aligned bases; $m_i = 0$ when $R_i$ is empty. The study's filtering cutoffs and distinct rules for counting and summarising intronic candidates are specified in the main Methods.

Sites outside the permitted contig set are discarded. Gene assignment uses `bedtools closest -s -D a -t all`, with ties resolved by the smallest signed difference between the site endpoint and canonical stop coordinate (negative upstream in transcript orientation). A gene lacking an inferred stop codon is selected only when no eligible CDS-bearing alternative exists. Overlapping or abutting same-strand sites are reduced to the one with greatest tail-read support. The classification priority is terminal exon, other exon, intron, then the 2-kb region beyond the canonical terminal exon, requiring at least 1-nt overlap with the relevant canonical-transcript interval. Positions not meeting those criteria, or lacking a suitable assigned gene, receive the native `Intergenic` label (unassigned for the analyses here; Supplementary Fig. 1a).

The general filtering parameter `--misprimed-ratio-threshold` has a package default of 1. In default `legacy` mode, it is applied to all sites during peak calling. In `terminal-aware` mode, the filter is applied after gene-context assignment: non-intronic sites must meet this threshold for counting, while intronic sites remain countable and use `--intronic-misprimed-ratio-threshold` (package default 0.5) for eligibility in intronic APA summaries. The intronic threshold replaces the general threshold for that purpose. Study-specific overrides are stated in the main Methods and Supplementary Table 3.

#### Site-specific UMI quantification

Once the final site intervals are fixed, molecules are assigned to those intervals afresh. This matters for threshold comparisons: changing a site boundary can redistribute UMIs, so revised APA outputs cannot be obtained simply by adding counts from discarded or merged intervals.

The final site annotation is supplied to featureCounts with `-t peak -g peak_name -F GTF -s 1 -f -R BAM`, requiring at least 1-nt sense-strand overlap. Its assignment records feed the `umi_count_xr` wrapper, which performs a second directional UMI error-correction step (edit distance 1) within each barcode–site group, across read positions. This second step is independent of the initial `--dedup-method`. Ambiguously assigned reads and multimappers are omitted. In `terminal-aware` mode the counting targets include intronic candidates that may subsequently fail the intronic APA summary filter.

With `--count-matrix`, sparse site-by-barcode tables contain only nonzero UMI counts. Enabling `--summarize-matrix` in terminal-aware mode also writes retained annotations, terminal gene–barcode counts, PSI, weighted genomic 3′ UTR distance and both iPUI forms (Supplementary Table 6). The terminal gene–barcode total measures eligible terminal molecules, not total gene expression. A dense cell-by-gene table is written only below `SCPOLYA_MAX_DENSE_CELL_GENE_MATRIX_CELLS` (default $5 \times 10^7$ entries); otherwise a JSON omission record is saved. `--rna-gene-matrix` optionally generates a separate exonic-expression matrix. Read-to-site assignments are exported when `--annotate-bam` or counting is requested; intermediate featureCounts BAMs are removed. Relevant limits include `--n-workers`, `--n-workers-annotate` and `SCPOLYA_FEATURECOUNTS_ANNOTATED_BAM_THREADS`, which defaults to one featureCounts thread independently of `--n-threads-bam-annotation`.

#### APA measures

The native APA outputs distinguish an eligible alternative end with no observed molecules from a gene–barcode combination for which APA cannot be estimated. The conventions below explain how transcript context, signed genomic distance and sparse matrix representation affect those distinctions.

The PSI, usage-weighted genomic $3'$ UTR distance and site- and gene-level iPUI equations are given in the main Methods. Eligible sites are keyed by gene identifier and strand. The terminal support requirement for PSI and weighted distance is `--psi-min-gene-umis` ($u_{\min}$, default 1); these two outputs require at least two retained terminal sites per gene. iPUI uses terminal support from all eligible terminal sites, including single-site genes. The gene-total iPUI includes only intronic sites that pass the intronic summary filter, although excluded intronic candidates can still have counts in the site matrix.

For a site stored as BED interval $[b_i, e_i)$ and the inferred stop position $s_g$, the signed coordinate used for ordering is $d_i = e_i - s_g$ on the plus strand and $d_i = s_g - b_i$ on the minus strand. The weighted-distance calculation uses $D_i = \max(0, d_i)$; negative signed distances are set to zero only for that calculation. The two strand formulas have a one-nucleotide coordinate asymmetry. Every terminal site tied for the greatest signed distance is marked distal, including in single-terminal-site genes; this flag does not itself establish multi-site APA eligibility. The distance excludes splicing and is not a reconstructed transcript length.

The native outputs are sparse, and a missing row is not automatically zero. For an eligible gene–barcode pair with terminal support $U_{gc} \geq u_{\min}$, a retained terminal site absent from the PSI table has zero observed use. PSI and weighted length are undefined when $U_{gc} = 0$ and unreported when $0 < U_{gc} < u_{\min}$. Intronic summaries are generated for barcodes with terminal or intronic count evidence, including barcodes with counts at intronic sites failing the summary filter. Site iPUI is unavailable if the site fails filtering or has no retained terminal partner. If a partner exists, the pseudocount permits an iPUI with $U_{gc} = 0$, including a value of zero when both the site and terminal counts are zero. Gene-total iPUI likewise equals zero when both eligible intronic and terminal counts are zero. Unobserved gene–barcode combinations are not emitted, and audit rows for unrepresented intronic sites can lack a barcode or calculated index.

The `legacy` preset groups filtered sites by gene name without a transcript-context restriction for PSI and weighted distance, and does not report iPUI. Thus APA summaries from a legacy agent execution and a terminal-aware benchmark should be interpreted under their respective eligibility rules.

#### Statistical tests used in this study

The native site matrices provide the observations for biological comparisons, while the hypothesis tests were implemented in study-specific analysis scripts. Reproducibility therefore requires the definition of each gene-level ratio, its eligible samples and the unit whose labels are permuted or paired.

The biological tests used study-specific scripts, as stated in the main Methods. The separately released donor-level `scpolya-diff-apa` module (edgeR and limma [6, 7]) was not used for these results. Each contrast records its eligible-gene population and multiple-testing family. For an unpaired comparison of two cell or spot groups, non-proximal terminal-site usage was defined as the sum of UMIs at all retained terminal sites strictly distal to the most proximal site divided by UMIs at all retained terminal sites; sites tied at the minimum distance were excluded from the numerator. The group-level ratio was computed from pooled numerator and denominator counts. The test statistic was the difference in group ratios divided by a standard error estimated from cell-level numerator-minus-ratio-times-denominator residuals. Group labels were permuted while preserving group sizes and recomputing this studentised statistic, with eligibility fixed from the observed labels. For $B$ permutations, a two-sided empirical $P$ value was $(1+e)/(B+1)$, where $e$ is the number of permuted absolute statistics at least as large as the observed value, allowing a numerical tolerance of $10^{-9}$. Benjamini–Hochberg correction was applied across eligible genes within each comparison. Analogous ratio tests compared IPA counts with IPA-plus-tandem counts and alternative-last-exon (ALE) counts with ALE-plus-tandem counts, with separate eligibility and multiple-testing families.

For paired-subject analyses, counts were pooled separately by subject and condition. Distal usage was transformed for each subject using $\log[(d+0.5)/(n-d+0.5)]$, where $d$ is the distal UMI count and $n$ is the total terminal UMI count. Condition differences in this quantity were tested across subjects with a two-sided one-sample $t$-test, followed by Benjamini–Hochberg adjustment across eligible genes. Label-permutation controls maintained patient membership and condition sizes and recomputed pseudobulks, eligibility, tests and adjusted $P$ values; these controls evaluated the number and directions of discoveries, without replacing the primary gene-level $P$ values. Aggregate APA contrasts used paired two-sided $t$-tests with $t$-based confidence intervals, after averaging section-level differences within patients when multiple sections were available. Wilcoxon signed-rank and sign tests provided sensitivity analyses. The analysis-specific eligibility thresholds, permutation numbers and adjustment families are specified in the corresponding supplementary sections.

### Task reuse and desktop application

Changing a discovery threshold can alter the set of candidate endpoints, the intervals assigned to them and their downstream UMI measurements. The task graph records which computational results can be reused and which must be regenerated for a revised setting. The implementation details below distinguish safe reuse under recorded task identities from changes that require fresh run state. We then distinguish the GUI's preview of a rerun from the separate experiment that actually executed one and checked its outputs.

#### Task identity and selective reanalysis

Task reuse is based on the parameters and completion records declared for each stage. It is not a general fingerprint of every byte in the input data, making the boundary between a valid parameter change and a new dataset important for interpreting the rerun experiments.

Task identities incorporate the pipeline stage, task type, declared parameters and identity fields such as sample and chromosome. SHA-256 hashes and completion records are stored in `<output-dir>/<run-id>/state/state.sqlite`. A recorded identity permits reuse, with stage-specific output checks. Peak combination and resolution, BED-to-bigBed and BED-to-GTF conversions, `annotate_slices` and matrix-summary tasks additionally require the expected output files to exist and be nonempty; tail extraction checks its BAM and index. Reference preparation, splitting, deduplication, barcode rewriting, coverage calculation, scaling, CDS masking and per-track thresholding rely on completion records without these additional output-presence tests. The separate hybrid reuse gate is disabled by default.

Parameters trigger recomputation only where included in a task identity or propagated derived configuration; downstream task identities are evaluated independently. Deduplication method, extraction criteria, coverage scaling and thresholds propagate to relevant dependencies, but seed, optional deduplication statistics and barcode-tag choices are not propagated to all dependent tasks. Concurrency limits are not part of task identities. Input registration and reference preparation have parameter-free global identities; input bytes are not fingerprinted. Fresh run state is required for changes to input BAMs, sample registration, gene GTF or barcode-tag settings, since the reference-change guard does not reliably detect the latter changes. These restrictions applied to the reported reuse experiments.

The desktop UI and Python API expose a reuse-and-recompute preview, which is not a separate native feature of the `run` CLI. `--repair-moved-run` updates the saved paths when a completed run directory is relocated; its dry-run preview reports path repair, not computational task reuse.

#### Desktop application

The main Methods describe the NiceGUI workspace, igv.js browser and Site scores view. The application evidence derives from two different datasets. The main GUI panels (Fig. 3) show a completed threshold-3 mouse-neuron run, including the *Gnb1* interval chr4:155,557,150–155,559,450, and a *preview* of a threshold change. The browser displays native counts pooled over cells and genomic distances from the first stop-codon base. The Site scores panel used scPolyA v2.1.0 and the scoring weights shipped with that version.

Supplementary Fig. 18 instead records an *executed* GUI rerun of the full 10x Genomics GEM-X 3′ v4 Human Donor1 PBMC library (5,710 cells; GRCh38 and GENCODE v32), changing `unscaled_depth_threshold_polyA` from 3 to 1 while retaining `minpolyA`=5. All captures were made with scPolyA v2.1.0. Numerical comparison with an independently started threshold-1 command-line analysis covered the eight primary-key-aligned native tables listed in the main Methods. Supplementary Fig. 19 and Source Data document the row-level checks and matching numerical values. The GUI illustration, the completed rerun and the DeepCPS scoring view are separate demonstrations.

### DeepCPS architecture, training and extended evaluation

The main Methods specify the DeepCPS networks and their chromosome-partitioned training design. Reproducing the evaluation also depends on how released models are adapted to the same candidate windows, how uncertainty is resampled, which scores are calibrated and which raw scores define selection thresholds. These details matter because reference-atlas scoring, cleavage-position localisation, site retention and recurrence in another library measure distinct properties. The following sections document the controls and denominators for each analysis without repeating the network architecture.

#### Input encoding, stopping rules and calibration

Training stopping rules and score calibration affect the reproducibility of the released model outputs even when the network architecture is fixed. Calibration is a separate step from model fitting and site-selection threshold choice, so we specify each operation independently.

The model inputs, layer dimensions, objectives and chromosome-partitioned training design are given in the main Methods. The remaining settings specify exact encoding and stopping behaviour. Ambiguous nucleotides (N) were encoded as zeros across all four nucleotide channels. The scoring convolutional stem used batch normalisation and ReLU. Three scoring models were fitted with random seeds 1–3; training stopped after four successive epochs without improved validation AUPRC. For the localisation model, the early-stopping patience was six epochs without improved validation negative log-likelihood, and the same three seeds were averaged. The localised coordinate was extracted from the average positional distribution, not by averaging three independently selected endpoint coordinates.

Isotonic regression for probability calibration was trained using validation-chromosome labels. Calibration was evaluated on the held-out test set using expected calibration error computed over ten equal-width probability bins. The calibrated estimate corresponds to proximity to a reference site within 25 nt, following the training label definition. Catalogue-selection operating thresholds were derived using uncalibrated ensemble outputs; the calibration results and threshold-based selection analyses therefore address different outputs of the same scoring model.

#### External-model adaptation, ablation controls and position rules

Comparisons with existing models require matching the candidate and evaluation set without silently changing the released model definitions. Small differences in input-window length and coordinate anchoring are therefore specified before interpreting differences between model scores or predicted cleavage positions.

To adapt the 201-nt DeepCPS-centred window to released scoring models, the first nucleotide was removed for DeepPASS and the last nucleotide for DeepPASTA, producing their respective 200-nt inputs [11, 12]. Human and mouse DeepPASS models were used in their released forms. The fitted logistic-regression sequence baseline used three features: presence of AATAAA or ATTAAA from $-50$ to $-1$ nt, the count of downstream adenosines from $+1$ to $+20$ nt and the longest A run within $\pm 10$ nt. This fitted baseline is distinct from the deterministic sequence-ranking rule used for independent PBMC recurrence.

The embedding ablations used frozen Nucleotide Transformer v2-50m features [13] projected to 32 dimensions, or frozen DeepPASS or DeepPASTA representations feeding a trained prediction head. The ablation arms shared the main Methods' chromosome partitions, convolutional backbone and training seeds; cross-species and cross-protocol training used only the designated training-side species or protocol. No change in the reported model input channels constituted an independent validation cohort.

For cleavage-position localisation, the PAS-offset rule selected the first upstream AATAAA or ATTAAA motif from $-40$ to $-10$ nt and placed the predicted cleavage coordinate 21 nt after it. APARENT v1 [14] used input anchoring selected from validation sites. These anchoring choices were kept fixed for test-set and independent PBMC evaluations.

#### Resampling units and tail-support strata

Nearby candidate sites can share their gene and sequence context. Locus-cluster resampling was used for paired performance differences, while read-support stratification asked whether model discrimination remained informative when direct tail-read counts were small.

The classification and localisation outcomes, together with the in silico substitution definition, are set out in the main Methods. For the paired scoring analysis, the 2,000 resamples clustered observations within 5,656 species–gene loci, giving a minimum attainable two-sided empirical $P$ value of 2/2001 before Benjamini–Hochberg correction within each comparison family. The support-focused subset included 9,108 atlas-positive and 3,694 A-rich atlas-negative candidates supported by at least three poly(A)-tail reads. Read-support effects were also examined in 11 individual bins (1–10 and $> 10$ reads), with five grouped categories displayed in Supplementary Fig. 3b. The support-focused denominator must not be substituted for that of the full labelled classification test set.

The localisation uncertainty analysis resampled 2,438 loci; the paired exact-coordinate comparison used an exact McNemar test on success and failure for the same test sites. The observed position was the modal deduplicated tail-read endpoint within each eligible reference-proximal site. Those endpoints were used as a read-based positional target, as distinguished from direct chemical measurements in the main Methods.

#### Transfer-test denominators and operating-point statistics

Protocol transfer, catalogue selection and usage precision were evaluated as separate questions. A site eligible for candidate scoring is not necessarily eligible for positional assessment or for distal-usage estimation, so their denominators and statistical tests are recorded separately.

Transfer tests held the trained ensembles and the validation-selected selection rules fixed. For the protocol-transfer panel (Fig. 2g), scoring performance on the eight evaluated dataset partitions was compared with DeepPASS using locus-cluster bootstrap tests followed by Benjamini–Hochberg adjustment across datasets. Paired exact-nucleotide localisation success was assessed against the original site endpoint using exact McNemar tests. In each evaluation, the threshold-1 scoring set was filtered for native internal priming, whereas the localisation set required atlas proximity and at least ten tail endpoints; these denominators should not be combined.

The two uncalibrated site-selection thresholds and their validation-derived matching targets are described in the main Methods. Their downstream effects were compared across eight held-out dataset partitions using exact Wilcoxon signed-rank tests with Benjamini–Hochberg correction within metric families (Supplementary Fig. 6a–c). Catalogue retention, PolyA_DB 3 proximity, annotation composition and distal-usage error after molecule downsampling were assessed as separate outcomes: a larger site catalogue does not necessarily provide more precise per-cell APA estimates.

#### Independent PBMC replication

Because atlas annotations supplied DeepCPS scoring labels, recurrence in independently generated PBMC data offered a complementary read-derived outcome. The comparison used held-out chromosomes, although the same libraries had contributed training examples on other chromosomes. It therefore evaluates cross-dataset site detection under a genomic holdout, not transfer to an entirely unseen library.

The 10x–Drop-seq recurrence analysis used the same held-out chromosome selection and native internal-priming filter as the scoring study (Fig. 2h–j; Supplementary Fig. 6d–f). Candidate rankings used a fixed-seed tie break. The deterministic sequence-ranking baseline first prioritised AATAAA or ATTAAA in the $-50$ to $-1$ nt upstream window and then fewer downstream adenosines at $+1$ to $+20$ nt; it was not the logistic-regression baseline fitted for held-out scoring. The tail-depth rule retained candidates supported by at least three tail reads. For the matched-depth analysis, the five bins were 1, 2, 3–5, 6–10 and $> 10$ reads; higher- and lower-score candidates within each bin were defined using the validation-selected uncalibrated threshold 0.292. Recurrence proportions have 95% candidate-bootstrap intervals.

The independent localisation comparison (Fig. 2k) searched for a same-strand site within 50 nt of the candidate and used the read-end mode of at least ten deduplicated tail reads at that independently observed site. The original site endpoint, the candidate's own observed mode, DeepCPS, the fixed PAS rule and APARENT were assessed against that single read-based position; APARENT anchoring was set using validation data. This independent-reference definition was held constant across all reported positional methods.

### Benchmark inputs, comparisons and resource measurements

The benchmarking experiments compare methods that accept different read structures and report different forms of poly(A)-site coordinates. A common biological input and a harmonised endpoint definition are necessary, but each tool must also be run within its supported workflow. We therefore document the input adaptations, successful and incomplete executions, reference and sequence metrics, cell-type comparisons and resource-accounting boundaries. These details define precisely which comparisons are like-for-like and how their conclusions should be interpreted.

#### Single-cell and spatial datasets

The single-cell benchmark comprised eight publicly available oligo(dT)-primed 3′-tag datasets from four protocol families (Supplementary Fig. 2c and Supplementary Table 2): 10x Genomics GEM-X 3′ v4 human PBMCs (5k Human Donor1) and mouse brain (10k Mouse Neurons); Drop-seq human PBMCs (GSE132044) and mouse P14 retina (GSE63472); Microwell-seq human and mouse spleen (GSE134355 and GSE153562); CEL-seq2 human pancreatic islets (GSE85241); and CEL-seq mouse intestinal organoids (GSM1524282). Datasets comprised 96–12,441 cells and 5.1–506.5 million retained aligned reads. Human libraries used GRCh38/GENCODE v32, mouse libraries GRCm38/GENCODE M23. Cell Ranger provided alignment and barcode correction for the 10x libraries; STARsolo 2.7.11b was used for the other single-cell protocols, with protocol-specific barcode and UMI structures. All compared methods received the same coordinate-sorted, primary BAM alignments with corrected barcode and UMI tags and the same cell list, subject to method-specific input conversions specified below.

Eight spatial sections from seven protocols were analysed (Fig. 5; Supplementary Table 2): human cSCC Visium P4 replicate 1 and legacy Spatial Transcriptomics (ST) P2 replicate 1 (GSE144239, within GSE144240) [22]; Visium sagittal mouse-brain anterior (V1_Mouse_Brain_Sagittal_Anterior); Visium HD 3′ Mouse Brain Fresh Frozen; Slide-seqV2 mouse testis (SRR12805494) [15]; Stereo-seq mouse brain replicate 1 (GSE256319, GSM8093734, SRR28048223) [16]; DBiT-seq E12 mouse embryo (SRR10517906) [17]; and HDST mouse olfactory bulb replicate 1 (SRR10125060) [18]. Units were 55-$\mu$m Visium spots, approximately 110-$\mu$m ST spots, 50-$\mu$m Visium HD bins, approximately 50-$\mu$m Slide-seqV2 bins, 50-$\mu$m Stereo-seq bin100 bins, 50-$\mu$m DBiT-seq pixels, and HDST bins of 25×25 wells (2-$\mu$m beads). Spatial inputs and exact unit counts are listed in Supplementary Table 2. Mouse sections used GRCm38/GENCODE M23; Visium and Visium HD BAMs came from their respective Space Ranger workflows, with Visium HD aligned in Space Ranger 4.1.0.

For the spatial cSCC libraries, the published data were realigned as described in the cSCC section. Slide-seqV2 aligned BAMs were converted to FASTQ without discarding soft clips, realigned with STARsolo and binned for analysis. For Stereo-seq, the first 20 million read pairs were used; 10 terminal nucleotides were removed from the 110-nt read 2 (`cutadapt -u -10`) before alignment because they matched an appended DNBSEQ sample-index sequence. These adaptations and barcode layouts are detailed in Supplementary Table 2.

**Matched method evaluation**

The comparison panel was selected to represent distinct established strategies for RNA-end discovery and quantification, not every available method. All eight single-cell datasets were evaluated with scPolyA and seven other tools spanning tail-read-guided discovery, read-coverage and fragment-distribution approaches, sequence-filtered peak calling and reference-guided quantification. Five other tools were run on spatial datasets where applicable. Site-coordinate metrics were harmonised across outputs, but the algorithms and supported settings differed. DeepCPS was evaluated separately against released models and baseline rules on matched candidate sets. The scPAISO Read2-only workflow, scDeepAPA and PolyaID were not included in the completed head-to-head experiments; the numerical comparisons therefore apply only to the methods actually evaluated. Where supported, minimum mapping quality was set to 10 and minimum poly(A) evidence to five bases. Full versions, settings and execution scope are in Supplementary Table 3.

*scPolyA.* The single-cell and spatial benchmark used revision 34c2f85 or ancestors ce004fc and 0b0d766, with intervening fixes that did not change site or UMI outputs (Code availability). The evaluated discovery and counting thresholds are specified in the main Methods; their software controls, defaults and edge cases are documented above and in Supplementary Table 3. Runs used GTF-derived references, corrected CB/UB tags, directional deduplication (seed 2026), `terminal-aware` summaries, `merged` site intervals and an unscaled tail track with scaled total-read coverage. The primary discovery threshold was 1 unless threshold 3 is explicitly indicated. Bulk-supported annotation and DeepCPS site filtering were disabled. Both full site-count matrices and APA summary outputs were requested.

*polyApipe.* polyApipe commit a6dd4bb and polyApiper 0.1.0 used the prepared tagged BAMs and an organism reference derived from the matched GTF. Parameters were tail-read depth 1, minimum poly(A) length 5, mapping quality 10 and no non-A bases (`--non_A_allowed 0`, its default). polyApiper used native defaults, including at least one tail read per site and fixed 250-nt counting regions. Single-cell sites came from its default output without an additional mispriming filter; spatial sites excluded those flagged as misprimed. Reported coordinates were the strand-aware 3′ peak boundaries. polyApipe returned no sites from the CEL-seq organoid library because the clipped tails ended in an ambiguous N that was not permitted by its extraction filter.

*scAPAtrap.* scAPAtrap 0.2.0 received the prepared BAMs with `tails.search="no"`, coverage cutoff 10, minimum 10 cells, minimum count 10, maximum peak width 1,000 and `d=100`. Single-cell analyses used the native `scAPAtrap()` function. Spatial execution used per-cell deduplication, chromosome-serial peak calling and chunked feature assignment to limit memory; Visium HD was run through the native function with the same tail-search option. Reported coordinates were peak 3′ ends. The Visium brain analysis did not complete.

*Sierra.* Sierra 0.99.27 used the matched GENCODE GTF and species-specific BSgenome reference, with junction count/proportion 50/0.05, peak count/proportion 200/0.05 and coverage cutoff 500. The single-cell analysis used the native `CountPeaks` output and retained its pA/pT flags as annotations without a further A/T filter. Spatial analysis applied the pA/pT filter. Site coordinates were the fitted peak 3′ ends as in Bi et al. [21].

*SCAPTURE.* SCAPTURE commit 8845e84 was run through `PAScall`, `PASmerge` and `PASquant` with GTF-derived annotation, a 400-nt window, overlap 0.5 and no external poly(A)-site database. Only DeepPASS-positive exonic or intronic peaks were retained, with coordinates at the peak 3′ end. SCAPTURE was evaluated on single-cell datasets only.

*SCAPE.* SCAPE 1.0.0 was run using `apamix` on UTR intervals extracted from a coordinate-sorted GENCODE GTF, allowing one to five sites per UTR and otherwise using native defaults. Sites were taken after the official `group_pa.py` collapsing step (`--group_threshold 100`). SCAPE was evaluated on single-cell datasets only.

*MAAPER.* MAAPER 1.1.1 received the matched BAMs, a PolyA_DB reference and exon annotations from the same GTF. It used single-sample mode (`run="all"`), minimum 50 reads per gene, 25 reads per PAS and PAS fraction 0.05. For Visium brain, two groups selected from a $k = 3$ clustering were split into two pseudoreplicates each, yielding four BAM inputs. MAAPER outputs coordinates from its reference, achieving 100% agreement with PolyA_DB 3 by construction. This value was included for comparison but was not interpreted as independent evidence of poly(A)-site discovery accuracy.

*scUTRquant.* scUTRquant 0.5.0 used the `utrome_hg38_v1` reference for human and `mm10_v2` for mouse, minimum UMI count 1 and BUS correction enabled. The method received prepared BAMs or, for Microwell-seq and Stereo-seq, FASTQ inputs converted from the tagged BAMs. Its evaluated site coordinates were unique strand-specific 3′ endpoints of expressed UTRome features.

scTail [19] was not included because its tested workflow requires read 1 to extend into cDNA. Among the eight single-cell libraries, only the two Microwell-seq libraries had sufficiently informative read-1 structure; 10x and Drop-seq read 1 contained barcode and UMI only (20–31 nt), and CEL-seq contained only 15–41 nt of cDNA. The distinction concerns the evaluated scTail workflow and does not imply that other recently released APA methods are incompatible with these inputs.

#### Site and atlas evaluation

A coordinate comparison becomes meaningful only after each caller's interval or peak definition has been translated into a common strand-aware endpoint. Atlas proximity and motif organisation then provide complementary observations about the resulting site map, with shifted references supplying a negative control.

For method-to-method comparisons, each reported site interval was reduced to one strand-aware 3′ coordinate: BED end on the plus strand and BED start +1 on the minus strand. Identical coordinates were collapsed and mitochondrial sites removed. All tools were assigned genomic contexts against the same gene annotation, with priority 3′ UTR, 5′ UTR, other exon, intron and intergenic. Reference support was defined as the fraction of reported coordinates within 25 nt of the nearest same-strand PolyA_DB 3 coordinate [9]. A negative control shifted reference sites by 500–5,000 nt. Secondary comparisons used PolyASite 2.0 [10] and PolyASite 3.0 (release 15 September 2024) [20], at stringency at least 47 (human) or 46 (mouse), with within-gene shifted controls. A match to a reference site indicates proximity, not independently established cleavage accuracy.

Threshold-3 scPolyA sites provided a within-method sensitivity reference in Supplementary Figs 7 and 11. scPolyA was compared with each evaluated method across datasets completed by both using two-sided exact Wilcoxon signed-rank tests ($n = 7$ or 8, smallest attainable two-sided $P = 0.0156$ or 0.0078). Benjamini–Hochberg adjustment was applied within each panel's specified family (13 tests in Fig. 4c, spanning PolyA_DB 3 and PolyASite 3.0); PolyASite 2.0 analyses were unadjusted. For matching to annotated transcript ends in Supplementary Fig. 7f, SCAPTURE minus-strand positions were shifted −1 nt in genomic orientation; no other method- or strand-specific offset was applied.

Canonical-PAS frequency was the fraction of sites carrying AATAAA or ATTAAA in the 50 nt upstream. Motifs are reported here using genomic DNA notation, corresponding to AAUAAA and AUUAAA in RNA. Strand-oriented base profiles counted A, C, G and T/U at positions −60 to +40 nt around each site. Up to 20,000 sites were sampled per tool and dataset (all sites for Sierra, SCAPE and SCAPTURE). The profile-distance metric in Fig. 4e was adapted from Bi et al. [21]: for each dataset, the site-based profile $p_b(x)$ was compared with the species-matched PolyA_DB profile $q_b(x)$ using

$$D = \frac{1}{|X|} \sum_{x \in X} \sum_{b \in \{A,C,G,U\}} \frac{\{p_b(x) - q_b(x)\}^2}{p_b(x) + q_b(x)}, \qquad X = \{-60, \ldots, +40\},$$

with $0/0 = 0$. The median $D$ across datasets summarised each method. Downstream A-richness was defined as at least 12 A bases at positions +1 to +20 nt. Overlap between methods was examined with pairwise same-strand containment within 25 nt and single-linkage groups of the six de novo tools' coordinates, linking consecutive sites no more than 25 nt apart.

For the strict atlas-absent examples (Fig. 4f), threshold-1 scPolyA candidates on primary chromosomes were required to meet four criteria. First, they had to lie in a named protein-coding host gene at least 200 nt from annotated transcript ends and at least 50 nt from annotated splice donors. Second, they had to be more than 100 nt from any same-strand PolyA_DB 3 coordinate or PolyASite 3.0 entry at any stringency. Third, they required AATAAA or ATTAAA at positions −50 to −1 nt and fewer than 12 A bases downstream at +1 to +20 nt. Fourth, at least ten independently deduplicated tail-read 3′ ends had to fall within 25 nt, comprising at least 50% of all tail-read support at the site. Reads were processed under scPolyA tail-read criteria using MAPQ $\geq 10$ and expression-labelled cells. Other tools were judged to recover a focal site if their nearest same-strand coordinate lay within 25 nt. The stringent atlas-absent candidate screen used threshold-1 sites, whereas the *Rorb* locus displayed in main Fig. 4f uses the threshold-3 catalogue. For this display, a coordinate was labelled reference-supported if an annotated transcript 3′ end or a PolyA_DB 3, PolyASite 2.0 or PolyASite 3.0 entry lay within 30 nt. Otherwise, it was labelled *de novo*.

Reference-atlas tissue coverage was assessed from the PolyA_DB 3.2 sample list and PolyASite 3.0 sample annotations accessed 21 October 2026 (Source Data). PolyASite 3.0 had no mouse library explicitly labelled retina, brain, testis or embryo, and 27 libraries had unknown tissue labels. Tissue-coverage analysis used threshold-1 sites meeting the strict evidence criteria above except for a minimum of three tail reads, no requirement for a named protein-coding host and no atlas-distance exclusion. Intronic, noncoding and intergenic sites were included. The far fraction was the proportion more than 100 nt (or, in sensitivity analyses, more than 25 nt) from all same-strand PolyA_DB 3 and PolyASite 3.0 entries. Spearman correlations related this fraction to $\log_{10}$ supported-site number and $\log_{10}$ aligned-read depth across the ten mouse datasets, treating the dataset as the observational unit.

In the spatial analysis, a site was present in a barcode-defined spot, bin or pixel if it had at least one UMI. Successful output required readable native result files, not merely a zero process exit code. Expressed scUTRquant UTRome features were classified with the harmonised downstream annotation for comparisons. Section-paired statistical tests followed the same procedure as in the single-cell analysis, with Benjamini–Hochberg adjustment across nine tests in Fig. 5d. For Supplementary Fig. 2a, read lengths of the mouse-brain Visium library were determined from the first 2,000 Space Ranger BAM records using the `CR`, `UR` and sequence fields. Clips at least 60 nt long, omitted from Supplementary Fig. 2d, comprised two of 2,039,210 extracted tail reads in 10x mouse neurons and none in the other libraries.

### Cell-type APA comparisons

The benchmark also asks whether alternative-site measurements recover biological differences, not only whether site coordinates match an atlas. Cell identities were derived from expression, independently of APA, and each tool's APA tests were carried out on its own eligible site-count matrix.

Cell identities for the single-cell benchmark were assigned from expression derived from the same alignments. Normalised, log-transformed gene counts were reduced to 2,000 highly variable genes and 30 principal components, followed by a 15-nearest-neighbour graph and Leiden clustering at resolution 1. Clusters were assigned identities from marker-gene modules; cell classes containing fewer than 20 cells were excluded. For consistent comparison, sites reported by every method were mapped to the same gene model. A site required at least 5% of its gene's UMIs, and a gene required at least two retained sites and sufficient expression (300 UMIs in at least 100 cells for droplet and Microwell-seq datasets; 50 UMIs in at least ten cells for plate-based datasets).

Per-cell representations comprised site usage, the fraction of terminal UMIs at sites strictly distal to the most proximal terminal site, usage-weighted 3′ UTR length, intronic fraction $I/(I+U)$ and proximal-site usage. These measures were stabilised with empirical-Bayes shrinkage ($\alpha = 2$) and pooling across 30 nearest neighbours found independently for each representation. The abundance-preserving null held each cell's per-gene UMI total constant and reassigned each UMI to a site according to that gene's dataset-pooled site-use distribution. Three null replicates were generated, and features, neighbours and classifiers were recomputed for each. Gene-level neuron-versus-glia comparisons used scPolyA threshold-3 counts in the 10x mouse-neuron dataset, comprising 6,315 neurons and 3,184 glial/progenitor cells. For each gene, pooled non-proximal terminal-site fractions were tested by the two-sided studentised permutation procedure (2,000 cell-label permutations). Eligible genes needed at least 20 terminal UMIs and at least ten expressing cells in each class; Benjamini–Hochberg FDR was controlled at 0.05 separately for each method. Null replicate 1 was subjected to the same gene test; all three null replicates were used in the cell-identity audit. Directional agreement between two methods was the fraction of genes significant in both with the same sign of neuron-minus-glia effect.

For the loci in Fig. 4i–k, tail-read coverage was recomputed separately for neurons and glia/progenitors with `bedtools genomecov -split` on UMI-tools-deduplicated reads. Upstream aligned blocks of spliced tail reads contributed to the tracks; aligner duplicate flags were not used. Total-read coverage used deduplicated sense-strand alignments with MAPQ at least 10. Each track has its own zero-to-maximum vertical axis for the displayed window.

#### APA classification and locus examples

The locus examples distinguish different transcript structures that can produce a change in RNA-end usage. Their classification rules and support screens are described separately from the genome-wide comparisons because the selected examples illustrate mechanisms; they are not an additional independent validation cohort.

The illustrative APA-type analysis used threshold-3 scPolyA sites and their counts, with the canonical transcript defined by the longest summed coding-sequence length. Site classes were assigned by downstream analysis scripts with the precedence tandem $3'$ UTR, alternative last exon (ALE) and intronic polyadenylation (IPA; Supplementary Fig. 1b). Tandem sites lay in the canonical terminal exon or the 2-kb downstream window. ALE sites occurred upstream of the canonical terminal-exon start and overlapped the last exon of a different annotated transcript on the same strand, including positions annotated as canonical introns. Other sites in canonical introns were IPA. Internal-priming scores had to be at most 0.7 for intronic sites (including intron-localised ALE sites) and at most 0.9 otherwise. Across this classification, 1,000 ALE sites were in canonical introns, 164 in upstream exons and 44 in intergenic contexts.

For each cell class, IPA and ALE usage were the UMIs at intronic or ALE sites divided by those plus tandem-site UMIs, and tandem distal usage was the fraction of tandem-site UMIs at distal sites. Gene-level differences were tested with two-sided studentised statistics over 2,000 cell-label permutations, with Benjamini–Hochberg adjustment within each APA class. Displayed usage values use only the focal IPA or ALE site and a reference tandem site; because the tests used a single library with cells as units and intronic usage may partly reflect unspliced RNA, the $q$ values in Fig. 4j,k served to select illustrative loci and genome-wide IPA and ALE results are not reported.

Illustrative genes were selected by prespecified evidence filters. IPA candidates were restricted to the final two canonical introns; ALE examples excluded events defined by a single-exon transcript or occurring in the canonical first exon. Candidate pairs were first enumerated among genes with $q < 0.01$ and at least 20 pair UMIs in each group. The focal site then required fewer than 12 downstream A bases in 20 nt, a reference-atlas entry within 25 nt, at least 20 tail-read endpoints within $\pm 25$ nt in the better-supported class and five in the other class, absolute pair-usage difference of at least 0.10 and gene-level $q < 0.05$. IPA examples additionally had to be at least 200 nt from any annotated same-strand transcript end. *Arglu1* satisfied the annotation criterion but was retained as an illustration despite 17 tail reads in the higher-support class and a pair-usage difference of 0.077, both below the general illustrative thresholds. These loci were selected to illustrate APA classes, not as independent validation.

#### Runtime measurement and resource limits

Computational comparisons ran on a 48-core server with 247 GiB of physical RAM, subject to a 16-CPU quota for all single-cell tool measurements. scPolyA, polyApipe, scAPAtrap, Sierra, SCAPTURE, SCAPE, MAAPER and scUTRquant ran serially with checks for other activity on the assigned cores and a 256-GiB cgroup memory limit. Each value represents one run. Peak memory was measured as the cgroup maximum, including page cache, and is therefore an upper bound on process-resident memory.

For Fig. 6a,b, native method-stage timing spans differed by software (Supplementary Table 3). scPolyA was timed through its complete invocation, including reference preparation, site counting and enabled APA summaries. polyApipe and Sierra timings summed their native calling and annotation or junction and peak-analysis stages, with memory reported as the maximum across those stages. MAAPER, scAPAtrap (tail search disabled) and scUTRquant were timed through their native analysis stages, with scUTRquant excluding BAM-to-FASTQ adaptation. SCAPTURE was timed through PAScall, PASmerge and PASquant, and SCAPE through apamix, excluding its subsequent site-collapsing step. Runs producing no sites remained visible in the figure but were excluded from median summaries and ranks. The two reference-based methods were not included in rankings among the six de novo methods. Ties were assigned their minimum rank and each method's within-dataset ranks summarised by the median across datasets. Aligned-read-normalised times are provided in Supplementary Table 3 and Source Data.

Stage shares in Fig. 6c used the sum of logged stage intervals, excluding invocation startup and finalisation. Spatial profiles used threshold 1, 16 CPUs, 128 GiB of memory and no swap. The Stereo-seq input for stage profiling retained its final 10 nt; the site benchmark used the trimmed read described under datasets.

For the selective-rerun experiment (Fig. 6d; Supplementary Table 4), a full 10x PBMC threshold-1 run was compared with an unchanged rerun and a rerun increasing the tail-read threshold from 1 to 3. Each rerun used a separate copy of the completed baseline directory, with stored paths repaired through `--repair-moved-run`. All were performed using scPolyA revision 34c2f85, a 16-CPU quota and 256-GiB memory limit, and each completed the count matrix and APA summaries. Reported durations sum the same seven logged stage categories for each run, and executed-task counts were obtained from invocation logs.

For scaling experiments (Fig. 6e,f), `samtools 1.19.2 view -s` with seed 2026 subsampled the prepared 10x mouse-neuron BAM to 1, 5, 10, 25 and 50% of its read records; the complete library was the 100% point. Records sharing a read name were retained or discarded together. Each level used the same cell list, reference, threshold-1 settings and a new output directory. BAM preparation was outside the timing interval. Runs were serial with a 16-CPU quota and 256-GiB limit. Least-squares lines with intercepts were fitted to six points: wall time $= 8.6+0.26N$ minutes and peak memory $= 15.4 + 0.18N$ GiB, with $N$ in millions of aligned reads (both $R^2 = 0.99$). Peak memory included cgroup page cache.

### cSCC data processing and detailed statistical analyses

The cSCC application combines matched tumour and normal single-cell data with spatial sections from overlapping patients. This design distinguishes APA changes within annotated cell populations from contrasts between spatial regions that contain mixtures of cell types. The detailed procedures below explain how comparable site measurements were generated, how tumour and normal samples or tissue regions were defined, and how patient-level replication was preserved during inference. They also specify the depth, gene-composition and cellular-mixture controls behind the main biological findings.

#### cSCC cohort and sample selection

The cutaneous squamous cell carcinoma (cSCC) analysis used public single-cell data from matched tumour and normal adjacent skin of ten patients (GSE144236; 25 10x Genomics 3′ v2 libraries) [22]. We retained 47,061 cells not labelled as multiplets in the original publication and used its published cell-type and keratinocyte-subtype annotations. Spatial data from the same study comprised 16 sections from six of these patients (GSE144239, part of GSE144240): four Visium sections from patients P4 and P6 (8,928 in-tissue spots, 55-$\mu$m resolution), and 12 legacy ST sections from P2, P5, P9 and P10 (8,671 spots, approximately 110-$\mu$m resolution). Visium P4 replicate 1 and ST P2 replicate 1 also contributed to the spatial method benchmark, while the P3 normal-skin library (GSM4284230) was included only in read-structure quality control (Supplementary Fig. 2). Patient and sample identifiers are given in Supplementary Table 2.

#### cSCC alignment and filtering

The original single-cell BAM files were converted to FASTQ with Cell Ranger 9.0.0 `bamtofastq` and realigned to GRCh38/GENCODE v32 using STARsolo 2.7.11b. Spatial FASTQ files were obtained from the Sequence Read Archive and aligned with STARsolo against the same reference; the four runs representing each Visium section were merged after alignment. Published cell identities and in-tissue spot lists served as the analysis whitelists. No additional cell-type labels were inferred from APA measurements.

#### cSCC site discovery and quantification

Each data modality was processed as its own discovery cohort before barcode-level counts were compared between conditions or regions. This preserves the respective pooled evidence available in single-cell, Visium and ST libraries without presuming a common discovered-site catalogue across modalities.

The single-cell BAMs formed one scPolyA cohort, with Visium and legacy ST sections forming two separate spatial cohorts. Sample- and section-specific barcode suffixes prevented collisions. All three cohorts used revision 0b0d766 and the benchmark configuration documented above, with pooled tail-read depth threshold 3 in `terminal-aware`, `merged` mode. DeepCPS site filtering was disabled. Processing the three cohorts separately established cohort-level discovery catalogues before barcode-resolved counts were compared within patients and regions.

#### Cell identities and spatial compartments

Cell-type comparisons use the source study's labels, whereas spatial comparisons require a definition of tissue regions. The marker-based compartment procedure is reported in detail because a region is enriched for a cell population and does not represent a purified population.

Cell identities and the keratinocyte subtypes basal, cycling, differentiating and tumour-specific keratinocytes (TSK) followed Ji et al. [22]. Keratinocytes were assigned to the tumour or normal group by the source library; subtype annotations were available for seven patients. For visualisation only (Fig. 7b), keratinocytes were embedded by UMAP after Harmony correction across patients. The embedding did not determine tested groups or effect sizes.

Spatial compartments were derived from expression-marker scores rather than pathology annotations. Gene counts were normalised to 10,000 per spot and smoothed by averaging with each spot's neighbours. For each marker set, the sum of smoothed expression was scaled by its platform-specific 99th percentile and capped at 1. Each spot was assigned to the epithelial, stromal or immune compartment with the highest scaled marker abundance. Epithelial spots were labelled TSK-enriched if their TSK score, the mean $\log_2$ ratio between smoothed TSK-marker expression and its platform mean, was positive and greater than the section's epithelial median plus one standard deviation. Among remaining epithelial spots, those with differentiation score at least 0.5 and greater than the epithelial score were assigned to differentiated epithelium; the remainder were labelled tumour-enriched. These terms describe marker-defined regions and do not imply purified cell populations or independent histopathological assignments.

#### cSCC APA estimators

The length analysis was designed to compare alternative-end use while reducing changes caused solely by different expressed-gene mixtures. It uses a genomic, gene-centred quantity, and the choice of reference mean depends on whether the comparison is made within a patient, section or display cohort.

The main Methods define the gene-centred terminal-UMI-weighted genomic $3'$ UTR estimator

$R(x)$, its reference means and the 20-terminal-UMI threshold for cell- or spot-level visualisations. For the native site set, genes with fewer than two eligible terminal sites were excluded from this estimator. The reference mean for each gene was calculated over the cohort for cell displays, over the corresponding section for spatial displays and within patient for paired pseudobulks; the resulting measures were not interpreted as spliced transcript lengths.

For gene-level distal usage, one eligible terminal site with the greatest signed stop-relative distance was selected and the proportion was computed only where retained-terminal UMIs were positive. Intronic usage for the cSCC comparisons was $I/(I+T)$, using intronic and terminal UMI totals, with the resulting proportion centred against a pooled gene-specific reference. This fraction is distinct from the native log-ratio iPUI. The choices of eligible sites and reference populations were maintained within each reported contrast.

#### Patient-matched single-cell comparisons

Patient matching allows the tumour–normal contrast to be evaluated within individuals rather than treating thousands of cells as independent replicates. The rules below define which patients contribute to each lineage or subtype comparison and how gene-level and aggregate tests differ.

For patient-level tests, eligible site counts were pooled by patient, tissue condition and annotated cell type. Patients entered a contrast only if both groups contained at least ten cells. Tumour versus normal keratinocyte comparisons and comparisons with non-TSK tumour keratinocytes included ten patients; TSK, basal and cycling subsets included six, differentiating keratinocytes seven, myeloid and lymphoid controls eight, fibroblasts five, and endothelial and melanocyte controls three. Mean within-patient differences were tested with two-sided paired $t$-tests (equivalently, one-sample $t$-tests of patient differences), accompanied by 95% $t$ confidence intervals; exact Wilcoxon signed-rank tests supplied sensitivity checks.

To assess dependence on library depth and gene composition, we repeated the mean $3'$ UTR contrast with fixed weights, depth-thinned pseudobulks and depth-matched individual cells (Supplementary Fig. 13a). The fixed-weight analysis required each gene to have at least ten terminal UMIs in both groups and at least five eligible genes. It assigned each gene the mean of its normalised terminal-UMI shares across the groups and summed these fixed-weight gene differences. Depth thinning reduced the deeper group's multi-site UMIs binomially to the shallower group's total (average of five repeats). Depth matching compared cells within deciles of $\log_{10}$ multi-site UMIs, weighted by the smaller group's cell count per decile. These exploratory and primary estimates were not jointly adjusted for multiple testing.

Keratinocyte-specificity was assessed by subtracting the patient's tumour–normal myeloid change from the corresponding keratinocyte change. The resulting difference-in-differences was tested across eight eligible patients by a two-sided one-sample $t$-test. This comparison and the other Fig. 7d cell-type contrasts were adjusted together across 14 tests using Benjamini–Hochberg.

For gene-level distal-site usage, eligible genes required at least four patients with ten or more terminal UMIs in both tumour and normal conditions. For each gene, patient and condition, the transformed usage was $\log[(d+0.5)/(n-d+0.5)]$, with $d$ distal UMIs and $n$ retained terminal UMIs. Tumour-minus-normal patient differences were tested by a two-sided one-sample $t$-test and Benjamini–Hochberg adjusted within each contrast (3,852 genes for tumour versus normal keratinocytes). In a negative control, cell labels were permuted 50 times within each patient while preserving sample membership and group sizes; the largest number of significant genes and the direction of the corresponding effects were compared with the observed analysis. These permutations are an internal control for the discovery procedure, not independent biological validation.

#### Spatial region comparisons

Spatial regions include mixtures of cells, and patients may contribute more than one section. Region-level contrasts were therefore computed within sections and summarised within patients before inference, with separate analyses testing whether the spot-level signal had spatial structure beyond the chosen nulls.

As described in the main Methods, section-level differences were averaged within patient for inference. Five sections from P9 and P10 lacked eligible stromal or immune spots for the primary region comparison, and no stromal spots were identified in the P5 sections for the stroma-only analysis. Region eligibility required at least ten spots per compared group. Exact sign tests, Wilcoxon signed-rank tests and section-level summaries were sensitivity analyses; the patient remained the primary replication unit.

Spot-level spatial autocorrelation of the gene-centred $3'$ UTR measure was evaluated with upper-tail Moran's $I$ tests. The first null preserved abundance distributions (199 resamples); a second spatial-permutation null (999 permutations) was applied after regression of sequencing depth and region assignment. The procedures and eligible section populations are documented in Source Data and Supplementary Fig. 14.

#### Cell composition and cross-modal agreement

The three-term tumour–reference decomposition is defined in the main Methods. All four constituent cell populations had to contain at least ten cells per patient. This yielded six patients with a stromal reference and nine with a combined stromal-plus-immune reference. Each group used its expressed multi-site genes, and gene-reference means were shared across the decomposition's terms within each patient. A common-gene sensitivity analysis restricted all four groups to genes with at least ten terminal UMIs. Eight reported decomposition terms were tested against zero with two-sided patient-level one-sample $t$-tests, with Benjamini–Hochberg adjustment across those terms independently of the 14 cell-type contrasts.

Cross-modal effects were compared over genes evaluable in both modalities using Spearman correlation and 2,000 gene-level bootstrap samples. The expected rate of agreeing signs was calculated from the marginal fractions of genes shortening in each modality; its significance was assessed by 10,000 permutations of single-cell gene effects (one-sided). Tissue-significant genes were defined by a two-sided one-sample $t$-test of section-level differences in logit distal usage between tumour/TSK-enriched and stroma-plus-immune regions ($P < 0.01$), with at least ten terminal UMIs per region per section and four or more eligible sections from at least two patients. Concordant genes additionally required a same-direction patient-level single-cell change at $P < 0.05$. The observed number of concordant genes was compared with a separate 20,000-permutation gene-label null (one-sided; minimum $P \approx 5 \times 10^{-5}$).

Candidate genes required spatial section-level $P < 0.01$, same-direction patient-level single-cell keratinocyte $P < 0.05$, and no same-direction myeloid change at $P < 0.05$. The exploratory tier-1 set additionally required single-cell FDR $< 0.05$ and a composition-control effect not significant in the tissue direction. These criteria were applied to the analysed data and do not constitute independent confirmation. At *PDCD6IP*, tail-read support was summarised from cleavage-associated endpoints within $\pm 20$ nt of each site. Distal UMI usage used the distal count divided by all retained terminal UMIs in the corresponding patient or section. Nine patients and nine sections had sufficient proximal and distal support in both comparison groups.

### Agent execution protocols and independent evidence review

The agent case studies assess whether biological questions can be converted into executable scPolyA analyses and whether conclusions are supported by the returned molecular evidence.

Their reproducibility depends on keeping the initiating questions separate from the operational skills, preserving the configuration of each native run and checking numerical claims against its outputs. The cases therefore document not only the sequence of analytical screens but also why each screen was needed and what kind of conclusion it could support.

#### Agent setup and execution records

The two agents, three initiating researcher questions and choice of displayed executions are described in the main Methods. Versioned scPolyA-specific skills, dataset paths and operational instructions were provided separately from the biological questions. The execution evidence comprises run configurations, available operation records, native output tables and independent numerical review. No complete agent transcripts are included in the deposited Source Data.

The six completed runs used scPolyA v2.1.0. Run-specific inputs, configurations and output inventories remained separate, particularly across the two PBMC questions. Supplementary Fig. 17 records the paired agents' assessments and their independent review, including findings not selected for the main illustration.

#### Case 1: PBMC execution and APA measurements

The first case tests the transition from producing a site matrix to deciding at what cellular resolution its usage can be compared. Its shipped legacy preset is recorded explicitly because it does not produce the same native terminal-aware outputs as the principal benchmarking configuration.

The displayed Case 1 agent used the shipped scPolyA v2.1.0 preset unchanged for the 10x Genomics GEM-X 3′ v4 Human Donor1 PBMC library (GRCh38; GENCODE v32). The preset included chromosomes 1–22, X and Y, excluded chrM and used `legacy` counting. Its tail-read criteria were five clipped A/T bases, at most one other clipped base and mapping quality at least 10; discovery used pooled threshold 3 and 10-nt merging. The read-based internal-priming rule used eight A/T bases among ten aligned bases. For the subsequent eligibility review, non-intronic and intronic sites were required to pass cutoffs 0.9 and 0.7, respectively. These cutoffs were part of output interpretation and did not convert the native legacy pipeline run into terminal-aware quantification.

The delivered files included annotated sites, a site-by-cell UMI matrix, per-cell APA tables, expression-defined cell metadata and pooled site counts by group. For the separate terminal-APA review, each eligible site needed a valid gene assignment, terminal-exon or downstream context and a relevant internal-priming pass. The resulting sites belonged to 4,978 genes; the site and barcode inventories and multi-site counts are stated in the main Methods. Group-level UMI totals were unnormalised and accompanied by contributing cell numbers. Among the 5,710 barcodes, expression QC retained 4,727: 4,620 assigned to nine immune-cell groups and 107 unassigned, giving ten reported groups.

#### Case 2: discovery stringency and terminal-UMI coverage

The second case tests whether more permissive site discovery increases the number of genes with sufficiently sampled APA, an outcome distinct from the raw number of sites reported. It therefore separates site-catalogue size, multi-terminal-gene eligibility and cell-level terminal-UMI depth.

The two native PBMC analyses in Case 2 differed in pooled per-base tail-read depth threshold (3 versus 1); their remaining recorded settings and input library were matched. Tail reads comprised 598,987 of 148,886,783 deduplicated alignments (0.40%). Eligibility required a valid gene assignment, terminal-exon or downstream context and all relevant site filters; a multi-site gene had at least two distinct eligible terminal sites. Catalogue sizes and multi-site gene counts are reported in the main Methods.

The additional depth screen tested terminal UMI support within each eligible gene: at least ten terminal UMIs in each of at least 20 expression-QC-passing cells. It did not require both alternative terminal sites to carry UMIs in the same cell. The main Methods report the resulting depth-supported gene counts. A threshold-1 candidate was counted as additional if its interval had no overlap with a threshold-3 interval under the recorded comparison, a different statistic from the difference in total catalogue sizes. Of these additional candidates, 88.8% had no more than two supporting tail reads, 67.2% had intronic context and 14.2% had eligible terminal context (Fig. 8c).

#### Case 3: genomic assessment of atlas-distant neuronal sites

The third case combines evidence for an endpoint with a check on whether the reference atlas could have represented that genomic region. A large distance to the nearest mapped atlas entry has a different interpretation when the surrounding sequence was not covered by the original atlas mapping.

The third case examined scPolyA sites from a single mouse-neuron library in mm10 coordinates. The bulk reference was the same-strand PolyA_DB atlas mapped from mm9 to mm10; the sequence covered by that mapping was tracked separately from the distance to the nearest mapped site. The native catalogue contained 16,738 sites. An initial screen retained 8,525 sites detected in at least 20 expression-QC-passing neurons with assigned counts, supported by at least ten pooled tail reads and having a native internal-priming ratio of at most 0.2. For 8,435 of these, at least one neuronal tail-read endpoint lay within the site interval; the most frequent such endpoint (the mode) was used for positional assessment. Of the 8,435 sites, 441 had no same-strand PolyA_DB entry within 100 nt of the mode, and 432 of those 441 were in sequence covered by the mapped atlas.

For the stringent candidate screen, a tail molecule was defined by a distinct cell-barcode–UMI pair. Within ±10 nt of the observed endpoint mode, a candidate required at least 20 tail molecules from at least ten neurons, comprising at least 60% of its neuronal tail molecules. Further requirements were at least three supporting cells in each of two fixed halves of the neuronal population; at least 90% of supporting molecules with a single reported alignment (SAM tag `NH:i:1`); at least ten molecules with observed tails of ten or more nucleotides; an AATAAA or ATTAAA motif beginning 10–35 nt upstream; fewer than six adenosines in the first ten downstream genomic bases; and fewer than 12 in the first 20 downstream bases. Three of the 441 atlas-distant sites passed this screen. Only one, in *Bcl11a*, lay in the reference atlas's mapped sequence domain and was prioritised (Fig. 8d). The remaining two stringent candidates, in *Ppp2r3d* and *Hmgcs1*, had local RNA-end evidence but lay in mm9-derived reference regions without atlas sequence coverage.

The prioritised *Bcl11a* site was at mm10 chr11:24,168,435 on the plus strand, reported as the *one-based last aligned nucleotide* of the tail reads. Among the original alignments with endpoints within ±10 nt, 33 distinct tail molecules were detected in 33 QC-passing neurons; all 39 contributing tail-read alignments carried `NH:i:1`. The AATAAA and ATTAAA signals began 21 nt and 12 nt upstream, respectively. The first ten and first 20 downstream genomic bases each contained one adenosine. The endpoint was 1,652 nt from the nearest eligible PolyA_DB entry but 6 nt from a GENCODE transcript end, identifying a bulk-atlas catalogue gap adjacent to an annotated transcript end.

A secondary, lower-support screen retained the other criteria while lowering the local minimum to ten tail molecules, including at least five with tails of ten or more nucleotides, and restricting candidates to atlas-covered sequence. It additionally identified *AA387200*, with 11 molecules in 11 neurons, 4 nt from a GENCODE transcript end and 11,334 nt from PolyA_DB. This secondary candidate was recorded separately from the stringent screen and the prioritised *Bcl11a* example.

#### Independent verification and reporting scope

Independent numerical review recalculated each execution's reported findings directly from its own native output files. Case-specific checks covered the input and output inventories, site and barcode counts, terminal-site eligibility, depth thresholds, additional-site denominators, strand-aware atlas distances and the read- and sequence-level neuronal candidate screen. Verification considered all six executions; the representative panels in Fig. 8 were chosen after that review, and the other agent's results remain documented in Supplementary Fig. 17. Source Data provides the numerical review tables without complete session transcripts.

The two PBMC questions were run separately even though they used the same donor library; their distinct 9,030- and 9,047-site native catalogues were not pooled or treated as repeated measurements of one run. The neuronal question was evaluated in one mouse-neuron library. These are workflow and evidence-review case studies, not estimates of biological replication across donors or neuronal datasets.

### Statistical reporting and software

Different figures use sites, genes, cells, sections or patients as their observational units. The reporting conventions below are interpreted together with the analysis-specific eligibility rules above, so that a resampling interval or adjusted significance symbol can be traced to its appropriate test family and replicate definition.

#### Statistical analysis and reproducibility

The main Methods specify the general inferential conventions and independent units for the principal tests. Figures use $***$ for $P < 0.001$, $**$ for $P < 0.01$, $*$ for $P < 0.05$ and ns for non-significant results; adjusted significance uses the specified multiple-testing family. The null intervals in Supplementary Fig. 13b are mean $\pm 1.96$ standard deviations, and numerical values below $10^{-16}$ are reported as $P < 10^{-16}$. Empirical permutation probabilities used $(k+1)/(B+1)$ unless a section above states a different construction.

Dataset-specific QC, unsuccessful method executions, eligibility and replicate definitions are documented above and in Supplementary Tables 2–4. For the spatial method comparison, independently analysed sections supplied the dataset-level units ($n = 7$–$8$), while multiple sections from the same cSCC patient were collapsed to patient-level effects before biological inference. The code used Python 3.10, NumPy 2.2, SciPy 1.15 and statsmodels 0.14 for statistical and plotting analyses; DeepCPS training used PyTorch 2.11 and scikit-learn 1.7. These package versions supplement the versioned method settings and execution boundaries in Supplementary Table 3.

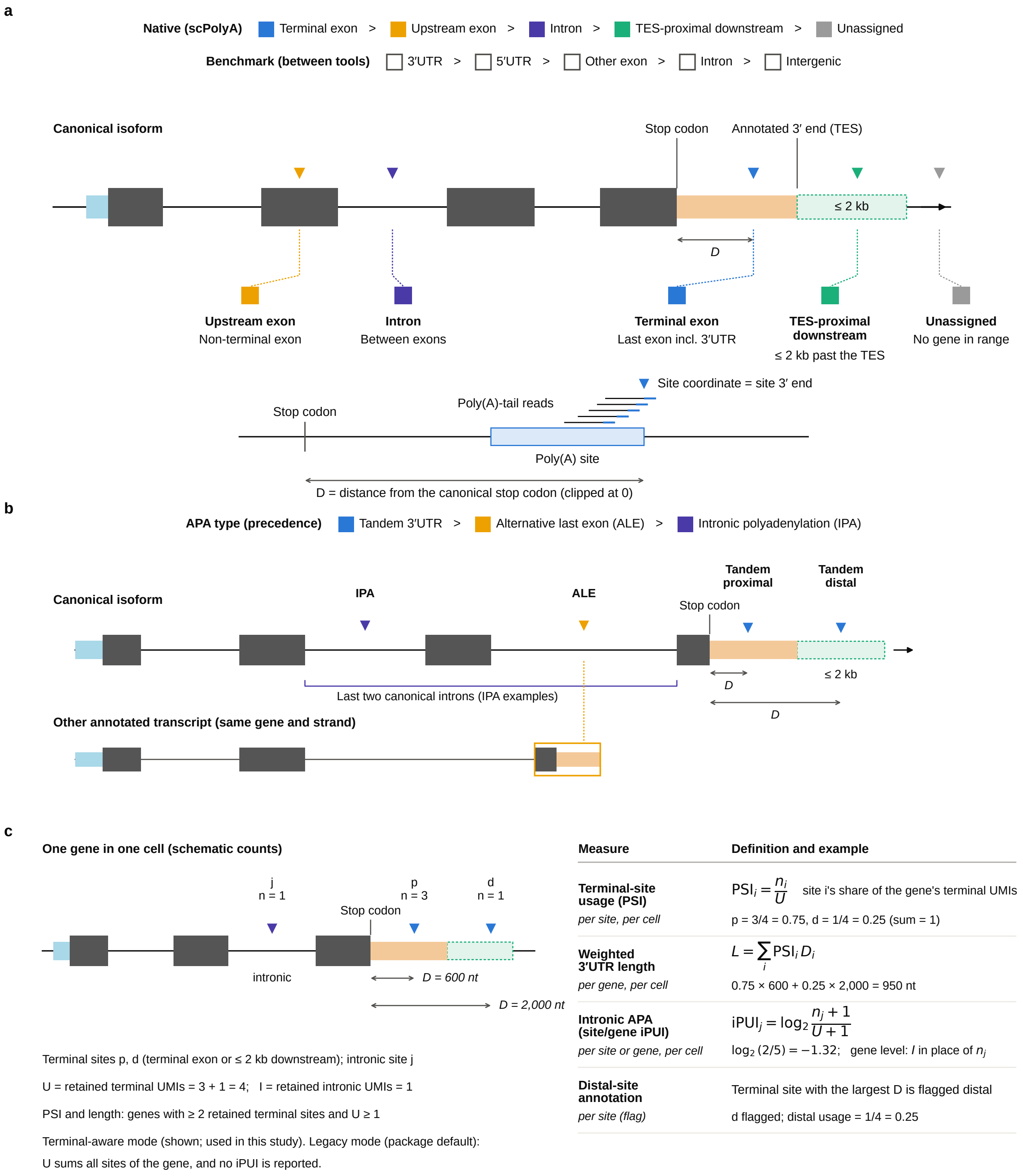
a
Native (scPolyA) Terminal exon > Upstream exon > Intron > TES-proximal downstream > Unassigned
Benchmark (between tools) 3′UTR > 5′UTR > Other exon > Intron > Intergenic
Canonical isoform
Stop codon
Annotated 3′ end (TES)
≤ 2 kb
D
Upstream exon
Non-terminal exon
Intron
Between exons
Terminal exon
Last exon incl. 3′UTR
TES-proximal downstream
≤ 2 kb past the TES
Unassigned
No gene in range
Site coordinate = site 3′ end
Poly(A)-tail reads
Stop codon
Poly(A) site
D = distance from the canonical stop codon (clipped at 0)
b
APA type (precedence) Tandem 3′UTR > Alternative last exon (ALE) > Intronic polyadenylation (IPA)
Tandem proximal
Tandem distal
Canonical isoform
IPA
ALE
Stop codon
≤ 2 kb
D
Last two canonical introns (IPA examples)
D
Other annotated transcript (same gene and strand)
c
One gene in one cell (schematic counts)
j
n = 1
p
n = 3
d
n = 1
Stop codon
intronic
D = 600 nt
D = 2,000 nt
Terminal sites p, d (terminal exon or ≤ 2 kb downstream); intronic site j
U = retained terminal UMIs = 3 + 1 = 4; I = retained intronic UMIs = 1
PSI and length: genes with ≥ 2 retained terminal sites and U ≥ 1
Terminal-aware mode (shown; used in this study). Legacy mode (package default):
U sums all sites of the gene, and no iPUI is reported.
Measure | Definition and example
Terminal-site usage (PSI)
per site, per cell
$\mathrm{PSI}_i = \frac{n_i}{U}$ site i's share of the gene's terminal UMIs
p = 3/4 = 0.75, d = 1/4 = 0.25 (sum = 1)
Weighted 3′UTR length
per gene, per cell
$L = \sum_i \mathrm{PSI}_i D_i$
0.75 × 600 + 0.25 × 2,000 = 950 nt
Intronic APA (site/gene iPUI)
per site or gene, per cell
$\mathrm{iPUI}_j = \log_2 \frac{n_j + 1}{U + 1}$
$\log_2(2/5) = -1.32$; gene level: $I$ in place of $n_j$
Distal-site annotation
per site (flag)
Terminal site with the largest D is flagged distal
d flagged; distal usage = 1/4 = 0.25

**Supplementary Figure 1 | Site annotation links candidate RNA ends to interpretable forms of APA.**

A genomic endpoint becomes biologically informative when it can be related to a gene and its transcript structure. scPolyA first assigns candidates to canonical-transcript contexts and then distinguishes the alternative-end choices that can underlie a change in RNA processing.

**a,** Sites are assigned in priority order to the canonical terminal exon, an upstream exon, an intron or a region extending downstream of the annotated transcript end; sites outside these categories remain unassigned (labelled Intergenic in native output). Colours indicate the annotation classes and open boxes the contexts used in cross-method comparisons. The strand-specific site endpoint and its genomic distance $D$ from the canonical stop codon define the reported position and $3'$ UTR distance. CDS, coding sequence; TES, transcript end.

**b,** Alternative sites are classified as tandem $3'$ UTR, alternative last-exon (ALE) or intronic polyadenylation (IPA) events by comparing the canonical transcript with other annotated isoforms of the same gene. The schematic shows the classification precedence and the terminal canonical introns considered for the illustrative IPA loci.

**c,** Site-specific UMIs are summarised as within-gene APA measures, illustrated for one gene in one cell. Terminal-site usage (PSI) gives each terminal site's share of the gene's terminal UMIs, usage-weighted $3'$ UTR length combines these shares with site distances, and site- and gene-level iPUI express intronic relative to terminal UMI support as log-ratios with pseudocounts; the farthest terminal site is flagged as distal. Values follow the terminal-aware mode used in this study.

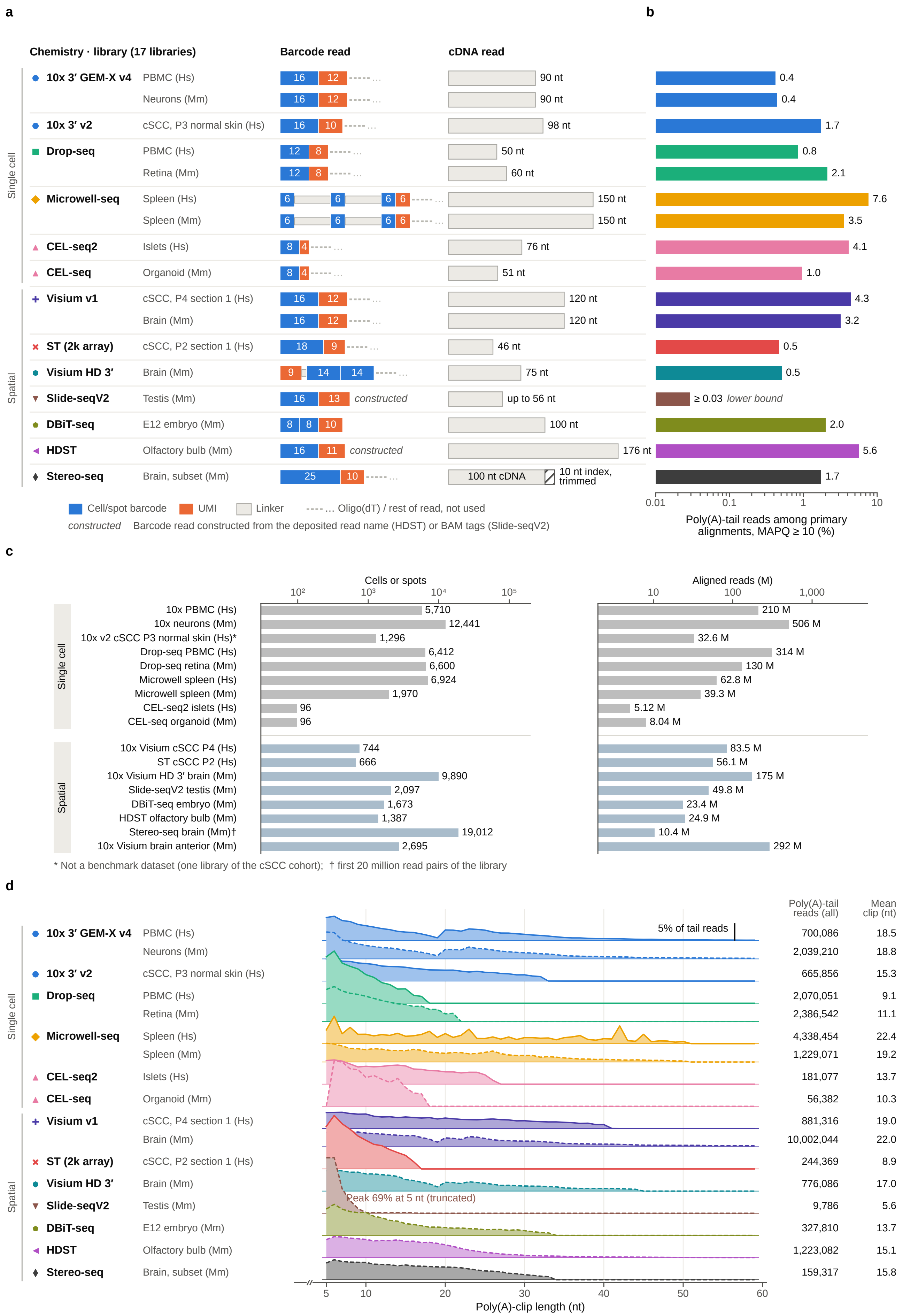
a
b
Chemistry · library (17 libraries)
Barcode read
cDNA read
Single cell
Spatial
10x 3′ GEM-X v4
PBMC (Hs)
Neurons (Mm)
10x 3′ v2
cSCC, P3 normal skin (Hs)
Drop-seq
PBMC (Hs)
Retina (Mm)
Microwell-seq
Spleen (Hs)
Spleen (Mm)
CEL-seq2
Islets (Hs)
CEL-seq
Organoid (Mm)
Visium v1
cSCC, P4 section 1 (Hs)
Brain (Mm)
ST (2k array)
cSCC, P2 section 1 (Hs)
Visium HD 3′
Brain (Mm)
Slide-seqV2
Testis (Mm)
DBiT-seq
E12 embryo (Mm)
HDST
Olfactory bulb (Mm)
Stereo-seq
Brain, subset (Mm)
16 12 90 nt 0.4
16 12 90 nt 0.4
16 10 98 nt 1.7
12 8 50 nt 0.8
12 8 60 nt 2.1
6 6 6 6 150 nt 7.6
6 6 6 6 150 nt 3.5
8 4 76 nt 4.1
8 4 51 nt 1.0
16 12 120 nt 4.3
16 12 120 nt 3.2
18 9 46 nt 0.5
9 14 14 75 nt 0.5
16 13 constructed up to 56 nt ≥ 0.03 lower bound
8 8 10 100 nt 2.0
16 11 constructed 176 nt 5.6
25 10 100 nt cDNA 10 nt index, trimmed 1.7
Cell/spot barcode
UMI
Linker
---- … Oligo(dT) / rest of read, not used
constructed Barcode read constructed from the deposited read name (HDST) or BAM tags (Slide-seqV2)
0.01 0.1 1 10
Poly(A)-tail reads among primary alignments, MAPQ ≥ 10 (%)
c
Cells or spots
$10^2$ $10^3$ $10^4$ $10^5$
Aligned reads (M)
10 100 1,000
10x PBMC (Hs) 5,710 210 M
10x neurons (Mm) 12,441 506 M
10x v2 cSCC P3 normal skin (Hs)* 1,296 32.6 M
Drop-seq PBMC (Hs) 6,412 314 M
Drop-seq retina (Mm) 6,600 130 M
Microwell spleen (Hs) 6,924 62.8 M
Microwell spleen (Mm) 1,970 39.3 M
CEL-seq2 islets (Hs) 96 5.12 M
CEL-seq organoid (Mm) 96 8.04 M
10x Visium cSCC P4 (Hs) 744 83.5 M
ST cSCC P2 (Hs) 666 56.1 M
10x Visium HD 3′ brain (Mm) 9,890 175 M
Slide-seqV2 testis (Mm) 2,097 49.8 M
DBiT-seq embryo (Mm) 1,673 23.4 M
HDST olfactory bulb (Mm) 1,387 24.9 M
Stereo-seq brain (Mm)† 19,012 10.4 M
10x Visium brain anterior (Mm) 2,695 292 M
* Not a benchmark dataset (one library of the cSCC cohort); † first 20 million read pairs of the library
d
Poly(A)-tail reads (all)
Mean clip (nt)
5% of tail reads
10x 3′ GEM-X v4 PBMC (Hs) 700,086 18.5
Neurons (Mm) 2,039,210 18.8
10x 3′ v2 cSCC, P3 normal skin (Hs) 665,856 15.3
Drop-seq PBMC (Hs) 2,070,051 9.1
Retina (Mm) 2,386,542 11.1
Microwell-seq Spleen (Hs) 4,338,454 22.4
Spleen (Mm) 1,229,071 19.2
CEL-seq2 Islets (Hs) 181,077 13.7
CEL-seq Organoid (Mm) 56,382 10.3
Visium v1 cSCC, P4 section 1 (Hs) 881,316 19.0
Brain (Mm) 10,002,044 22.0
ST (2k array) cSCC, P2 section 1 (Hs) 244,369 8.9
Visium HD 3′ Brain (Mm) 776,086 17.0
Slide-seqV2 Testis (Mm) 9,786 5.6
Peak 69% at 5 nt (truncated)
DBiT-seq E12 embryo (Mm) 327,810 13.7
HDST Olfactory bulb (Mm) 1,223,082 15.1
Stereo-seq Brain, subset (Mm) 159,317 15.8
5 10 20 30 40 50 60
Poly(A)-clip length (nt)

**Supplementary Figure 2 | Library architecture shapes the direct RNA-end evidence available for APA analysis.**

Oligo(dT)-primed libraries capture RNA ends through different read arrangements, and only a subset of their alignments retains a recognisable poly(A) tail. These differences help explain why the same discovery threshold can expose different sets of candidate sites across single-cell and spatial protocols.

**a,** Read layouts are compared across 17 libraries (eight single-cell benchmark libraries, one additional cSCC P3 normal-skin library and eight spatial sections), showing where cell or spatial barcodes, UMIs, linkers and cDNA sequence occur. Constructed barcode reads for HDST and Slide-seqV2 were recovered from deposited identifiers; Stereo-seq's terminal 10-nt sample index was removed before alignment. The drawn Slide-seqV2 cDNA length reflects its deposited reads. Dataset-specific preprocessing is detailed in Supplementary Table 2.

**b,** The fraction of primary alignments containing identifiable poly(A)-tail evidence varies markedly among libraries (mapping quality $\geq 10$; logarithmic scale). The Slide-seqV2 estimate reflects the tail evidence retained in its deposited reads.

**c,** Dataset sizes span different numbers of cells or spatial units and aligned reads (logarithmic axes). The asterisk denotes the additional cSCC P3 normal-skin library used for read-structure assessment; the dagger denotes the Stereo-seq subset comprising the first 20 million read pairs.

**d,** Poly(A)-containing soft clips also differ in length across protocols. Curves show each library's clip-length distribution from 5 to 60 nt (solid, human; dashed, mouse), with the total tail-read count and mean clip length given at right. The scale bar denotes 5% of that library's extracted tail reads; longer clips were rare.

**a**

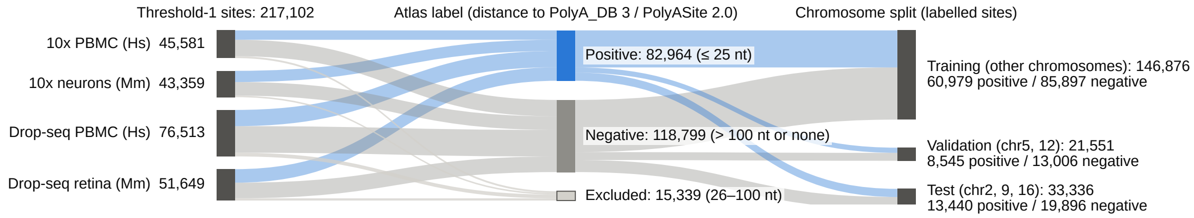

**b**

| | | Dataset | | | | Poly(A)-tail reads per site | | | | | Genomic context | | | | |
|---|---|---|---|---|---|---|---|---|---|---|---|---|---|---|---|
| | | 10x PBMC (Hs) | 10x neurons (Mm) | Drop-seq PBMC (Hs) | Drop-seq retina (Mm) | 1 | 2 | 3–5 | 6–10 | > 10 | Terminal exon | Down-stream | Other exon | Intron | Inter-genic |
| AUROC | **DeepCPS scoring** ● | **0.947** | **0.956** | **0.925** | **0.933** | **0.898** | **0.908** | **0.924** | **0.944** | **0.959** | **0.948** | **0.948** | **0.741** | **0.886** | **0.912** |
| | DeepPASS† ◆ | 0.927 | 0.930 | 0.897 | 0.898 | 0.863 | 0.871 | 0.892 | 0.917 | 0.943 | 0.940 | 0.938 | 0.680 | 0.852 | 0.885 |
| | Sequence rule ▲ | 0.889 | 0.890 | 0.813 | 0.847 | 0.783 | 0.817 | 0.839 | 0.872 | 0.907 | 0.890 | 0.884 | 0.578 | 0.765 | 0.828 |
| | DeepPASTA ■ | 0.874 | 0.881 | 0.775 | 0.805 | 0.766 | 0.792 | 0.809 | 0.848 | 0.858 | 0.865 | 0.863 | 0.658 | 0.772 | 0.790 |
| | Poly(A)-tail-read count ▼ | 0.720 | 0.813 | 0.738 | 0.745 | – | – | – | – | – | 0.726 | 0.725 | 0.635 | 0.646 | 0.666 |
| | Internal-priming score (−) × | 0.563 | 0.583 | 0.648 | 0.707 | 0.584 | 0.628 | 0.684 | 0.744 | 0.801 | 0.656 | 0.654 | 0.511 | 0.563 | 0.618 |
| | Prevalence (random AUPRC) | 0.35 | 0.43 | 0.37 | 0.46 | 0.20 | 0.29 | 0.43 | 0.60 | 0.83 | 0.86 | 0.74 | 0.25 | 0.16 | 0.30 |
| AUPRC | **DeepCPS scoring** ● | **0.935** | **0.957** | **0.910** | **0.938** | **0.785** | **0.862** | **0.923** | **0.968** | **0.991** | **0.991** | **0.983** | **0.592** | **0.729** | **0.859** |
| | DeepPASS† ◆ | 0.905 | 0.930 | 0.870 | 0.901 | 0.707 | 0.782 | 0.876 | 0.946 | 0.988 | 0.989 | 0.980 | 0.523 | 0.602 | 0.796 |
| | Sequence rule ▲ | 0.856 | 0.894 | 0.788 | 0.857 | 0.604 | 0.739 | 0.833 | 0.922 | 0.980 | 0.980 | 0.960 | 0.423 | 0.547 | 0.733 |
| | DeepPASTA ■ | 0.841 | 0.876 | 0.715 | 0.801 | 0.548 | 0.674 | 0.787 | 0.901 | 0.967 | 0.974 | 0.948 | 0.490 | 0.505 | 0.651 |
| | Poly(A)-tail-read count ▼ | 0.608 | 0.791 | 0.652 | 0.720 | – | – | – | – | – | 0.932 | 0.873 | 0.374 | 0.293 | 0.465 |
| | Internal-priming score (−) × | 0.380 | 0.469 | 0.453 | 0.602 | 0.237 | 0.365 | 0.570 | 0.784 | 0.940 | 0.896 | 0.804 | 0.252 | 0.174 | 0.355 |
| | Positive / negative | 2,354 / 4,373 | 3,176 / 4,240 | 4,083 / 6,809 | 3,827 / 4,474 | 2,639 / 10,313 | 1,693 / 4,095 | 2,458 / 3,292 | 1,890 / 1,241 | 4,760 / 955 | 8,746 / 1,475 | 433 / 150 | 309 / 910 | 2,651 / 14,274 | 1,301 / 3,087 |

0.0 0.2 0.4 0.6 0.8 1.0
Lift over random ranking (0 = random, 1 = perfect)

**c**

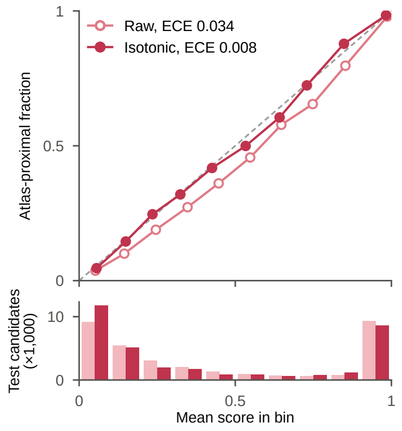

**Supplementary Figure 3 | DeepCPS distinguishes reference-proximal RNA ends across libraries and read-support levels.**

Candidate-site assessment must remain informative when tail reads are sparse and when the genomic context differs between libraries. DeepCPS was evaluated using sites generated by scPolyA itself, with chromosomes separated for training, model selection and testing; calibrated scores additionally make atlas proximity interpretable on a probability scale.

**a,** The training and evaluation cohort begins with 217,102 threshold-1 candidates from four droplet datasets. Candidates within 25 nt of PolyA_DB 3 or PolyASite 2.0 were labelled positive, those more than 100 nt away negative, and intermediate candidates excluded. Chromosomes 2, 9 and 16 were held out for testing and chromosomes 5 and 12 for validation; ribbon widths follow candidate counts.

**b,** Across individual datasets, tail-read-count strata and genomic contexts, DeepCPS is compared with DeepPASS, DeepPASTA, a fitted sequence rule, tail-read count and internal-priming score. The upper and lower heat maps show AUROC and AUPRC, with colours representing performance above random ranking; sample sizes and positive-class frequencies appear below. Hatched entries indicate that read count is constant within its own stratum. The dagger marks chromosomes represented in the DeepPASS training atlas.

**c,** Calibration on validation chromosomes brings predicted scores into closer agreement with the observed frequency of atlas proximity on test chromosomes. The reliability plot contrasts raw and isotonic-calibrated estimates and shows the candidates in each probability bin; expected calibration error (ECE) decreases from 0.034 to 0.008. The released calibrated score estimates atlas proximity, while the site-selection thresholds in Supplementary Fig. 6 use raw scores.

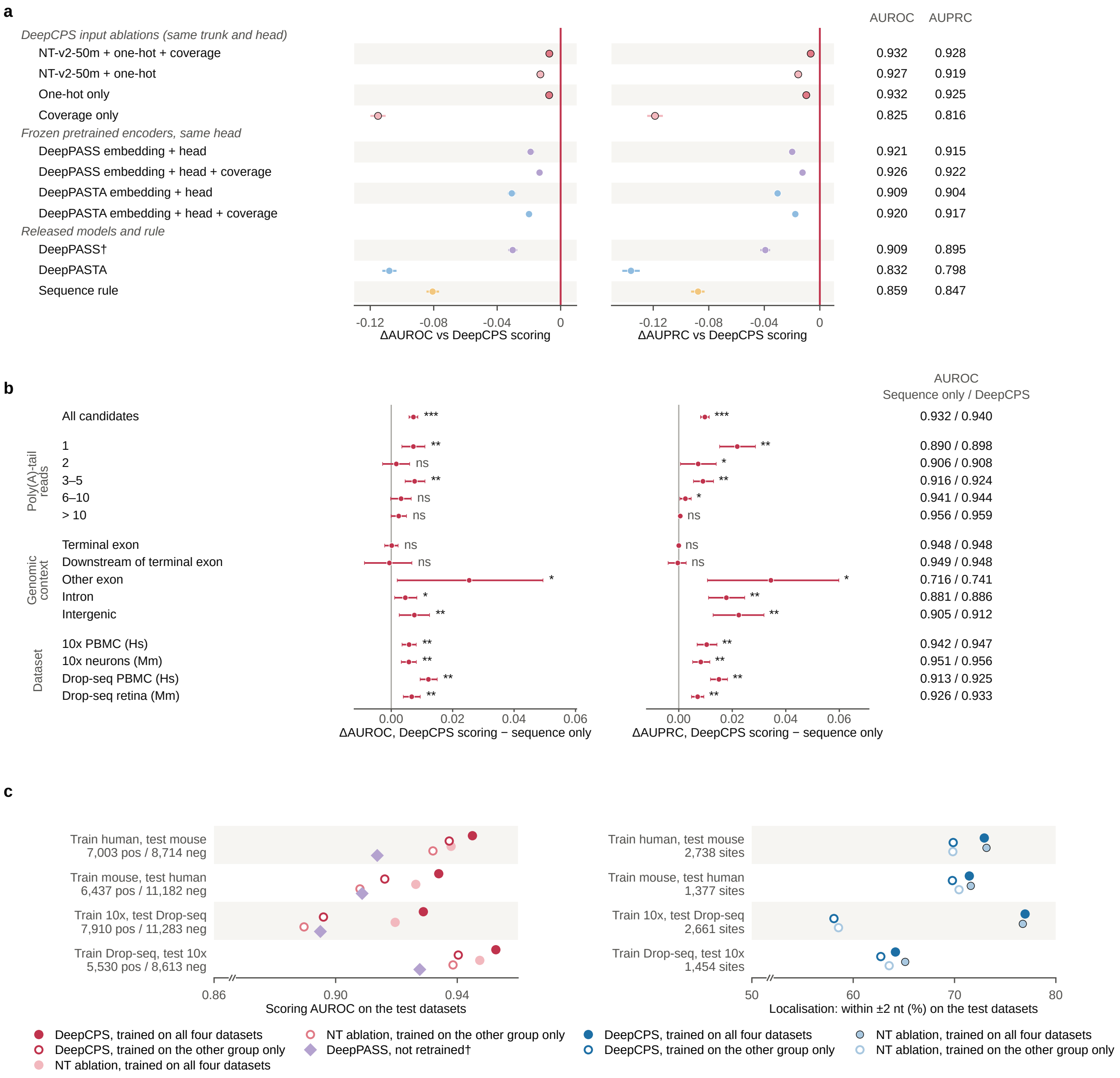

a
AUROC AUPRC
DeepCPS input ablations (same trunk and head)
NT-v2-50m + one-hot + coverage 0.932 0.928
NT-v2-50m + one-hot 0.927 0.919
One-hot only 0.932 0.925
Coverage only 0.825 0.816
Frozen pretrained encoders, same head
DeepPASS embedding + head 0.921 0.915
DeepPASS embedding + head + coverage 0.926 0.922
DeepPASTA embedding + head 0.909 0.904
DeepPASTA embedding + head + coverage 0.920 0.917
Released models and rule
DeepPASS† 0.909 0.895
DeepPASTA 0.832 0.798
Sequence rule 0.859 0.847
-0.12 -0.08 -0.04 0
ΔAUROC vs DeepCPS scoring
ΔAUPRC vs DeepCPS scoring
b
AUROC
Sequence only / DeepCPS
All candidates *** *** 0.932 / 0.940
Poly(A)-tail reads
1 ** ** 0.890 / 0.898
2 ns * 0.906 / 0.908
3–5 ** ** 0.916 / 0.924
6–10 ns * 0.941 / 0.944
> 10 ns ns 0.956 / 0.959
Genomic context
Terminal exon ns ns 0.948 / 0.948
Downstream of terminal exon ns ns 0.949 / 0.948
Other exon * * 0.716 / 0.741
Intron * ** 0.881 / 0.886
Intergenic ** ** 0.905 / 0.912
Dataset
10x PBMC (Hs) ** ** 0.942 / 0.947
10x neurons (Mm) ** ** 0.951 / 0.956
Drop-seq PBMC (Hs) ** ** 0.913 / 0.925
Drop-seq retina (Mm) ** ** 0.926 / 0.933
0.00 0.02 0.04 0.06
ΔAUROC, DeepCPS scoring − sequence only
ΔAUPRC, DeepCPS scoring − sequence only
c
Train human, test mouse
7,003 pos / 8,714 neg
Train mouse, test human
6,437 pos / 11,182 neg
Train 10x, test Drop-seq
7,910 pos / 11,283 neg
Train Drop-seq, test 10x
5,530 pos / 8,613 neg
0.86 0.90 0.94
Scoring AUROC on the test datasets
Train human, test mouse
2,738 sites
Train mouse, test human
1,377 sites
Train 10x, test Drop-seq
2,661 sites
Train Drop-seq, test 10x
1,454 sites
50 60 70 80
Localisation: within ±2 nt (%) on the test datasets
DeepCPS, trained on all four datasets
DeepCPS, trained on the other group only
NT ablation, trained on all four datasets
NT ablation, trained on the other group only
DeepPASS, not retrained†
DeepCPS, trained on all four datasets
DeepCPS, trained on the other group only
NT ablation, trained on all four datasets
NT ablation, trained on the other group only

**Supplementary Figure 4 | Sequence provides the principal DeepCPS recognition signal, with added information from read coverage.**

DeepCPS combines local sequence features with the coverage pattern observed in each library. Input ablations reveal what each source contributes, while transfer analyses test how the resulting predictions behave across species and sequencing chemistries.

**a,** Matched model variants are compared on 33,336 held-out candidates. The scoring network is evaluated with sequence alone, coverage alone, frozen Nucleotide Transformer v2 representations and released DeepPASS or DeepPASTA representations, alongside the original models and a sequence rule. Points show differences from the full DeepCPS model in AUROC and AUPRC, with 95% locus-cluster bootstrap intervals; absolute scores are given at right.

**b,** The contribution of coverage beyond sequence is examined overall and separately by tail-read support, genomic context and dataset. Differences between the full and sequence-only models are shown with bootstrap intervals and adjusted significance symbols; the absolute AUROCs of both models appear at right. The full model reaches AUROC 0.940 compared with 0.932 for sequence alone.

**c,** Models trained on one species or droplet protocol are evaluated on the other, using held-out chromosomes. Scoring AUROC (left) and localisation within 2 nt of the observed read-end mode (right) are compared with the model trained on all four datasets, Nucleotide Transformer variants and released DeepPASS where applicable. Open symbols identify training that excludes the evaluated group; filled symbols use the combined training cohort. The dagger identifies the DeepPASS atlas overlap and // an axis break.

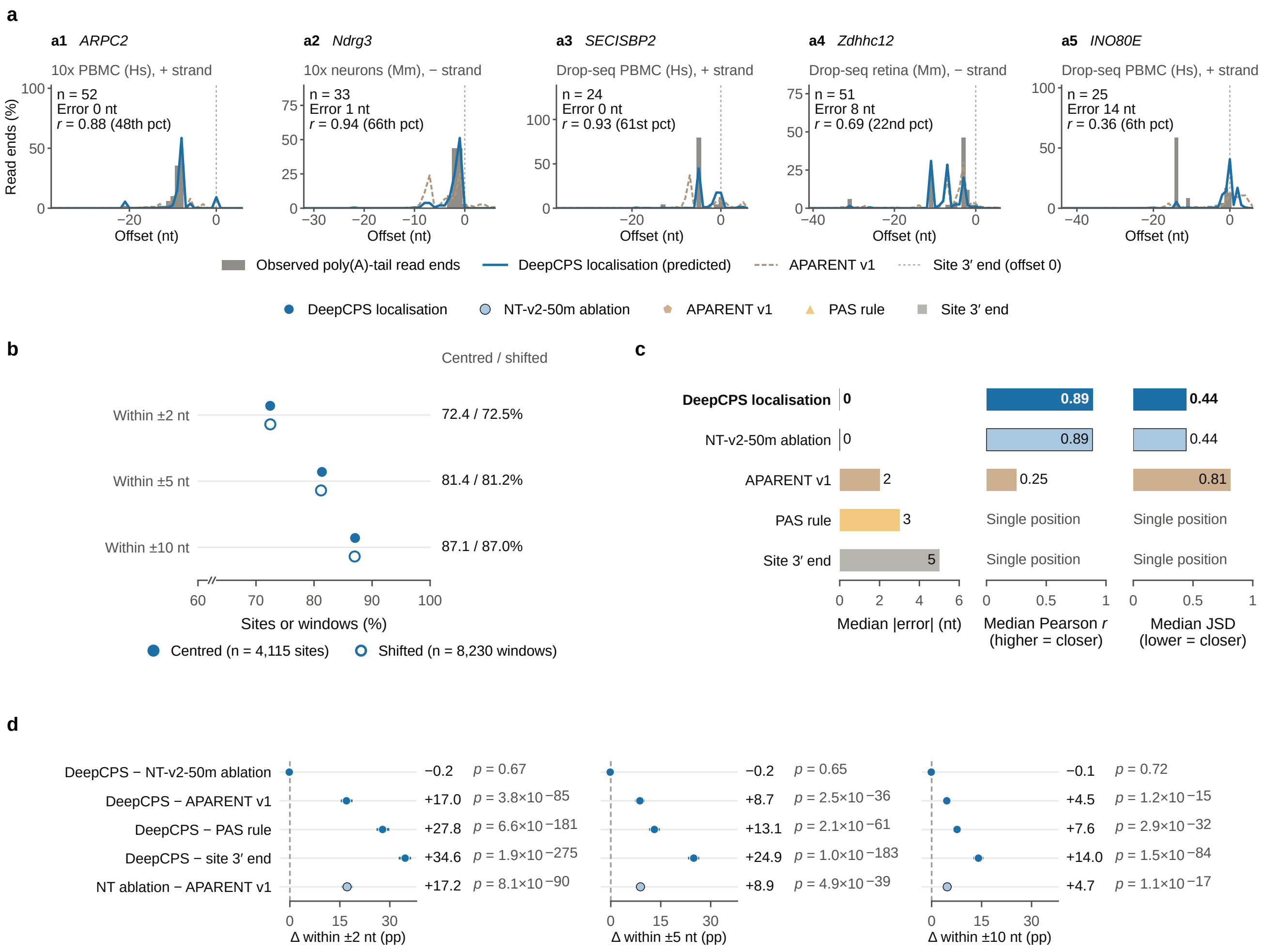
a
a1 ARPC2
10x PBMC (Hs), + strand
n = 52
Error 0 nt
r = 0.88 (48th pct)
a2 Ndrg3
10x neurons (Mm), − strand
n = 33
Error 1 nt
r = 0.94 (66th pct)
a3 SECISBP2
Drop-seq PBMC (Hs), + strand
n = 24
Error 0 nt
r = 0.93 (61st pct)
a4 Zdhhc12
Drop-seq retina (Mm), − strand
n = 51
Error 8 nt
r = 0.69 (22nd pct)
a5 INO80E
Drop-seq PBMC (Hs), + strand
n = 25
Error 14 nt
r = 0.36 (6th pct)
Read ends (%)
Offset (nt)
Observed poly(A)-tail read ends
DeepCPS localisation (predicted)
APARENT v1
Site 3′ end (offset 0)
DeepCPS localisation
NT-v2-50m ablation
APARENT v1
PAS rule
Site 3′ end
b
Centred / shifted
Within ±2 nt
72.4 / 72.5%
Within ±5 nt
81.4 / 81.2%
Within ±10 nt
87.1 / 87.0%
Sites or windows (%)
Centred (n = 4,115 sites)
Shifted (n = 8,230 windows)
c
DeepCPS localisation
NT-v2-50m ablation
APARENT v1
PAS rule
Site 3′ end
0
0
2
3
5
0.89
0.89
0.25
Single position
Single position
0.44
0.44
0.81
Single position
Single position
Median |error| (nt)
Median Pearson r (higher = closer)
Median JSD (lower = closer)
d
DeepCPS − NT-v2-50m ablation
DeepCPS − APARENT v1
DeepCPS − PAS rule
DeepCPS − site 3′ end
NT ablation − APARENT v1
−0.2 p = 0.67
+17.0 p = 3.8×10−85
+27.8 p = 6.6×10−181
+34.6 p = 1.9×10−275
+17.2 p = 8.1×10−90
Δ within ±2 nt (pp)
−0.2 p = 0.65
+8.7 p = 2.5×10−36
+13.1 p = 2.1×10−61
+24.9 p = 1.0×10−183
+8.9 p = 4.9×10−39
Δ within ±5 nt (pp)
−0.1 p = 0.72
+4.5 p = 1.2×10−15
+7.6 p = 2.9×10−32
+14.0 p = 1.5×10−84
+4.7 p = 1.1×10−17
Δ within ±10 nt (pp)

**Supplementary Figure 5 | DeepCPS localises cleavage-associated read ends with nucleotide-level precision.**

Predicting the position of an RNA end is a separate task from deciding whether a candidate resembles an established poly(A) site. DeepCPS learns positional distributions from observed tail-read endpoints, and its localisation was examined at 4,115 atlas-supported test sites with at least ten tail reads.

**a,** Five loci illustrate how the predicted endpoint distribution compares with the observed tail-read ends, including both close matches and larger positional differences. Grey bars show observed endpoints, blue the DeepCPS distribution, dashed lines APARENT v1 and dotted lines the original scPolyA boundary. Labels give tail-read count, modal-position error and the correlation between predicted and observed profiles.

**b,** Endpoint predictions remain similar when the input window is shifted by as much as 40 nt. Curves compare the percentage of sites localised within 2, 5 or 10 nt using centred and shifted windows (two shifts per site).

**c,** Positional and profile-level agreement are summarised by median absolute modal error, Pearson correlation and Jensen–Shannon divergence. The median DeepCPS modal error is zero nucleotides, indicating exact modal agreement for more than half of test sites; position-only rules do not generate profiles.

**d,** Paired improvements in localisation within 2, 5 and 10 nt are shown with 95% locus-cluster bootstrap intervals and exact McNemar tests. The Nucleotide Transformer input variant performs similarly to the selected DeepCPS architecture.

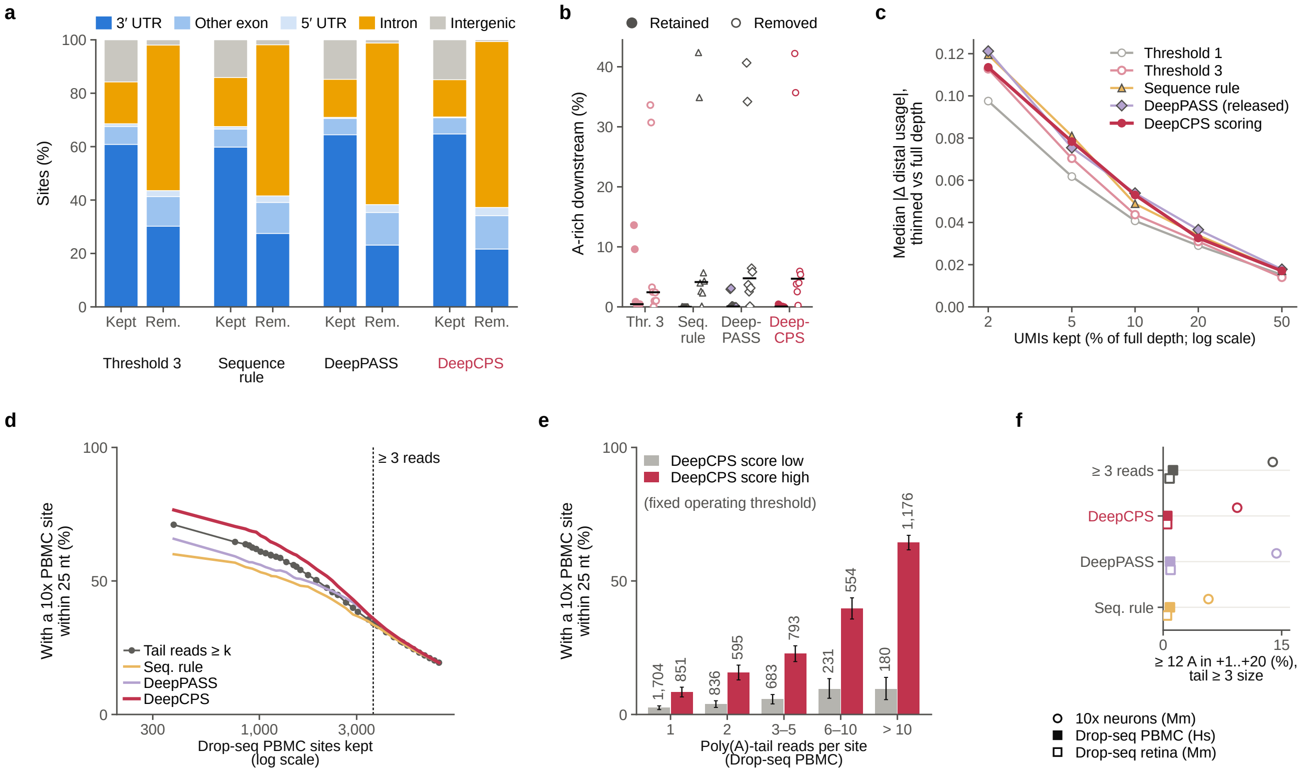
a
3′ UTR
Other exon
5′ UTR
Intron
Intergenic
Sites (%)
Kept
Rem.
Threshold 3
Sequence rule
DeepPASS
DeepCPS
b
Retained
Removed
A-rich downstream (%)
Thr. 3
Seq. rule
Deep-PASS
Deep-CPS
c
Median |Δ distal usage|, thinned vs full depth
Threshold 1
Threshold 3
Sequence rule
DeepPASS (released)
DeepCPS scoring
UMIs kept (% of full depth; log scale)
d
With a 10x PBMC site within 25 nt (%)
≥ 3 reads
Tail reads ≥ k
Seq. rule
DeepPASS
DeepCPS
Drop-seq PBMC sites kept (log scale)
e
With a 10x PBMC site within 25 nt (%)
DeepCPS score low
DeepCPS score high
(fixed operating threshold)
1,704
851
836
595
683
793
231
554
180
1,176
Poly(A)-tail reads per site (Drop-seq PBMC)
1
2
3–5
6–10
> 10
f
≥ 3 reads
DeepCPS
DeepPASS
Seq. rule
≥ 12 A in +1..+20 (%), tail ≥ 3 size
10x neurons (Mm)
Drop-seq PBMC (Hs)
Drop-seq retina (Mm)

**Supplementary Figure 6 | DeepCPS site selection improves catalogue evidence while usage precision remains linked to molecular depth.**

Candidate filtering changes which RNA ends enter the catalogue, but the precision of their usage estimates also depends on how many UMIs are observed. Selection strategies are compared at matched catalogue size, followed by a UMI-downsampling analysis of distal-site usage and cross-dataset tests of PBMC site reproducibility. For panels a–c, score thresholds were chosen on validation chromosomes and applied unchanged to eight test datasets.

**a,** Retained and excluded candidates are compared by genomic context, showing which classes are preferentially removed by each size-matched selection rule. The eight test sets are pooled.

**b,** DeepCPS selection reduces the prevalence of downstream A-rich sites among retained candidates. Filled and open symbols denote kept and removed sites, respectively; points are datasets and bars medians. A-richness is defined as at least 12 adenosines in the 20 nt downstream.

**c,** Distal-usage estimates after binomial downsampling of site-specific UMI counts are compared with their full-depth values for the same eligible genes. Median absolute errors remain similar across filtering strategies, indicating that usage precision depends primarily on the available site-specific molecules.

**d,** Independent site recurrence is evaluated in the reverse PBMC direction to Fig. 2h: Drop-seq candidates are ranked and checked for a matching 10x PBMC site within 25 nt. Curves follow recurrence as the retained catalogue grows; the dashed line marks the tail-read-threshold-3 size.

**e,** Within each Drop-seq tail-read-support stratum, candidates above the validation-selected DeepCPS score threshold are more frequently detected in 10x PBMC than lower-scoring candidates. Bars show 95% bootstrap intervals and numbers the candidate counts.

**f,** The downstream A-rich fraction is also compared at matched site count for three additional datasets, extending the evidence beyond the primary PBMC selection.

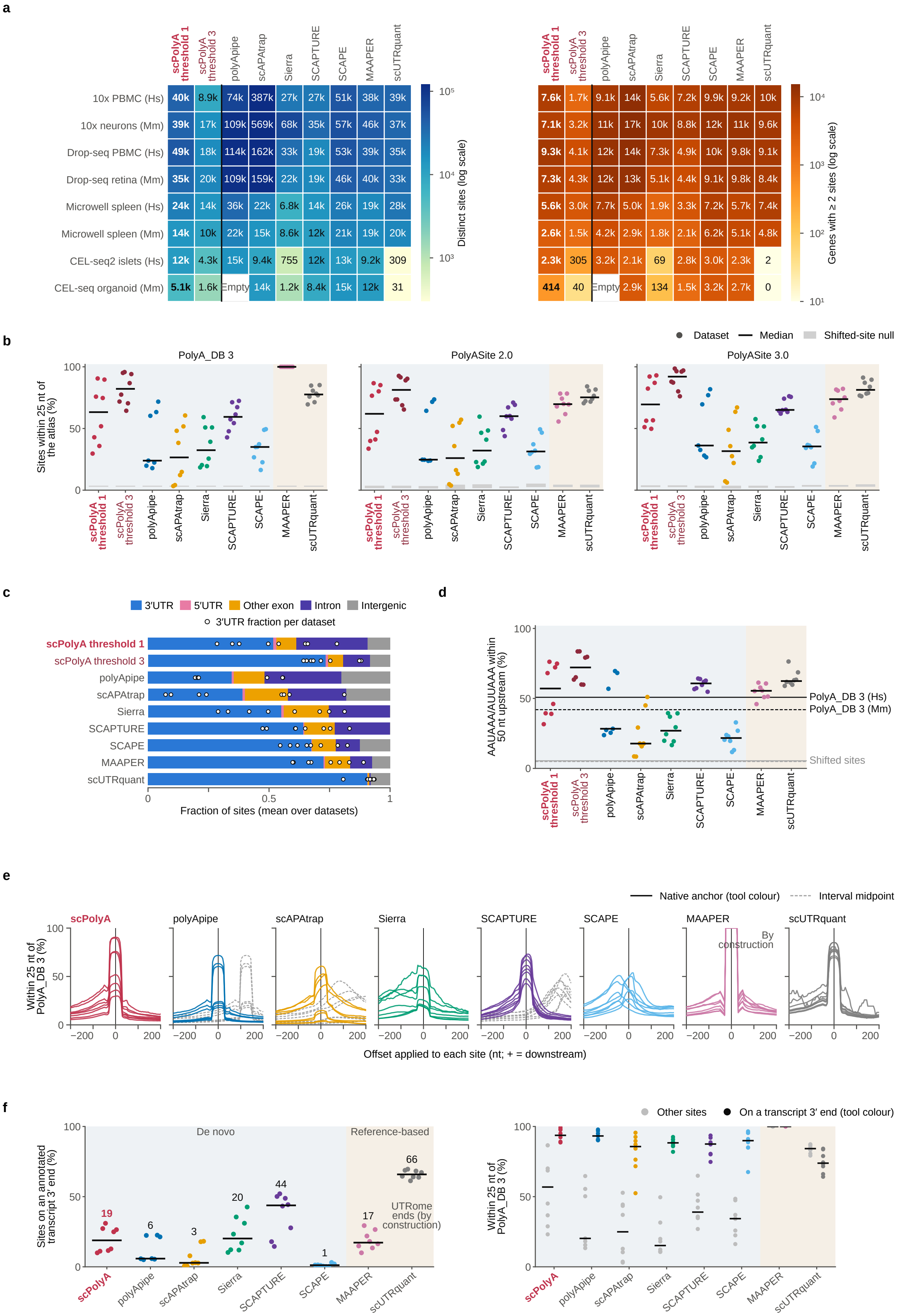
a
scPolyA threshold 1
scPolyA threshold 3
polyApipe
scAPAtrap
Sierra
SCAPTURE
SCAPE
MAAPER
scUTRquant
10x PBMC (Hs) 40k 8.9k 74k 387k 27k 27k 51k 38k 39k
10x neurons (Mm) 39k 17k 109k 569k 68k 35k 57k 46k 37k
Drop-seq PBMC (Hs) 49k 18k 114k 162k 33k 19k 53k 39k 35k
Drop-seq retina (Mm) 35k 20k 109k 159k 22k 19k 46k 40k 33k
Microwell spleen (Hs) 24k 14k 36k 22k 6.8k 14k 26k 19k 28k
Microwell spleen (Mm) 14k 10k 22k 15k 8.6k 12k 21k 19k 20k
CEL-seq2 islets (Hs) 12k 4.3k 15k 9.4k 755 12k 13k 9.2k 309
CEL-seq organoid (Mm) 5.1k 1.6k Empty 14k 1.2k 8.4k 15k 12k 31
Distinct sites (log scale)
7.6k 1.7k 9.1k 14k 5.6k 7.2k 9.9k 9.2k 10k
7.1k 3.2k 11k 17k 10k 8.8k 12k 11k 9.6k
9.3k 4.1k 12k 14k 7.3k 4.9k 10k 9.8k 9.1k
7.3k 4.3k 12k 13k 5.1k 4.4k 9.1k 9.8k 8.4k
5.6k 3.0k 7.7k 5.0k 1.9k 3.3k 7.2k 5.7k 7.4k
2.6k 1.5k 4.2k 2.9k 1.8k 2.1k 6.2k 5.1k 4.8k
2.3k 305 3.2k 2.1k 69 2.8k 3.0k 2.3k 2
414 40 Empty 2.9k 134 1.5k 3.2k 2.7k 0
Genes with ≥ 2 sites (log scale)
b
Dataset
Median
Shifted-site null
PolyA_DB 3
PolyASite 2.0
PolyASite 3.0
Sites within 25 nt of the atlas (%)
c
3′UTR
5′UTR
Other exon
Intron
Intergenic
3′UTR fraction per dataset
Fraction of sites (mean over datasets)
d
AAUAAA/AUUAAA within 50 nt upstream (%)
PolyA_DB 3 (Hs)
PolyA_DB 3 (Mm)
Shifted sites
e
Native anchor (tool colour)
Interval midpoint
scPolyA
By construction
Within 25 nt of PolyA_DB 3 (%)
Offset applied to each site (nt; + = downstream)
f
De novo
Reference-based
Sites on an annotated transcript 3′ end (%)
19
6
3
20
44
1
17
66
UTRome ends (by construction)
Other sites
On a transcript 3′ end (tool colour)
Within 25 nt of PolyA_DB 3 (%)

**Supplementary Figure 7 | Cross-protocol comparisons reveal differences in site yield, reference agreement and reported endpoint position.**

Poly(A)-site methods differ in how they define an RNA end, which affects catalogue size and the relationship between reported coordinates, annotated transcript ends and reference atlases. The extended benchmarking analyses compare these properties using harmonised coordinates and matched datasets.

**a,** The number of distinct sites and genes with at least two reported sites varies by dataset and tool (logarithmic scales). scPolyA is shown separately at tail-read thresholds 1 and 3; the empty CEL-seq organoid entry indicates that polyApipe returned no sites.

**b,** Atlas proximity is assessed independently against PolyA_DB 3, PolyASite 2.0 and PolyASite 3.0. Points represent datasets, black bars medians and grey bars shifted-reference controls; blue and brown shading distinguish *de novo* from reference-guided methods. MAAPER uses PolyA_DB coordinates.

**c,** Genomic-context summaries show how each method distributes its sites among $3'$ UTRs, other transcript regions and intergenic positions. Bars are dataset averages and points the $3'$ UTR fraction for each dataset.

**d,** The frequency of upstream AAUAAA or AUUAAA is compared across methods, alongside the frequencies at reference and shifted-reference sites for human and mouse.

**e,** Systematically shifting reported endpoints across neighbouring positions shows how atlas proximity depends on coordinate placement. Each tool's native site coordinate is also compared, where available, with the midpoint of its reported interval; native definitions include peak boundaries, fitted positions and reference-transcript ends. The offset axis follows transcript orientation.

**f,** Exact agreement with annotated GENCODE transcript ends (left) and atlas proximity for annotated versus other reported sites (right) show how annotation-based positions contribute to reference support. Points are datasets and bars medians; the tools and their settings are detailed in Supplementary Table 3.

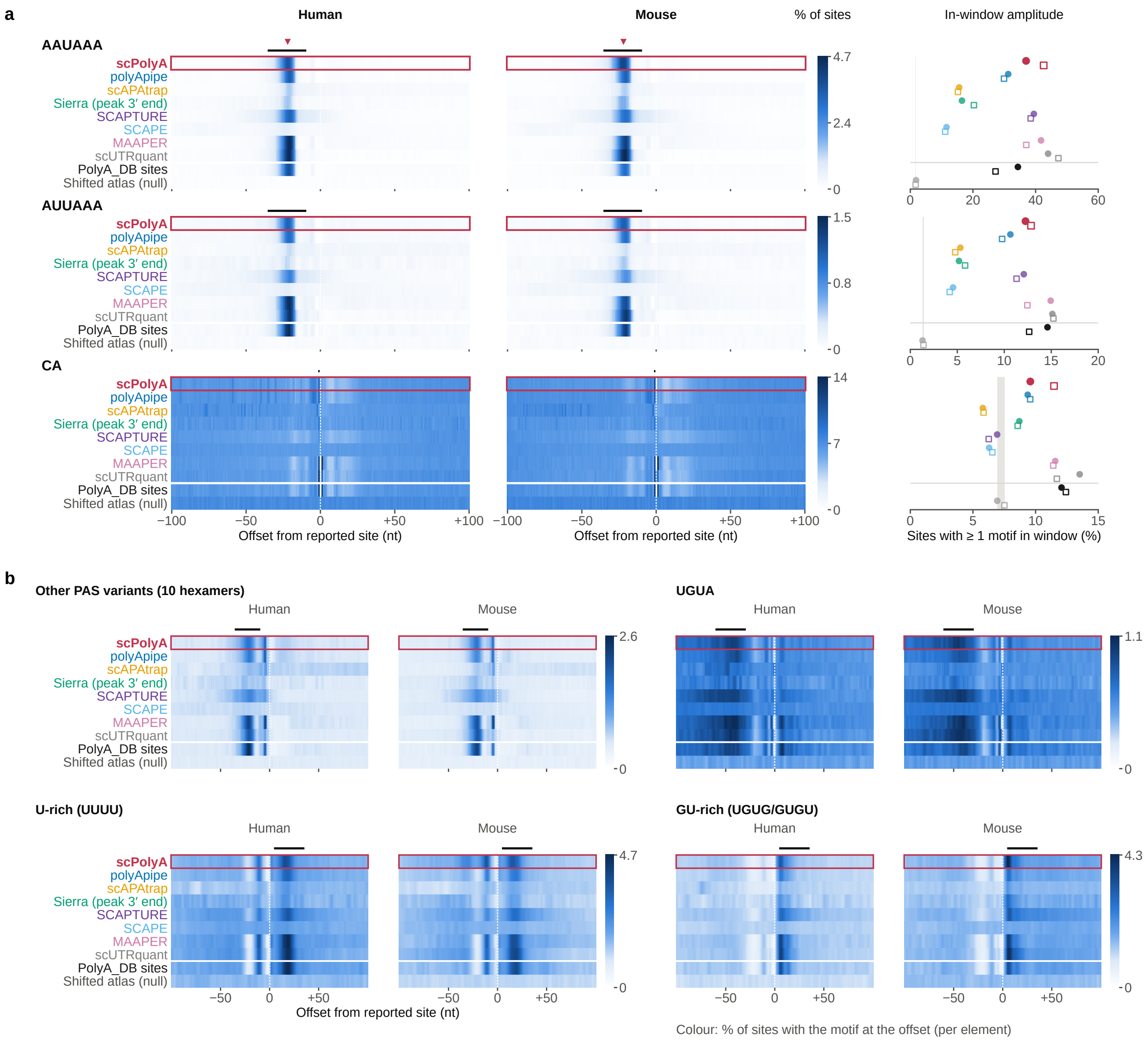

a
Human
Mouse
% of sites
In-window amplitude
AAUAAA
AUUAAA
CA
scPolyA
polyApipe
scAPAtrap
Sierra (peak 3′ end)
SCAPTURE
SCAPE
MAAPER
scUTRquant
PolyA_DB sites
Shifted atlas (null)
4.7
2.4
0
1.5
0.8
14
7
−100
−50
0
+50
+100
Offset from reported site (nt)
0
20
40
60
0
5
10
15
20
Sites with ≥ 1 motif in window (%)
b
Other PAS variants (10 hexamers)
UGUA
U-rich (UUUU)
GU-rich (UGUG/GUGU)
2.6
1.1
4.3
Colour: % of sites with the motif at the offset (per element)

**Supplementary Figure 8 | scPolyA site coordinates recover the sequence architecture of cleavage and polyadenylation.**

The sequences around recognised transcript ends contain upstream polyadenylation signals, cleavage-associated dinucleotides and downstream regulatory elements. Their characteristic positions provide a complementary view of site placement across RNA-end discovery methods.

**a,** AAUAAA and AUUAAA signal positions and the cleavage-associated CA dinucleotide are profiled around sites from each method and from PolyA_DB, with shifted atlas sites as a reference. Colours indicate positional motif frequencies; bars mark the expected windows and the triangle the AAUAAA peak near scPolyA sites. The right-hand summaries show the proportion containing each motif in its canonical window (filled circles, human; open squares, mouse), with a grey shifted-reference range.

**b,** The extended analysis examines ten further PAS hexamers and the upstream UGUA, downstream U-rich and GU-rich motifs. Each motif is summarised over its characteristic window (PAS, −35 to −10 nt; UGUA, −60 to −30 nt; downstream elements, +5 to +35 nt). Profiles use up to 20,000 sites per set, with three-nucleotide smoothing except for CA; scPolyA is shown at threshold 1.

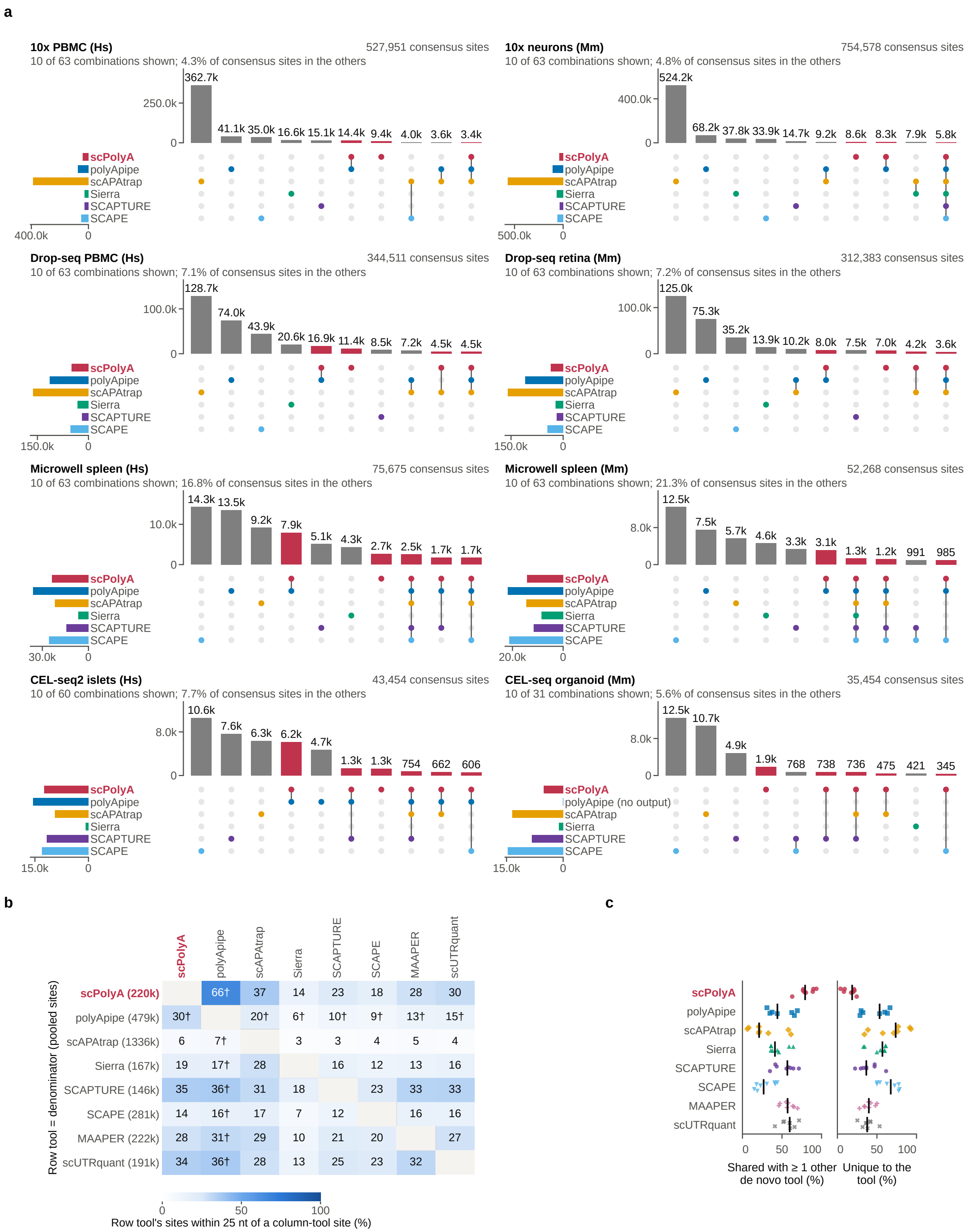

a
10x PBMC (Hs)
527,951 consensus sites
10 of 63 combinations shown; 4.3% of consensus sites in the others
10x neurons (Mm)
754,578 consensus sites
10 of 63 combinations shown; 4.8% of consensus sites in the others
Drop-seq PBMC (Hs)
344,511 consensus sites
10 of 63 combinations shown; 7.1% of consensus sites in the others
Drop-seq retina (Mm)
312,383 consensus sites
10 of 63 combinations shown; 7.2% of consensus sites in the others
Microwell spleen (Hs)
75,675 consensus sites
10 of 63 combinations shown; 16.8% of consensus sites in the others
Microwell spleen (Mm)
52,268 consensus sites
10 of 63 combinations shown; 21.3% of consensus sites in the others
CEL-seq2 islets (Hs)
43,454 consensus sites
10 of 60 combinations shown; 7.7% of consensus sites in the others
CEL-seq organoid (Mm)
35,454 consensus sites
10 of 31 combinations shown; 5.6% of consensus sites in the others
scPolyA
polyApipe
polyApipe (no output)
scAPAtrap
Sierra
SCAPTURE
SCAPE
b
Row tool = denominator (pooled sites)
| | scPolyA | polyApipe | scAPAtrap | Sierra | SCAPTURE | SCAPE | MAAPER | scUTRquant |
| scPolyA (220k) | | 66† | 37 | 14 | 23 | 18 | 28 | 30 |
| polyApipe (479k) | 30† | | 20† | 6† | 10† | 9† | 13† | 15† |
| scAPAtrap (1336k) | 6 | 7† | | 3 | 3 | 4 | 5 | 4 |
| Sierra (167k) | 19 | 17† | 28 | | 16 | 12 | 13 | 16 |
| SCAPTURE (146k) | 35 | 36† | 31 | 18 | | 23 | 33 | 33 |
| SCAPE (281k) | 14 | 16† | 17 | 7 | 12 | | 16 | 16 |
| MAAPER (222k) | 28 | 31† | 29 | 10 | 21 | 20 | | 27 |
| scUTRquant (191k) | 34 | 36† | 28 | 13 | 25 | 23 | 32 | |
0 50 100
Row tool's sites within 25 nt of a column-tool site (%)
c
scPolyA
polyApipe
scAPAtrap
Sierra
SCAPTURE
SCAPE
MAAPER
scUTRquant
Shared with ≥ 1 other de novo tool (%)
Unique to the tool (%)

**Supplementary Figure 9 | RNA-end catalogues share substantial site evidence but also contain method-specific candidates.**

Distinct discovery strategies recover overlapping, yet nonidentical, groups of transcript ends from the same libraries. Comparing the site sets clarifies how much of each catalogue is shared and how differences in catalogue size affect apparent overlap.

**a,** Site coordinates from six *de novo* methods are grouped within 25 nt on the same strand. UpSet diagrams show the ten most frequent tool combinations for each dataset, highlighting those containing scPolyA; header summaries give the number of site groups and combinations, and side bars the groups containing each method.

**b,** The proportion of sites from one method lying within 25 nt of another method's sites is shown as directional containment. Because the row method defines the denominator, overlap can differ between the two directions. Results are pooled over datasets completed by both methods.

**c,** Most scPolyA sites are also represented by at least one other *de novo* method. Across datasets, the proportions shared with another method and unique to each method are shown as individual points and median bars. scPolyA uses threshold 1; polyApipe yielded no sites for the CEL-seq organoid.

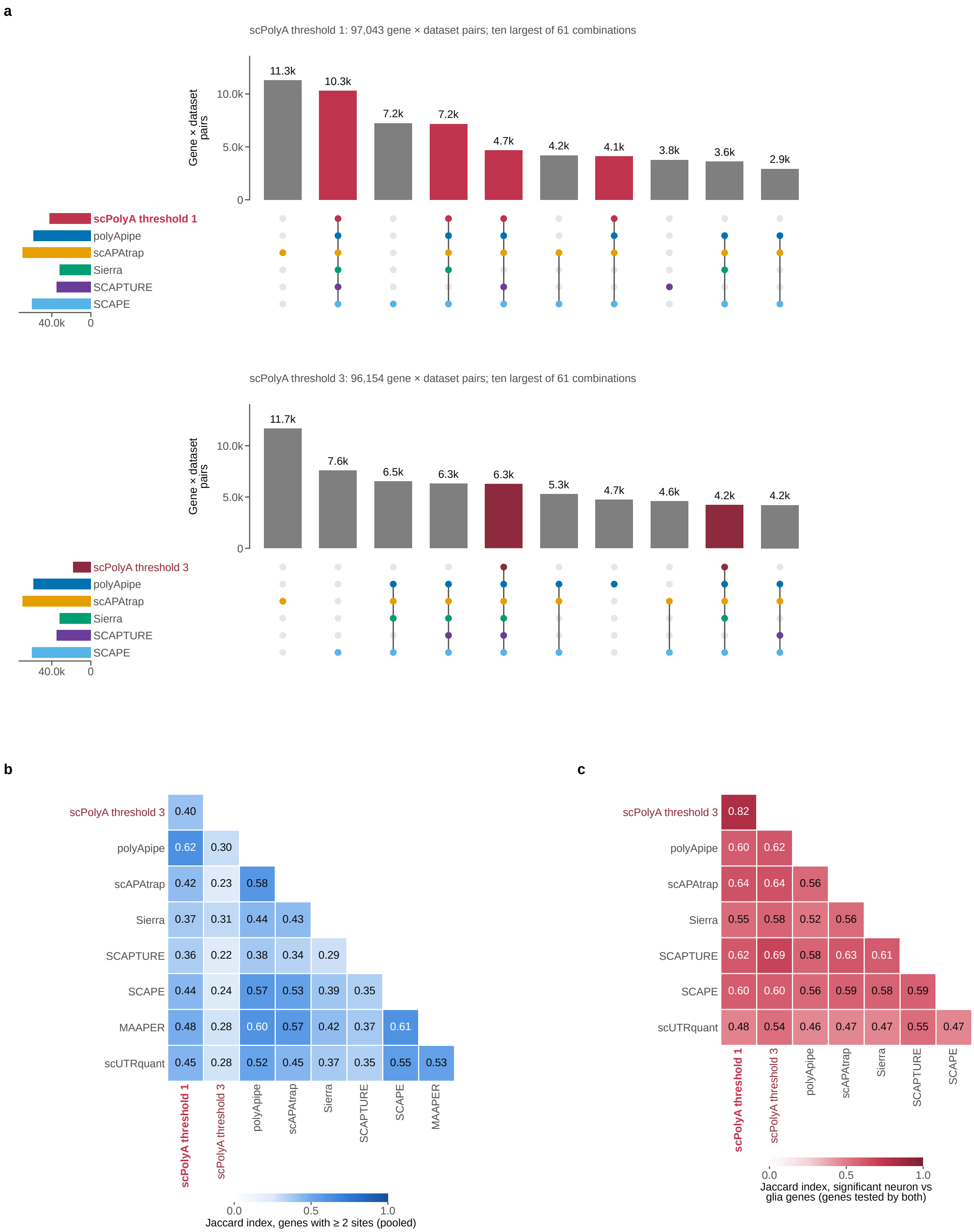
a
scPolyA threshold 1: 97,043 gene × dataset pairs; ten largest of 61 combinations
Gene × dataset pairs
11.3k
10.3k
7.2k
7.2k
4.7k
4.2k
4.1k
3.8k
3.6k
2.9k
scPolyA threshold 1
polyApipe
scAPAtrap
Sierra
SCAPTURE
SCAPE
40.0k
0
scPolyA threshold 3: 96,154 gene × dataset pairs; ten largest of 61 combinations
Gene × dataset pairs
11.7k
7.6k
6.5k
6.3k
6.3k
5.3k
4.7k
4.6k
4.2k
4.2k
scPolyA threshold 3
b
Jaccard index, genes with ≥ 2 sites (pooled)
c
Jaccard index, significant neuron vs glia genes (genes tested by both)

**Supplementary Figure 10 | The definition of a multi-site gene shapes which APA differences each method can detect.**

A poly(A)-site catalogue determines which genes have more than one candidate end, and these gene sets vary among methods. The analysis follows that variation from multi-site gene identification to the genes reported as differentially used between neurons and glia or progenitors.

**a,** UpSet diagrams compare gene-by-dataset pairs with at least two sites across six *de novo* methods and eight single-cell datasets. scPolyA is shown at thresholds 1 (top) and 3 (bottom); the ten largest method combinations are displayed, highlighting those containing scPolyA.

**b,** Pairwise Jaccard similarity quantifies overlap of these multi-site gene sets using datasets in which both methods reported results.

**c,** In 10x mouse-neuron data, the corresponding similarity is calculated for genes showing significant differences in distal-site usage between neurons and glia/progenitors (Benjamini–Hochberg FDR $< 0.05$). The comparison is restricted to genes tested by both methods and includes the scPolyA threshold-3 results from Fig. 4g.

a

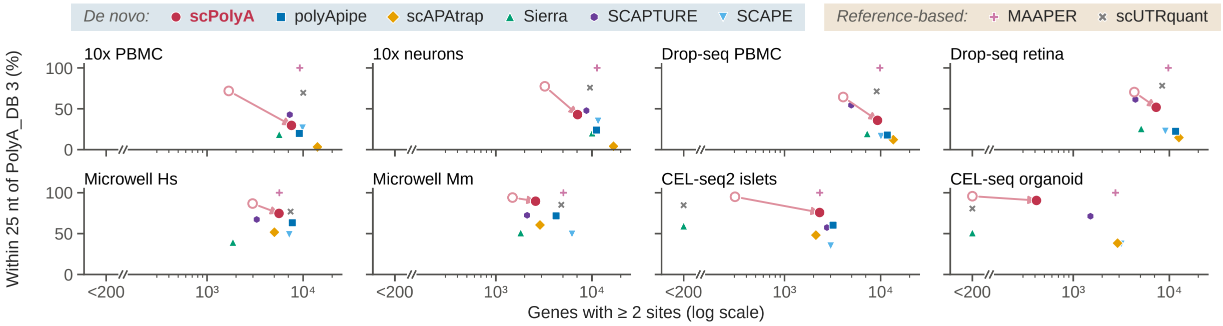

b

| | Human | | | | Mouse | | | |
|---|---|---|---|---|---|---|---|---|
| | 10x PBMC | Drop-seq PBMC | Microwell spleen | CEL-seq2 islets | 10x neurons | Drop-seq retina | Microwell spleen | CEL-seq organoid |
| **scPolyA (threshold 1)** | **28.3** | **2.2** | **0.2** | **1.7** | **22.1** | **1.4** | **0.3** | **0.7** |
| polyApipe | 23.8 | 2.3 | 0.3 | 2.2 | 18.7 | 2.0 | 0.6 | No output |
| scAPAtrap | 4.5 | 9.6 | 5.2 | 6.0 | 5.7 | 9.4 | 4.3 | 5.7 |
| Sierra (peak 3′ end) | 5.9 | 2.6 | 1.4 | 4.5 | 14.7 | 2.0 | 1.8 | 3.5 |
| SCAPTURE | 7.1 | 3.3 | 2.5 | 3.4 | 11.9 | 4.7 | 4.7 | 4.7 |
| SCAPE | 5.2 | 2.4 | 3.3 | 3.1 | 6.6 | 2.6 | 3.9 | 2.7 |
| MAAPER | 2.1 | 2.3 | 1.8 | 3.0 | 4.0 | 3.5 | 3.3 | 4.1 |
| scUTRquant | 3.5 | 3.7 | 3.7 | 3.6† | 5.1 | 5.2 | 4.2 | 9.7† |
| PolyA_DB sites | | 1.0 | | | | 1.8 | | |
| Shifted atlas (null) | | 2.0 | | | | 1.5 | | |

Colour scale: Sites with ≥ 12 A in +1 to +20 nt (%) (0–25)

† Fewer than 500 sites

c d

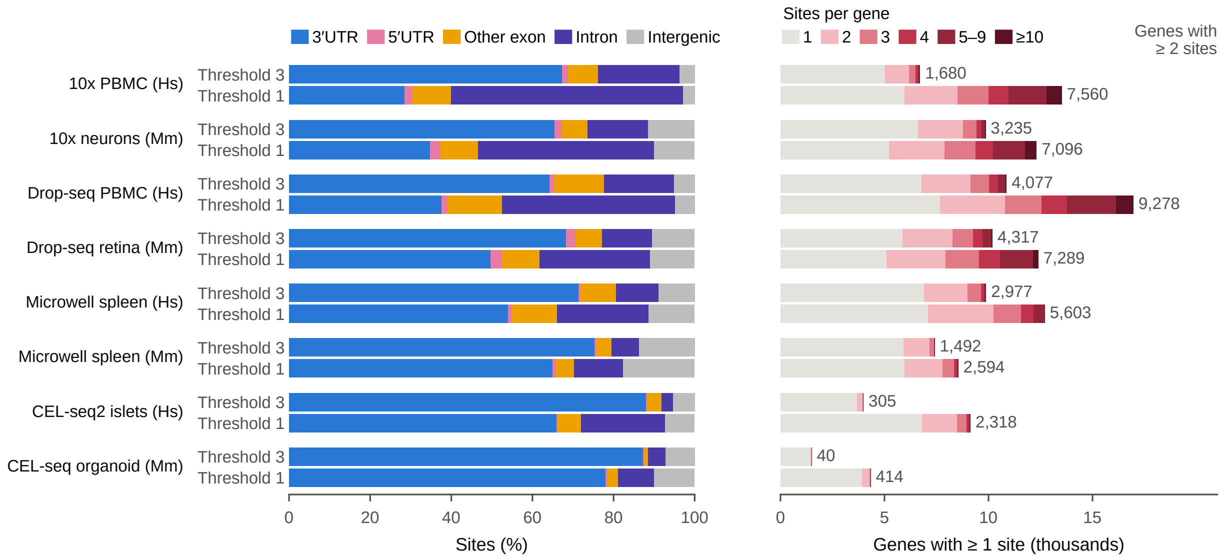

**Supplementary Figure 11 | Relaxing tail-read evidence expands candidate discovery with protocol-dependent changes in site support.**

Reducing the minimum pooled tail-read depth makes additional RNA ends accessible, but the support and transcript context of those candidates differ between library types. The analyses compare catalogue expansion with reference proximity, downstream A-richness and the number of genes eligible for APA studies.

**a,** For each dataset, the number of genes with at least two sites is plotted against the proportion of sites close to PolyA_DB 3. Arrows trace the change in scPolyA from threshold 3 (open) to threshold 1 (filled), allowing the increase in multi-site genes to be considered alongside atlas agreement.

**b,** Downstream A-richness is compared across datasets and methods, using a window of 20 nt after the reported endpoint with at least 12 adenosines. Species-matched atlas and shifted-reference sites provide sequence-context references; a dagger marks methods reporting fewer than 500 sites.

**c,** The genomic context of scPolyA candidates shifts as the tail-read threshold is relaxed, including an increased contribution from intronic sites.

**d,** Gene counts are partitioned by the number of discovered sites per gene at both thresholds, with the multi-site totals annotated. The scPolyA catalogues shown here use read-based discovery without DeepCPS score filtering.

**a**

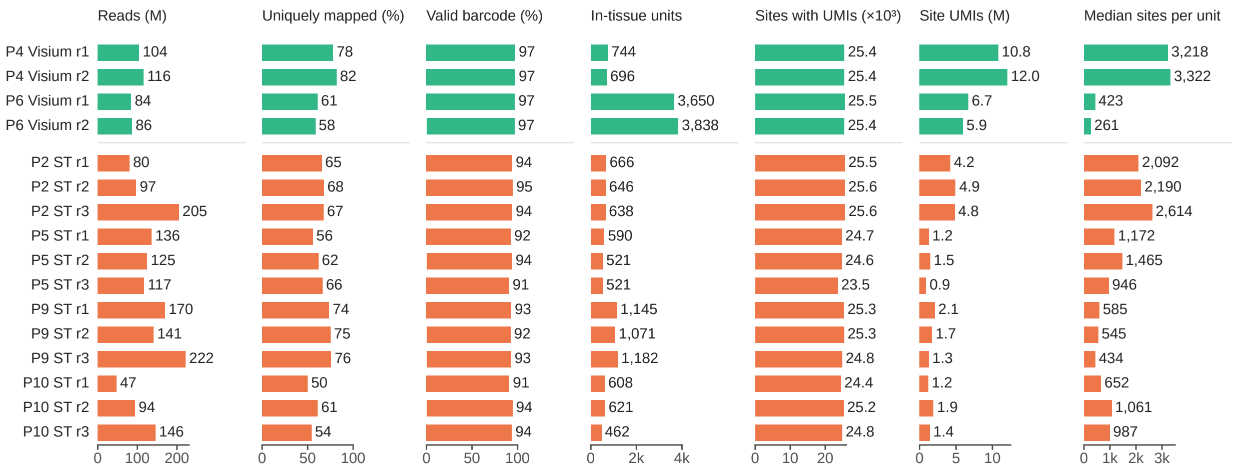

Reads (M)
Uniquely mapped (%)
Valid barcode (%)
In-tissue units
Sites with UMIs (×10³)
Site UMIs (M)
Median sites per unit
P4 Visium r1 104 78 97 744 25.4 10.8 3,218
P4 Visium r2 116 82 97 696 25.4 12.0 3,322
P6 Visium r1 84 61 97 3,650 25.5 6.7 423
P6 Visium r2 86 58 97 3,838 25.4 5.9 261
P2 ST r1 80 65 94 666 25.5 4.2 2,092
P2 ST r2 97 68 95 646 25.6 4.9 2,190
P2 ST r3 205 67 94 638 25.6 4.8 2,614
P5 ST r1 136 56 92 590 24.7 1.2 1,172
P5 ST r2 125 62 94 521 24.6 1.5 1,465
P5 ST r3 117 66 91 521 23.5 0.9 946
P9 ST r1 170 74 93 1,145 25.3 2.1 585
P9 ST r2 141 75 92 1,071 25.3 1.7 545
P9 ST r3 222 76 93 1,182 24.8 1.3 434
P10 ST r1 47 50 91 608 24.4 1.2 652
P10 ST r2 94 61 94 621 25.2 1.9 1,061
P10 ST r3 146 54 94 462 24.8 1.4 987
0 100 200
0 50 100
0 50 100
0 2k 4k
0 10 20
0 5 10
0 1k 2k 3k

**b**

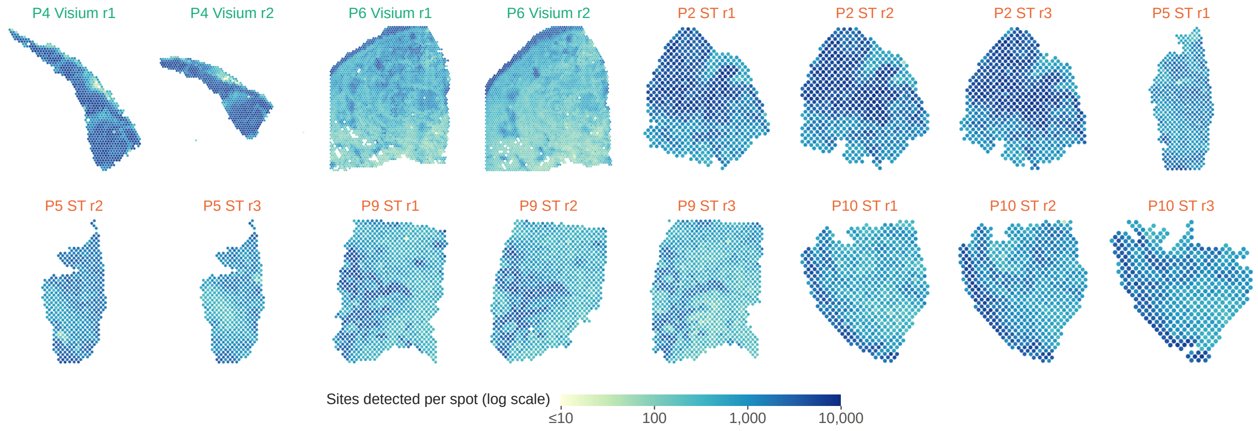

P4 Visium r1
P4 Visium r2
P6 Visium r1
P6 Visium r2
P2 ST r1
P2 ST r2
P2 ST r3
P5 ST r1
P5 ST r2
P5 ST r3
P9 ST r1
P9 ST r2
P9 ST r3
P10 ST r1
P10 ST r2
P10 ST r3
Sites detected per spot (log scale)
≤10
100
1,000
10,000

**Supplementary Figure 12 | Spatial library depth influences the RNA-end measurements available across the cSCC cohort.**

The single-cell and spatial cSCC case study combines sections generated by two technologies with different read lengths and spot-level coverage. These section-level summaries establish the data supporting the tumour-region APA comparisons in Fig. 7.

**a,** Four Visium sections (green; 120-nt cDNA reads) and twelve legacy ST sections (orange; 46-nt cDNA reads) are compared for read depth, mapping and barcode fractions, in-tissue spots, sites with UMIs, total site UMIs and median sites per spot. All sections were analysed with the cohort threshold-3 configuration.

**b,** The number of poly(A) sites detected in each spatial unit is mapped for all 16 sections on a shared logarithmic colour scale. Sections are displayed in separate frames, retaining their within-section distributions while using independent spatial scales.

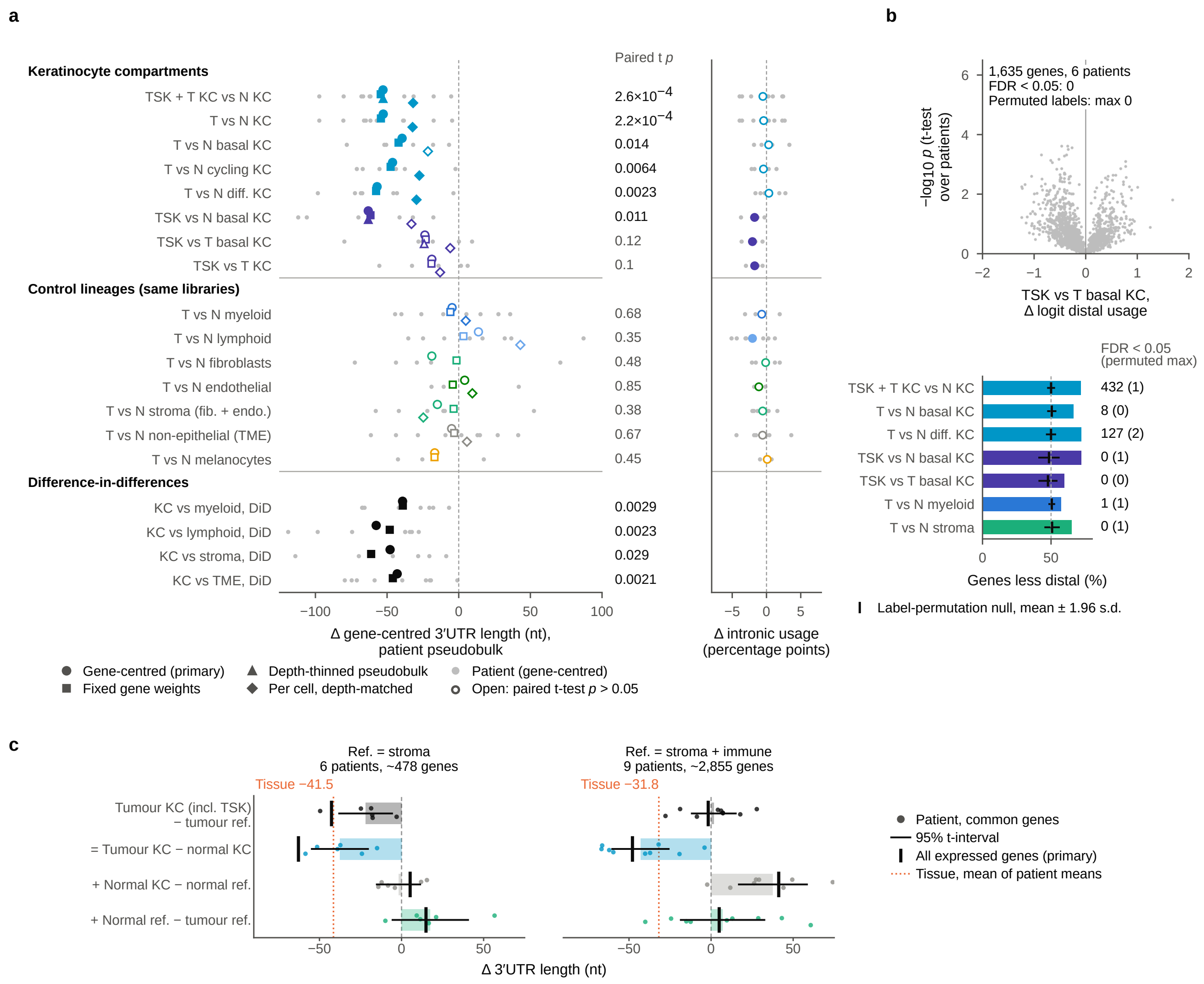
a
b
c
Keratinocyte compartments
TSK + T KC vs N KC
T vs N KC
T vs N basal KC
T vs N cycling KC
T vs N diff. KC
TSK vs N basal KC
TSK vs T basal KC
TSK vs T KC
Control lineages (same libraries)
T vs N myeloid
T vs N lymphoid
T vs N fibroblasts
T vs N endothelial
T vs N stroma (fib. + endo.)
T vs N non-epithelial (TME)
T vs N melanocytes
Difference-in-differences
KC vs myeloid, DiD
KC vs lymphoid, DiD
KC vs stroma, DiD
KC vs TME, DiD
Paired t p
2.6×10−4
2.2×10−4
0.014
0.0064
0.0023
0.011
0.12
0.1
0.68
0.35
0.48
0.85
0.38
0.67
0.45
0.0029
0.0023
0.029
0.0021
Δ gene-centred 3′UTR length (nt), patient pseudobulk
Δ intronic usage (percentage points)
Gene-centred (primary)
Fixed gene weights
Depth-thinned pseudobulk
Per cell, depth-matched
Patient (gene-centred)
Open: paired t-test p > 0.05
1,635 genes, 6 patients
FDR < 0.05: 0
Permuted labels: max 0
−log10 p (t-test over patients)
TSK vs T basal KC, Δ logit distal usage
FDR < 0.05 (permuted max)
TSK + T KC vs N KC 432 (1)
T vs N basal KC 8 (0)
T vs N diff. KC 127 (2)
TSK vs N basal KC 0 (1)
TSK vs T basal KC 0 (0)
T vs N myeloid 1 (1)
T vs N stroma 0 (1)
Genes less distal (%)
Label-permutation null, mean ± 1.96 s.d.
Ref. = stroma
6 patients, ~478 genes
Ref. = stroma + immune
9 patients, ~2,855 genes
Tissue −41.5
Tissue −31.8
Tumour KC (incl. TSK) − tumour ref.
= Tumour KC − normal KC
+ Normal KC − normal ref.
+ Normal ref. − tumour ref.
Δ 3′UTR length (nt)
Patient, common genes
95% t-interval
All expressed genes (primary)
Tissue, mean of patient means

**Supplementary Figure 13 | Tumour-keratinocyte 3′ UTR shortening persists across depth and gene-composition analyses.**

The patient-matched cSCC pattern is examined using alternative estimators and controls for sequencing depth and gene representation. The supplementary analyses also test whether the tissue contrast retains a keratinocyte-state component when the same genes are used across reference populations.

**a,** Tumour-minus-normal 3′ UTR differences across keratinocyte states and control lineages are compared using gene-centred pseudobulks, fixed gene weights, read-depth thinning and depth-matched cells. Symbols distinguish estimators, grey points show patients, and reported $P$ values come from patient-level paired tests. Intronic-usage contrasts appear at right; open symbols mark nonsignificant comparisons.

**b,** Gene-level distal-usage shifts are shown for tumour-specific keratinocytes (TSK) versus tumour basal keratinocytes (top). The lower summary compares the fraction of tested genes favouring proximal ends across contrasts with a within-patient label-permutation distribution; counts indicate genes meeting the FDR criterion.

**c,** The tumour-state, cellular-composition and reference-state components are recalculated on genes with sufficient UMIs in every compared group, using stroma or stroma plus immune cells as the reference. Patient values and 95% intervals are shown alongside the all-gene decomposition from Fig. 7d and the corresponding tissue estimate from Fig. 7g.

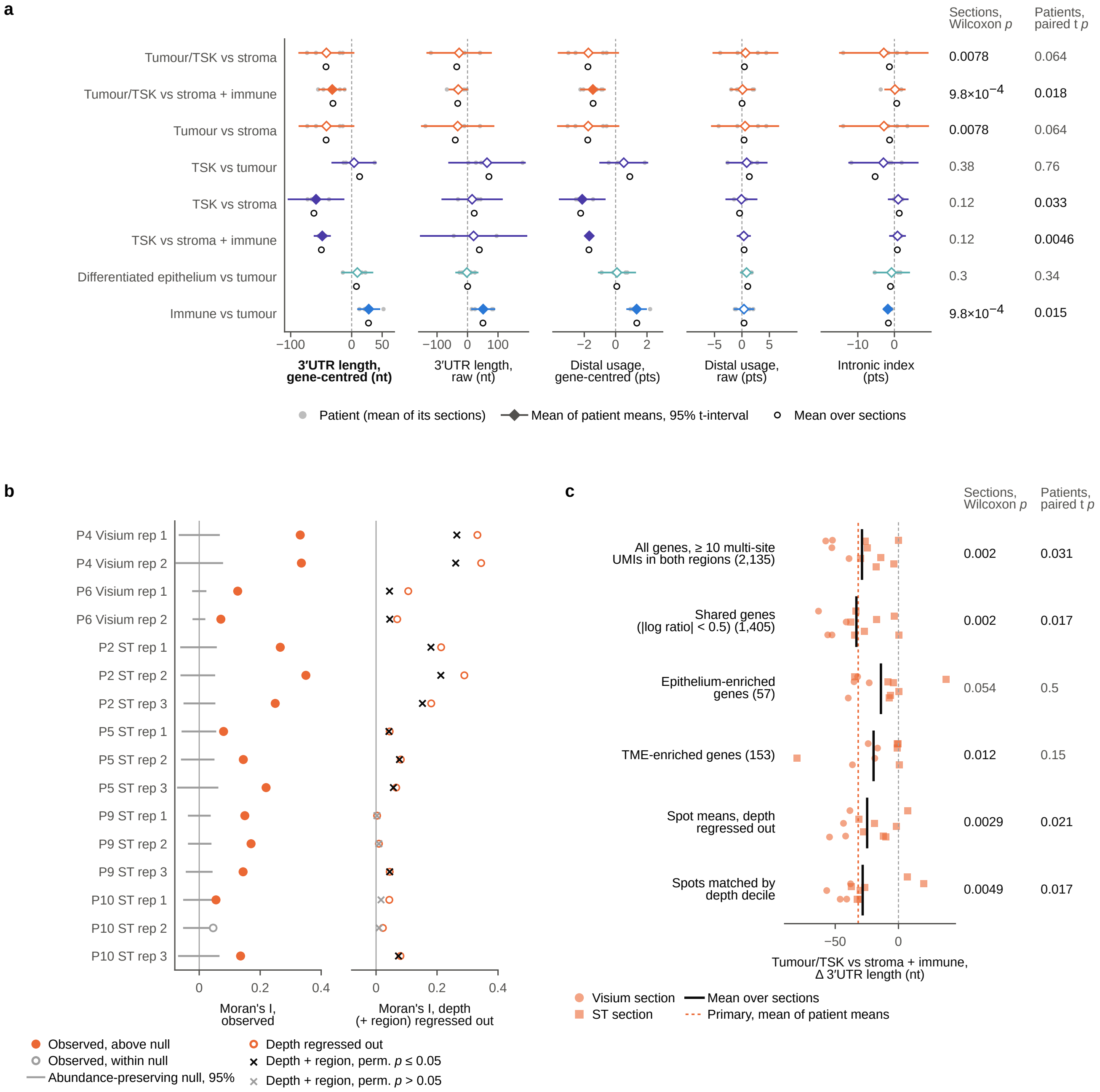

a
Sections, Wilcoxon p
Patients, paired t p
Tumour/TSK vs stroma 0.0078 0.064
Tumour/TSK vs stroma + immune 9.8×10−4 0.018
Tumour vs stroma 0.0078 0.064
TSK vs tumour 0.38 0.76
TSK vs stroma 0.12 0.033
TSK vs stroma + immune 0.12 0.0046
Differentiated epithelium vs tumour 0.3 0.34
Immune vs tumour 9.8×10−4 0.015
3′UTR length, gene-centred (nt)
3′UTR length, raw (nt)
Distal usage, gene-centred (pts)
Distal usage, raw (pts)
Intronic index (pts)
Patient (mean of its sections)
Mean of patient means, 95% t-interval
Mean over sections
b
P4 Visium rep 1
P4 Visium rep 2
P6 Visium rep 1
P6 Visium rep 2
P2 ST rep 1
P2 ST rep 2
P2 ST rep 3
P5 ST rep 1
P5 ST rep 2
P5 ST rep 3
P9 ST rep 1
P9 ST rep 2
P9 ST rep 3
P10 ST rep 1
P10 ST rep 2
P10 ST rep 3
Moran's I, observed
Moran's I, depth (+ region) regressed out
Observed, above null
Observed, within null
Abundance-preserving null, 95%
Depth regressed out
Depth + region, perm. p ≤ 0.05
Depth + region, perm. p > 0.05
c
Sections, Wilcoxon p
Patients, paired t p
All genes, ≥ 10 multi-site UMIs in both regions (2,135) 0.002 0.031
Shared genes (|log ratio| < 0.5) (1,405) 0.002 0.017
Epithelium-enriched genes (57) 0.054 0.5
TME-enriched genes (153) 0.012 0.15
Spot means, depth regressed out 0.0029 0.021
Spots matched by depth decile 0.0049 0.017
Tumour/TSK vs stroma + immune, Δ 3′UTR length (nt)
Visium section
ST section
Mean over sections
Primary, mean of patient means

**Supplementary Figure 14 | Spatial $3'$ UTR shortening remains visible across readouts and follows tissue organisation.**

The tumour-associated APA pattern is examined using alternative outcome definitions, gene sets and sequencing-depth controls. Spatial autocorrelation analyses further assess whether the local measurements have a structured distribution across tissue sections.

**a,** Five APA readouts are compared between tumour/TSK-enriched and reference regions. Grey points indicate patient-level contrasts, diamonds their mean and 95% confidence interval, and open circles section averages; open diamonds denote patient-level $P > 0.05$, and section-level and patient-level test results appear at right.

**b,** Moran's $I$ summarises spatial structure in spot-level $3'$ UTR length. Observed values are compared with an abundance-preserving null, then evaluated after depth adjustment and after adjustment for depth and region identity. Twelve of sixteen sections retain significant structure after both adjustments; black and grey crosses identify significant and nonsignificant spatial-permutation results, respectively.

**c,** The tumour/TSK-versus-stroma-plus-immune contrast is repeated with shared-gene restrictions and sequencing-depth controls. Points represent sections (circles, Visium; squares, legacy ST), bars section-level averages and the dashed line the primary patient-averaged estimate. Patient-level tests account for multiple sections from the same individual; all sites derive from threshold-3 analyses.

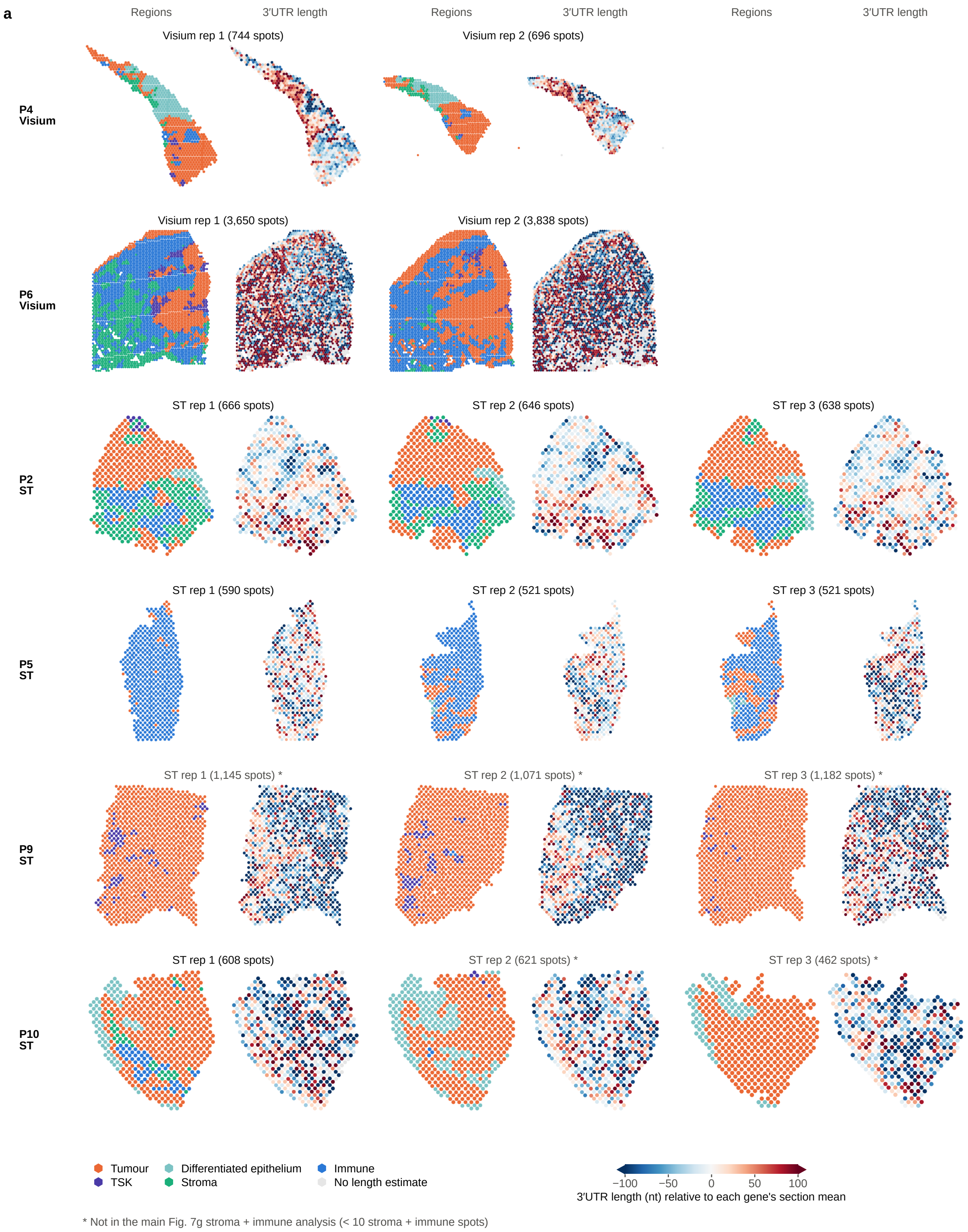
a
Regions
3′UTR length
Regions
3′UTR length
Regions
3′UTR length
P4 Visium
Visium rep 1 (744 spots)
Visium rep 2 (696 spots)
P6 Visium
Visium rep 1 (3,650 spots)
Visium rep 2 (3,838 spots)
P2 ST
ST rep 1 (666 spots)
ST rep 2 (646 spots)
ST rep 3 (638 spots)
P5 ST
ST rep 1 (590 spots)
ST rep 2 (521 spots)
ST rep 3 (521 spots)
P9 ST
ST rep 1 (1,145 spots) *
ST rep 2 (1,071 spots) *
ST rep 3 (1,182 spots) *
P10 ST
ST rep 1 (608 spots)
ST rep 2 (621 spots) *
ST rep 3 (462 spots) *
Tumour
TSK
Differentiated epithelium
Stroma
Immune
No length estimate
−100
−50
0
50
100
3′UTR length (nt) relative to each gene's section mean
* Not in the main Fig. 7g stroma + immune analysis (< 10 stroma + immune spots)

**Supplementary Figure 15 | RNA-end usage and marker-defined tissue regions are mapped across the complete cSCC spatial cohort.**

For each of the 16 sections, the left map shows expression-marker-defined tumour/TSK, differentiated-epithelial, stromal and immune regions, while the matching right map shows spot-level 3′ UTR length centred on each gene's section mean. Sections are grouped by patient to connect local APA patterns with tissue context. Asterisks mark sections with fewer than ten stromal-plus-immune spots, which do not enter the corresponding contrast in Fig. 7g. All maps use the threshold-3 scPolyA analysis.

**a**

| Rule (applied in order) | Entering | Tested | Untestable (control missing) | Passed (log scale) | Removed |
|---|---|---|---|---|---|
| 1 Tested in both data types (tissue: tumour/TSK vs stroma + immune; single cell: T vs N KC) | 2,878 tissue 3,852 single cell | 2,388 | – | **2,388** | – |
| 2 Tissue section-level t-test $p < 0.01$ | 2,388 | 2,388 | 0 | **121** | 2,267 |
| 3 Single-cell KC change of the same sign, patient-level t-test $p < 0.05$ | 121 | 121 | 0 | **55** Chance 34.8 ± 4.8 perm. $p = 5.0{\times}10^{-5}$ | 66 |
| 4 No same-sign myeloid T vs N change ($p < 0.05$): candidates | 55 | 49 | 6 | **50** | 5 |
| 5 Single-cell FDR < 0.05 and composition control not significant in the tissue direction: tier 1 | 50 | 50 | 0 | **13** | 37 |

Passed axis: 10, 100, 1,000

Step-3 overlap under 20,000 permutations, mean ± s.d.

**b**

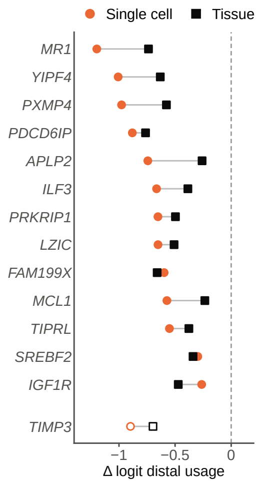

| Gene | Tissue $p < 0.01$ | Single cell, same sign, $p < 0.05$ | Single cell FDR < 0.05 | Myeloid control | Composition control (tumour KC vs TME) |
|---|---|---|---|---|---|
| *MR1* | ● 0.0075 | ● 0.0016 | ● 0.026 | ● 0.18 | ● 0.27 |
| *YIPF4* | ● 0.009 | ● 0.0022 | ● 0.031 | ● 0.57 | ● 0.083 |
| *PXMP4* | ● 0.0015 | ● $3.4{\times}10^{-4}$ | ● 0.013 | ○ Not testable | ● 0.0012, opp. |
| *PDCD6IP* | ● $2.0{\times}10^{-5}$ | ● $8.4{\times}10^{-6}$ | ● 0.0031 | ● 0.16 | ● 0.39 |
| *APLP2* | ● $3.2{\times}10^{-4}$ | ● $4.5{\times}10^{-4}$ | ● 0.014 | ● 0.45 | ● 0.52 |
| *ILF3* | ● 0.0065 | ● $2.7{\times}10^{-6}$ | ● 0.0026 | ● 0.084 | ● 0.21, opp. |
| *PRKRIP1* | ● 0.0061 | ● 0.0014 | ● 0.024 | ● 0.18 | ● 0.88, opp. |
| *LZIC* | ● 0.003 | ● $2.1{\times}10^{-5}$ | ● 0.0038 | ● 0.069 | ● 0.093, opp. |
| *FAM199X* | ● 0.0045 | ● $4.4{\times}10^{-4}$ | ● 0.014 | ● 0.19 | ● 0.29 |
| *MCL1* | ● 0.0067 | ● 0.0017 | ● 0.027 | ● 0.69 | ● 0.33, opp. |
| *TIPRL* | ● 0.0059 | ● 0.0022 | ● 0.031 | ● 0.94 | ● 0.33 |
| *SREBF2* | ● 0.0011 | ● 0.0038 | ● 0.041 | ● 0.39 | ● 0.23, opp. |
| *IGF1R* | ● 0.0066 | ● 0.0033 | ● 0.038 | ○ Not testable | ● 0.46 |
| *TIMP3* | ● 0.0019 | ● 0.0013 | ● 0.023 | ○ Not testable | × 0.043 |

● Passed ○ Not testable × Failed Value: $p$ (FDR); opp., opposite to the tissue direction

**c**

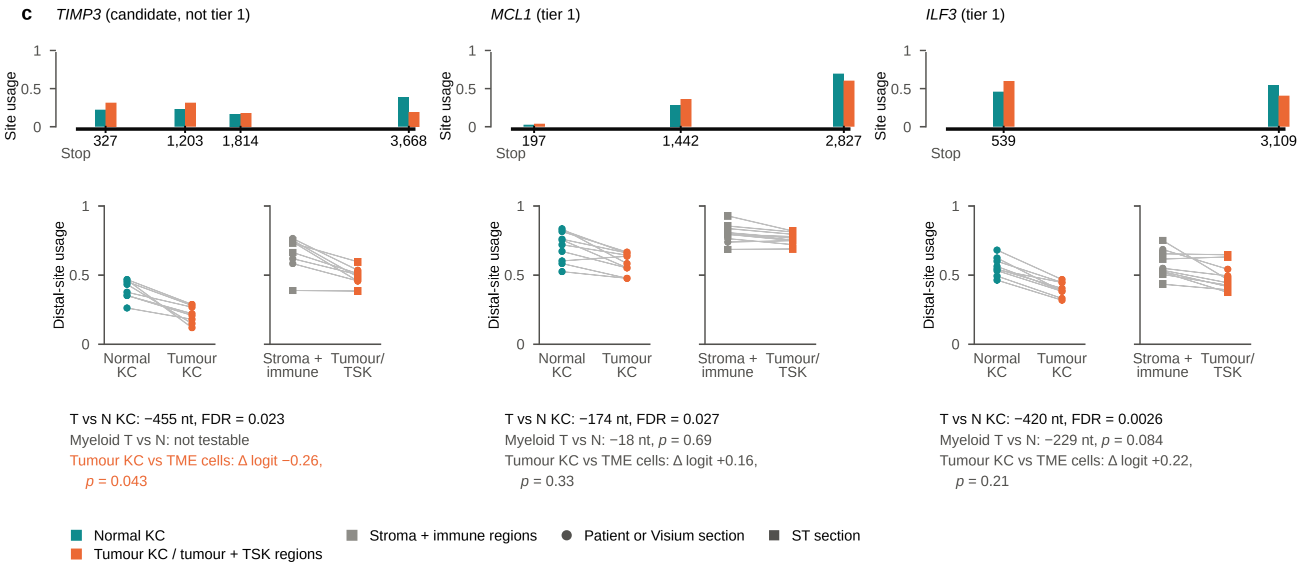

**Supplementary Figure 16 | Convergent single-cell and spatial APA changes prioritise a focused set of cSCC genes.**

Patient-matched cell-type measurements identify changes in tumour keratinocytes, while spatial data indicate which of those changes also occur in tumour-enriched regions. A staged gene-level analysis brings these evidence sources together and identifies thirteen tier-1 candidates for further investigation.

**a,** The selection flow tracks genes through testing, cross-modal agreement and additional cell-composition and myeloid controls. Bars show the number passing each successive criterion on a logarithmic scale, with untestable and excluded genes recorded separately; the diamond shows the overlap expected from 20,000 gene-value permutations.

**b,** For the thirteen tier-1 genes and *TIMP3*, single-cell and spatial effect sizes are aligned with the individual selection criteria and their $P$ or FDR values. Circles and squares denote single-cell and tissue estimates; "opp." identifies a control effect opposite to the tissue direction, and untestable criteria are distinguished from failed tests.

**c,** *TIMP3*, *MCL1* and *ILF3* illustrate how changes in usage at specific terminal ends appear across cellular groups, patients and spatial sections. The accompanying values show the criteria used in gene prioritisation. The complete candidate list is provided in Supplementary Table 5.

**a  Case 1 · Human PBMC**

Claude Opus 5.5 · Claude Desktop · medium effort

Shipped preset unchanged (per-cell metric minimum U = 1); chrM excluded. Delivered 9,030 sites × 5,710 cells, per-cell APA tables and cell-type pooled sums; stated that single cells are sparse.

**Limitations found on checking (run shown in Fig. 8)**

- No substantive limitation found.

GPT-6-Astra · GPT desktop app · medium effort

Preset with per-cell metric minimum U = 10; chrM included (16 sites). Delivered a 9,047-site catalogue, site × cell counts and per-cell APA tables; no cell-type pooling.

**Limitations found on checking**

- Per-cell sparsity and the route to cell-type comparison not stated: at minimum 10, 197 of 593 genes are measurable in QC cells and 94.7% of site-usage rows are blank.
- Intronic-usage support not quantified: 38.6% of gene-level values have no terminal UMIs.

Both runs give identical counts at the 8,990 sites with identical coordinates.

**b  Case 2 · Human PBMC**

GPT-6-Astra · GPT desktop app · medium effort

Matched runs at tail-read depth threshold 3 and 1: sites 9,047 → 45,641, genes with ≥ 2 eligible terminal sites 593 → 2,006, agent screen 80 → 98; added sites audited at read level.

**Limitations found on checking (run shown in Fig. 8)**

- The screen is in effect a gene-depth screen; requiring two sites in the same cell gives 188 → 264 genes.
- Per-cell availability overstates measurability (195,269 → 264,613 gene×cell values have two sites).
- No sequence-context reliability evidence.

Claude Opus 5.5 · Claude Desktop · medium effort

Same matched runs (3 vs 1); diagnosed tail coverage of expression peaks directly and scored sequence context (poly(A) signal, annotated ends, A-richness). Proposed a support-filtered catalogue (317 → 823 genes) and “about 500 additional measurable genes”.

**Limitations found on checking**

- Per-cell availability was taken as measurability. With two supported sites seen in the same cell (U ≥ 5, ≥ 50 QC cells), genes go 68 → 71.
- The support-filtered count is a post hoc catalogue proxy. 2,791 sites called duplicate-only include 527 with non-duplicate coverage; equal site counts are not unchanged sites (356 of 393 differ).

Both runs produced identical native site catalogues, counts and distal-usage tables; they differ in analysis and interpretation. Only the Claude run analysed sequence-context reliability.

**c  Case 3 · Mouse neurons**

GPT-6-Astra · GPT desktop app · medium effort

Screened sites on tail-read ends, unique alignment, signal, A content, annotation and atlas coverage: 16,738 → 8,525 → 8,435 → 441 (>100 nt from PolyA_DB) → 3 stringent → 1 atlas-covered. Priority *Bcl11a*; lower-support *AA387200* (11 molecules); *Ppp2r3d* and *Hmgcs1* set aside because the mm9-based atlas does not cover their sequence.

**Limitations found on checking (run shown in Fig. 8)**

- The lower-support tier was defined after the first screen; *AA387200* sits at its minimum support (10 molecules in the screen).
- *AA387200*’s nearest PolyA_DB site (11,334 nt) is annotated to *AA387200* itself: the atlas covers this gene but places its end downstream of the GENCODE end that the reads support.

Claude Opus 5.5 · Claude Desktop · medium effort

Called a site novel if no PolyA_DB site or GENCODE transcript end lay within its peak ±200 nt (2,175 of 16,738); fixed evidence filters left 380. Tail-read checks against annotated control sites showed most were internal-priming-like (65% A-rich downstream; 6% with a canonical hexamer), leaving *Ppp2r3d* as the single tier-A site and 6 tier-B sites without a hexamer.

**Limitations found on checking**

- *Ppp2r3d* lies in sequence the mm9-based atlas does not cover, so its absence from the atlas is no evidence of a bulk omission.
- A 1-nt strand offset left the run’s own A-run flag unapplied to two minus-strand tier-B sites (*Igfbpl1*, *Psmb1*); applied consistently, 4 tier-B sites remain.
- Counting GENCODE transcript ends as annotation removed 471 sites without a PolyA_DB match, 107 of which pass every evidence filter, among them *Bcl11a*.

Both runs analysed the same native sites with different definitions of novelty: GPT measured distance to PolyA_DB; Claude also excluded sites near GENCODE transcript ends, which removes *Bcl11a* (6 nt from a GENCODE end, 1,652 nt from PolyA_DB).

Human: one donor, one library; mouse: one library. Cell labels from expression.
All runs used scPolyA 2.1.0.

**Supplementary Figure 17 | Independent agent analyses reveal consistent RNA-end measurements and different evidence-based decisions.**

The same three APA questions were examined independently by Claude Opus 5.5 in Claude Desktop and GPT-6-Astra in the GPT desktop application, each at medium reasoning effort. Comparing the two executions of each question shows where scPolyA outputs agree and where the interpretation depends on the scope of the analysis or the reference used to assess a candidate. Both agents used scPolyA v2.1.0 on the same input data, and numerical results were independently recomputed from the outputs. Thick borders identify the executions presented in Fig. 8. For each question, the displayed execution was selected after independent numerical review as having fewer substantive limitations; both executions are shown here.

**a,** In the human PBMC matrix-generation case, UMI counts agree at all 8,990 sites shared at identical genomic coordinates. The runs differ in chromosome scope, including the treatment of chrM, and in the products delivered for downstream APA analysis.

**b,** In the PBMC threshold-comparison case, both agents produce identical native site catalogues, counts and distal-usage tables. Their subsequent analyses and interpretation of the results differ, despite the common quantitative outputs.

**c,** In the mouse-neuron candidate-review case, both agents examine the same native sites but apply different definitions of an atlas-absent end. A screen based on distance to PolyA_DB retains *Bcl11a*, whereas an additional screen against annotated GENCODE transcript ends excludes it, illustrating how genomic context shapes the candidate set.

The PBMC data comprise one donor and library and the mouse data one library; cell identities were assigned from gene expression.

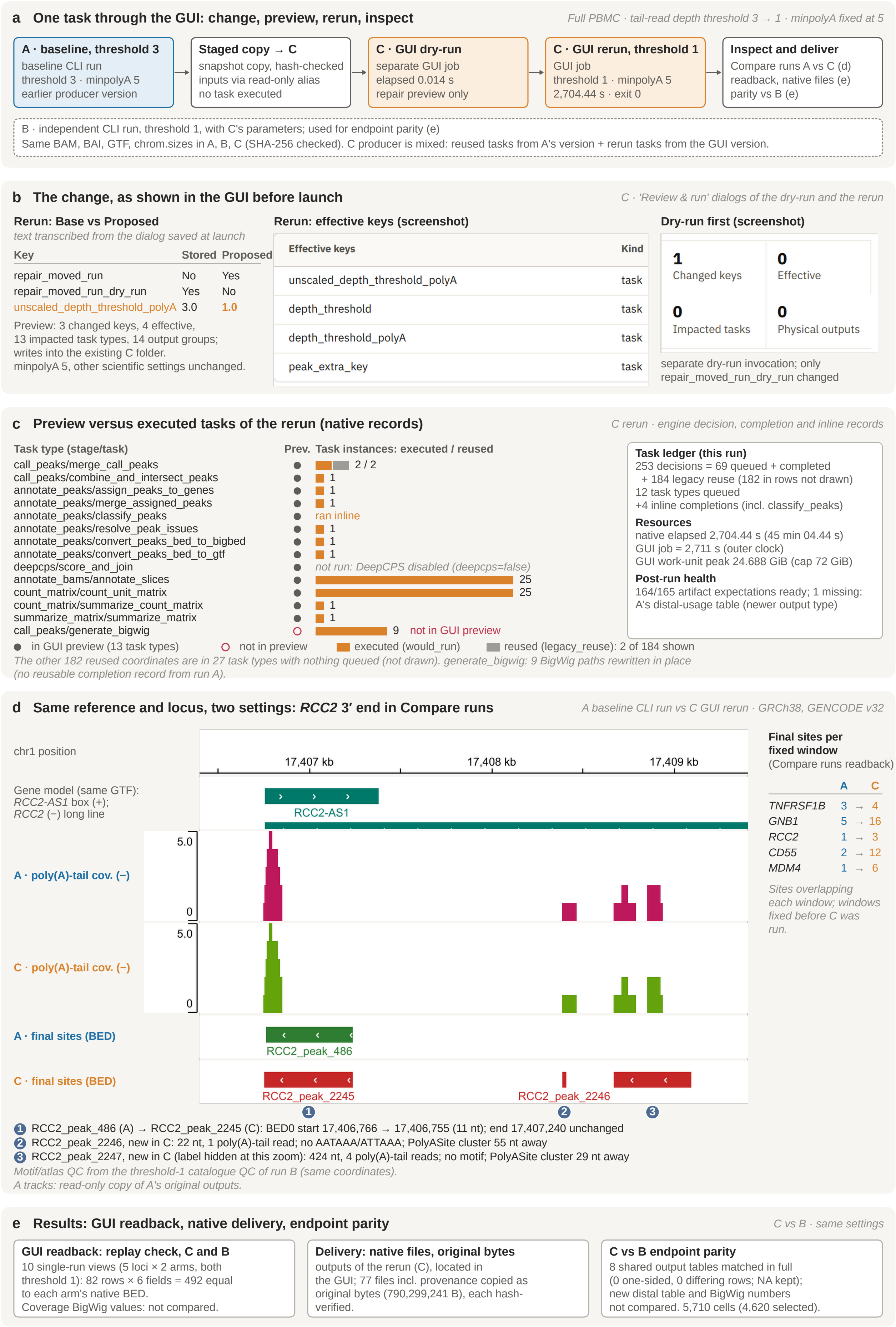
a One task through the GUI: change, preview, rerun, inspect
Full PBMC · tail-read depth threshold 3 → 1 · minpolyA fixed at 5
A · baseline, threshold 3
baseline CLI run
threshold 3 · minpolyA 5
earlier producer version
Staged copy → C
snapshot copy, hash-checked
inputs via read-only alias
no task executed
C · GUI dry-run
separate GUI job
elapsed 0.014 s
repair preview only
C · GUI rerun, threshold 1
GUI job
threshold 1 · minpolyA 5
2,704.44 s · exit 0
Inspect and deliver
Compare runs A vs C (d)
readback, native files (e)
parity vs B (e)
B · independent CLI run, threshold 1, with C's parameters; used for endpoint parity (e)
Same BAM, BAI, GTF, chrom.sizes in A, B, C (SHA-256 checked). C producer is mixed: reused tasks from A's version + rerun tasks from the GUI version.
b The change, as shown in the GUI before launch
C · 'Review & run' dialogs of the dry-run and the rerun
Rerun: Base vs Proposed
text transcribed from the dialog saved at launch
Key
Stored
Proposed
repair_moved_run
No
Yes
repair_moved_run_dry_run
Yes
No
unscaled_depth_threshold_polyA
3.0
1.0
Preview: 3 changed keys, 4 effective,
13 impacted task types, 14 output groups;
writes into the existing C folder.
minpolyA 5, other scientific settings unchanged.
Rerun: effective keys (screenshot)
Effective keys
Kind
unscaled_depth_threshold_polyA
task
depth_threshold
task
depth_threshold_polyA
task
peak_extra_key
task
Dry-run first (screenshot)
1
Changed keys
0
Effective
0
Impacted tasks
0
Physical outputs
separate dry-run invocation; only
repair_moved_run_dry_run changed
c Preview versus executed tasks of the rerun (native records)
C rerun · engine decision, completion and inline records
Task type (stage/task)
Prev.
Task instances: executed / reused
call_peaks/merge_call_peaks
2 / 2
call_peaks/combine_and_intersect_peaks
1
annotate_peaks/assign_peaks_to_genes
1
annotate_peaks/merge_assigned_peaks
1
annotate_peaks/classify_peaks
ran inline
annotate_peaks/resolve_peak_issues
1
annotate_peaks/convert_peaks_bed_to_bigbed
1
annotate_peaks/convert_peaks_bed_to_gtf
1
deepcps/score_and_join
not run: DeepCPS disabled (deepcps=false)
annotate_bams/annotate_slices
25
count_matrix/count_unit_matrix
25
count_matrix/summarize_count_matrix
1
summarize_matrix/summarize_matrix
1
call_peaks/generate_bigwig
9
not in GUI preview
in GUI preview (13 task types)
not in preview
executed (would_run)
reused (legacy_reuse): 2 of 184 shown
The other 182 reused coordinates are in 27 task types with nothing queued (not drawn). generate_bigwig: 9 BigWig paths rewritten in place
(no reusable completion record from run A).
Task ledger (this run)
253 decisions = 69 queued + completed
+ 184 legacy reuse (182 in rows not drawn)
12 task types queued
+4 inline completions (incl. classify_peaks)
Resources
native elapsed 2,704.44 s (45 min 04.44 s)
GUI job ≈ 2,711 s (outer clock)
GUI work-unit peak 24.688 GiB (cap 72 GiB)
Post-run health
164/165 artifact expectations ready; 1 missing:
A's distal-usage table (newer output type)
d Same reference and locus, two settings: RCC2 3′ end in Compare runs
A baseline CLI run vs C GUI rerun · GRCh38, GENCODE v32
chr1 position
17,407 kb
17,408 kb
17,409 kb
Gene model (same GTF):
RCC2-AS1 box (+);
RCC2 (−) long line
RCC2-AS1
A · poly(A)-tail cov. (−)
5.0
0
C · poly(A)-tail cov. (−)
5.0
0
A · final sites (BED)
RCC2_peak_486
C · final sites (BED)
RCC2_peak_2245
RCC2_peak_2246
1
2
3
Final sites per
fixed window
(Compare runs readback)
A C
TNFRSF1B 3 → 4
GNB1 5 → 16
RCC2 1 → 3
CD55 2 → 12
MDM4 1 → 6
Sites overlapping
each window; windows
fixed before C was
run.
1 RCC2_peak_486 (A) → RCC2_peak_2245 (C): BED0 start 17,406,766 → 17,406,755 (11 nt); end 17,407,240 unchanged
2 RCC2_peak_2246, new in C: 22 nt, 1 poly(A)-tail read; no AATAAA/ATTAAA; PolyASite cluster 55 nt away
3 RCC2_peak_2247, new in C (label hidden at this zoom): 424 nt, 4 poly(A)-tail reads; no motif; PolyASite cluster 29 nt away
Motif/atlas QC from the threshold-1 catalogue QC of run B (same coordinates).
A tracks: read-only copy of A's original outputs.
e Results: GUI readback, native delivery, endpoint parity
C vs B · same settings
GUI readback: replay check, C and B
10 single-run views (5 loci × 2 arms, both
threshold 1): 82 rows × 6 fields = 492 equal
to each arm's native BED.
Coverage BigWig values: not compared.
Delivery: native files, original bytes
outputs of the rerun (C), located in
the GUI; 77 files incl. provenance copied as
original bytes (790,299,241 B), each hash-
verified.
C vs B endpoint parity
8 shared output tables matched in full
(0 one-sided, 0 differing rows; NA kept);
new distal table and BigWig numbers
not compared. 5,710 cells (4,620 selected).

**Supplementary Figure 18 | A completed GUI rerun connects a discovery-threshold change to the work that must be repeated.**

An existing scPolyA analysis can be revisited by changing site-selection criteria while reusing unaffected computation. The human 10x Genomics GEM-X 3′ v4 Donor1 PBMC example (GRCh38, GENCODE v32) follows a threshold change through the desktop application's review, execution and genomic inspection steps, and compares the resulting outputs with an independently executed command-line analysis.

**a,** Three related runs establish the comparison: a completed tail-read-depth-3 baseline (A), a GUI-initiated rerun at depth 1 (C) and an independent command-line run at depth 1 (B).

**b,** The configuration and review dialogs record the changed discovery threshold and anticipated task effects before the completed GUI rerun was initiated.

**c,** The predicted task impact is compared with the work actually executed or reused, tracing how a change in the discovery threshold propagates through the analysis.

**d,** The comparative genome browser brings the *RCC2* 3′ end into view across runs, connecting the changed analysis settings to site-level genomic evidence.

**e,** Saved outputs are read back and checked against the independent threshold-1 command-line run. The agreement across eight shared output tables is detailed in Supplementary Fig. 19. The proposed rerun shown in Fig. 3b is a separate mouse-neuron preview; run and invocation identifiers and hashes for this completed PBMC analysis are recorded in the Source Data.

#### Eight shared native output tables agree with an independent run

GUI threshold-1 rerun (C) vs independent threshold-1 command-line run (B); rows matched on primary keys, NA kept, absolute and relative tolerance $10^{-12}$.

| Table | Rows, GUI rerun | Rows, CLI run | Matched | Only in one | Differing |
|---|---:|---:|---:|---:|---:|
| final sites | 45,641 | 45,641 | 45,641 | 0 | 0 |
| site–gene annotation | 45,641 | 45,641 | 45,641 | 0 | 0 |
| site counts | 11,076,021 | 11,076,021 | 11,076,021 | 0 | 0 |
| gene × cell counts | 1,610,746 | 1,610,746 | 1,610,746 | 0 | 0 |
| site usage | 1,920,011 | 1,920,011 | 1,920,011 | 0 | 0 |
| weighted 3′UTR length | 1,610,746 | 1,610,746 | 1,610,746 | 0 | 0 |
| site-level iPUI | 32,909,618 | 32,909,618 | 32,909,618 | 0 | 0 |
| gene-level iPUI | 3,850,542 | 3,850,542 | 3,850,542 | 0 | 0 |

Same settings on both sides: a reproducibility test.
The newer distal-usage table and BigWig values were not compared. Run records: Supplementary Fig. 18; identifiers in the Source Data.

**Supplementary Figure 19 | Matched GUI and command-line analyses reproduce the same native APA measurements.**

The human 10x Genomics GEM-X 3′ v4 Donor1 PBMC library (GRCh38, GENCODE v32) was analysed at tail-read depth 1 through a completed GUI rerun and an independent command-line run. All rows matched across the eight shared native tables, covering site calls, site–gene annotations, site and gene–cell UMI counts, site usage, usage-weighted 3′ UTR length and site- and gene-level intronic polyadenylation indices (iPUI). Rows were matched by primary key, missing values agreed, and floating-point comparisons used an absolute plus relative tolerance of $10^{-12}$; the figure gives the number of rows checked in each table. The independent runs and execution records are described in Supplementary Fig. 18, with identifiers provided in the Source Data. Distal-usage summaries and BigWig tracks were outside this eight-table comparison.

### Supplementary Table legends

**Supplementary Table 1 | Tail-read support predicts reference proximity across single-cell datasets.**

For each of the eight single-cell datasets, the table relates the number of poly(A)-tail reads supporting a candidate to its proximity to established poly(A) sites in PolyA_DB 3 and PolyASite 3.0. Counts and support fractions are presented by tail-read stratum and for the full catalogue, showing how the evidence associated with a fixed read threshold varies between protocols.

**Supplementary Table 2 | Public datasets and spatial sections used throughout the study.**

The table identifies the single-cell benchmarks, spatial platform evaluations and patient-matched cSCC samples, including accessions, species, tissue, protocol, read structure and preprocessing. Cell, spot or bin counts and input read numbers define the analysed populations; cSCC patient totals aggregate the indicated libraries or sections.

**Supplementary Table 3 | Software implementations and execution settings used for the comparative analyses.**

For each method, the table records its evaluated revision, analysis mode, input preparation, principal parameters and supported datasets. It also defines the timed computation stages and resource allocations underlying Fig. 6, allowing the runtime and memory results to be interpreted alongside the distinct operations each method performs.

**Supplementary Table 4 | Parameter-aware reuse reduces the cost of a revised scPolyA analysis.**

The table compares a fresh 10x PBMC analysis with an unchanged rerun and a rerun that raises the pooled tail-read threshold from 1 to 3. Wall time is calculated from the logged pipeline stages and reported with executed-task counts and percentages of the fresh run, showing where completed work is retained and where site-dependent counting and APA summaries are updated (Fig. 6d).

**Supplementary Table 5 | Genes showing corresponding APA changes in tumour keratinocytes and tissue regions.**

The table presents candidate genes from the cSCC analysis, ordered by evidence tier and then single-cell false-discovery rate. It brings together tumour-minus-normal keratinocyte effects (KC), tumour/TSK-minus-reference-region effects (Tissue), contributing patients and sections, and the myeloid and cell-composition controls. For tier-2 candidates, FDR and composition annotations specify which additional tier-1 criteria were not met; complete gene-level measurements are available in Source Data.

**Supplementary Table 6 | Native scPolyA output files link discovered sites to barcode-resolved APA measurements.**

For each output of the evaluated scPolyA pipeline, the table provides the file pattern, observational level, columns and conditions under which it is written. The inventory distinguishes site calls, gene assignments, site-by-cell UMI matrices and APA summaries from optional expression outputs and run metadata, supporting interpretation and reuse of the reported analyses.

### Supplementary Table 1 | Atlas support by number of poly(A)-tail reads

| Dataset | Measure | 1 | 2 | 3 | 4 | 5 | 6 | 7 | 8 | 9 | 10 | >10 | All |
|---|---|---|---|---|---|---|---|---|---|---|---|---|---|
| 10x PBMC | Sites (n) | 23,481 | 8,766 | 3,519 | 2,095 | 1,244 | 970 | 696 | 553 | 441 | 321 | 3,495 | 45,581 |
| 10x PBMC | PolyA_DB 3 (%) | 13.9 | 21.7 | 34.6 | 38.9 | 51.5 | 48.2 | 58.9 | 59.5 | 67.8 | 68.8 | 78.4 | 27.0 |
| 10x PBMC | PolyASite 3.0 s47 (%) | 36.3 | 48.1 | 60.0 | 62.5 | 72.7 | 68.5 | 76.7 | 76.3 | 80.7 | 86.0 | 88.0 | 49.1 |
| 10x neurons | Sites (n) | 14,618 | 6,867 | 3,587 | 2,317 | 1,609 | 1,306 | 1,027 | 867 | 727 | 606 | 9,828 | 43,359 |
| 10x neurons | PolyA_DB 3 (%) | 14.9 | 22.2 | 31.0 | 36.8 | 46.7 | 52.9 | 54.4 | 54.4 | 64.1 | 66.5 | 84.1 | 39.8 |
| 10x neurons | PolyASite 3.0 s47 (%) | 29.2 | 42.4 | 53.8 | 56.2 | 64.2 | 67.7 | 69.6 | 70.6 | 78.1 | 78.2 | 88.5 | 54.0 |
| Drop-seq PBMC | Sites (n) | 32,592 | 14,226 | 6,566 | 4,130 | 2,667 | 1,924 | 1,460 | 1,137 | 957 | 862 | 9,992 | 76,513 |
| Drop-seq PBMC | PolyA_DB 3 (%) | 11.6 | 17.4 | 25.1 | 29.6 | 37.7 | 40.9 | 45.1 | 47.8 | 50.2 | 55.0 | 68.0 | 26.0 |
| Drop-seq PBMC | PolyASite 3.0 s47 (%) | 28.9 | 37.6 | 47.8 | 53.4 | 59.4 | 62.5 | 68.9 | 67.8 | 71.8 | 74.0 | 81.5 | 44.6 |
| Drop-seq retina | Sites (n) | 13,984 | 7,772 | 4,518 | 3,166 | 2,140 | 1,750 | 1,392 | 1,186 | 1,023 | 881 | 13,837 | 51,649 |
| Drop-seq retina | PolyA_DB 3 (%) | 19.5 | 25.1 | 31.2 | 37.5 | 42.0 | 41.5 | 47.1 | 48.4 | 52.4 | 51.9 | 69.5 | 40.2 |
| Drop-seq retina | PolyASite 3.0 s47 (%) | 29.9 | 36.3 | 44.3 | 50.5 | 53.4 | 53.1 | 55.0 | 60.9 | 62.9 | 60.6 | 76.2 | 50.1 |
| Microwell-seq spleen (Hs) | Sites (n) | 8,155 | 3,809 | 2,165 | 1,531 | 1,044 | 934 | 702 | 623 | 533 | 451 | 8,089 | 28,036 |
| Microwell-seq spleen (Hs) | PolyA_DB 3 (%) | 44.6 | 56.6 | 68.5 | 73.4 | 76.5 | 78.3 | 78.6 | 83.3 | 81.8 | 85.4 | 87.7 | 67.5 |
| Microwell-seq spleen (Hs) | PolyASite 3.0 s47 (%) | 70.3 | 79.4 | 87.8 | 90.7 | 92.4 | 92.6 | 92.9 | 94.2 | 94.7 | 96.5 | 94.8 | 84.6 |
| Microwell-seq spleen (Mm) | Sites (n) | 3,139 | 1,947 | 1,351 | 970 | 762 | 644 | 532 | 422 | 372 | 315 | 5,988 | 16,442 |
| Microwell-seq spleen (Mm) | PolyA_DB 3 (%) | 62.1 | 71.2 | 77.4 | 79.9 | 84.6 | 91.0 | 91.9 | 92.9 | 93.8 | 89.5 | 94.8 | 82.6 |
| Microwell-seq spleen (Mm) | PolyASite 3.0 s47 (%) | 79.1 | 83.2 | 88.0 | 89.6 | 92.1 | 95.5 | 95.3 | 96.2 | 96.8 | 94.9 | 97.9 | 90.7 |
| CEL-seq2 islets | Sites (n) | 10,759 | 2,605 | 1,253 | 704 | 473 | 331 | 288 | 227 | 176 | 135 | 1,365 | 18,316 |
| CEL-seq2 islets | PolyA_DB 3 (%) | 37.7 | 68.4 | 80.2 | 86.2 | 92.0 | 89.1 | 91.0 | 95.6 | 95.5 | 93.3 | 95.2 | 56.0 |
| CEL-seq2 islets | PolyASite 3.0 s47 (%) | 55.2 | 83.3 | 91.5 | 96.6 | 97.5 | 97.9 | 99.0 | 98.2 | 98.9 | 97.0 | 99.0 | 70.3 |
| CEL-seq organoid | Sites (n) | 3,148 | 1,070 | 517 | 292 | 207 | 127 | 89 | 88 | 60 | 46 | 380 | 6,024 |
| CEL-seq organoid | PolyA_DB 3 (%) | 75.4 | 89.9 | 93.2 | 95.5 | 95.7 | 96.9 | 94.4 | 93.2 | 93.3 | 95.7 | 91.8 | 83.5 |
| CEL-seq organoid | PolyASite 3.0 s47 (%) | 83.1 | 92.6 | 94.4 | 96.2 | 96.1 | 97.6 | 95.5 | 95.5 | 91.7 | 100.0 | 95.3 | 88.5 |

### Supplementary Table 2 | Datasets and sections

| Block | Dataset | Accession | Protocol | Species | Tissue | Units | Barcode read (nt) | cDNA read (nt) | Aligned reads | Input / preprocessing |
|---|---|---|---|---|---|---|---|---|---|---|
| Single cell | 10x PBMC | 10x Genomics public dataset 5k_ Human_Donor1_PBMC_3p_gem-x | 10x Genomics 3’ GEM-X v4 | human | PBMC (healthy donor 1) | 5,710 cells | 28 | 90 | 209,965,277 | Sequencer FASTQ (10x); R1 = barcode read, R2 = cDNA; Cell Ranger. |
| Single cell | 10x neurons | 10x Genomics public dataset 10k_ Mouse_Neurons_3p_gemx | 10x Genomics 3’ GEM-X v4 | mouse | brain neurons | 12,441 cells | 28 | 90 | 506,490,799 | Sequencer FASTQ (10x); R1 = barcode read, R2 = cDNA; Cell Ranger. |
| Single cell | Drop-seq PBMC | GSE132044 (Ding et al. 2020) GSM3837164/GSM3837169; SRR9169210,SRR9169215 | Drop-seq | human | PBMC (PBMC2) | 6,412 cells | 50 | 50 | 313,530,207 | STARsolo; SRA _1 = barcode read, _2 = cDNA (STARsolo readFilesIn _2 _1); two runs (SRR9169210, SRR9169215), first shown. |
| Single cell | Drop-seq retina | GSE63472 GSM1626793 (Macosko et al. 2015); SRR1853178 | Drop-seq | mouse | retina (P14) | 6,600 cells | 20 | 60 | 130,455,523 | STARsolo; SRA _1 = barcode read, _2 = cDNA. |
| Single cell | Microwell-seq spleen (Hs) | GSE134355 GSM4008649; SRR9887674 | Microwell-seq | human | adult spleen | 6,924 cells | 150 | 150 | 62,797,866 | STARsolo; SRA _1 = barcode read (150 nt; barcodes, linkers, UMI, oligo(dT); STARsolo CB_UMI_Complex), _2 = cDNA. |
| Single cell | Microwell-seq spleen (Mm) | GSE153562 GSM4647301 (MCA3); SRR12119321 | Microwell-seq | mouse | adult spleen | 1,970 cells | 150 | 150 | 39,310,470 | STARsolo; SRA _1 = barcode read (150 nt; barcodes, linkers, UMI, oligo(dT); STARsolo CB_UMI_Complex), _2 = cDNA. |
| Single cell | CEL-seq2 islets | GSE85241 (Muraro et al. 2016); SRR4003803 | CEL-seq2 | human | pancreatic islets (donor D30, library 1) | 96 cells | 75–76 | 75–76 | 5,119,940 | STARsolo; SRA _1 = barcode read, _2 = cDNA; lengths vary by 1 nt between records. |
| Single cell | CEL-seq organoid | GSM1524282 (series of GSM1524282-GSM1524287); SRR1610598 | CEL-seq | mouse | intestinal organoid | 96 cells | 51 | 51 | 8,044,049 | STARsolo; SRA _1 = barcode read, _2 = cDNA. |
| Spatial | 10x Visium cSCC P4 (Hs) | GSE144239 (P4 rep1); within GSE144240 | 10x Visium | human | cSCC | 744 spots (55 $\mu$m) | 28 | 120 | 83,482,631 | Submitted BAM $\rightarrow$ bamtofastq $\rightarrow$ STARsolo (GRCh38/v32); submitted in-tissue spots; bamtofastq from Cell Ranger 9.0.0; STARsolo 2.7.11b |
| Spatial | ST cSCC P2 (Hs) | GSE144239 (P2 rep1); within GSE144240 | ST | human | cSCC | 666 spots (110 $\mu$m) | 31 | 46 | 56,089,000 | Submitted BAM $\rightarrow$ bamtofastq $\rightarrow$ STARsolo (GRCh38/v32); submitted in-tissue spots; bamtofastq from Cell Ranger 9.0.0; STARsolo 2.7.11b |
| Spatial | 10x Visium brain anterior (Mm) | 10x V1_Mouse_Brain_Sagittal_Anterior | 10x Visium | mouse | brain | 2,695 spots (55 $\mu$m) | 16 (CB) + 12 (UMI) | 120 | 291,555,054 | Space Ranger BAM filtered to primary CB/UB reads of in-tissue spots |
| Spatial | 10x Visium HD 3′ brain (Mm) | 10x Visium HD 3′ Mouse Brain Fresh Frozen | 10x Visium HD 3′ | mouse | brain | 9,890 bins (50 $\mu$m) | 43 | 75 | 175,010,450 | Space Ranger 4.1.0 against GRCm38/M23; 50-$\mu$m bins |
| Spatial | Slide-seqV2 testis (Mm) | SRR12805494 | Slide-seqV2 | mouse | testis | 2,097 bins (50 $\mu$m) | XC 14 + XM 9 (tags) | 31–56 | 49,788,478 | Deposited aligned BAM $\rightarrow$ FASTQ (soft clips kept) $\rightarrow$ STARsolo; beads binned ~50 $\mu$m (77 pixels); read lengths suggest poly(A) trimming before deposition |
| Spatial | DBiT-seq embryo (Mm) | SRR10517906 | DBiT-seq | mouse | E12 embryo | 1,673 pixels (50 $\mu$m) | 26 | 100 | 23,377,560 | STARsolo; 50-$\mu$m pixels as given |
| Spatial | HDST olfactory bulb (Mm) | SRR10125060 | HDST | mouse | olfactory bulb | 1,387 bins | UMI 6 + x, y (read name) | 176 | 24,923,356 | STARsolo; 2-$\mu$m beads merged into 25 $\times$ 25-well bins |
| Spatial | Stereo-seq brain (Mm) | GSE256319, GSM8093734; SRR28048223 | Stereo-seq | mouse | brain | 19,012 bins (50 $\mu$m) | 35 | 110 | 10,361,484 | First 20 M read pairs; raw R2 110 nt, terminal 10 nt removed with cutadapt -u -10 before alignment (100 nt retained); terminal sequence consistent with an appended DNBSEQ sample-index read; STAR; bin100 (50 $\mu$m) |
| cSCC cohort | cSCC scRNA-seq P1 | GSE144236 | 10x 3′ v2 | human | skin, 4 libraries (2 normal, 2 tumor) | 2,380 cells | n.a. | 98 | 552,015,486 | Submitted BAM $\rightarrow$ bamtofastq $\rightarrow$ STARsolo (GRCh38/v32); published cells as whitelist |
| cSCC cohort | cSCC scRNA-seq P2 | GSE144236 | 10x 3′ v2 | human | skin, 2 libraries (1 normal, 1 tumor) | 6,824 cells | n.a. | 98 | 386,476,803 | Submitted BAM $\rightarrow$ bamtofastq $\rightarrow$ STARsolo (GRCh38/v32); published cells as whitelist |
| cSCC cohort | cSCC scRNA-seq P3 | GSE144236 | 10x 3′ v2 | human | skin, 3 libraries (1 normal, 2 tumor) | 1,742 cells | n.a. | 98 | 89,983,006 | Submitted BAM $\rightarrow$ bamtofastq $\rightarrow$ STARsolo (GRCh38/v32); published cells as whitelist |
| cSCC cohort | cSCC scRNA-seq P4 | GSE144236 | 10x 3′ v2 | human | skin, 2 libraries (1 normal, 1 tumor) | 7,956 cells | n.a. | 98 | 443,026,088 | Submitted BAM $\rightarrow$ bamtofastq $\rightarrow$ STARsolo (GRCh38/v32); published cells as whitelist |
| cSCC cohort | cSCC scRNA-seq P5 | GSE144236 | 10x 3′ v2 | human | skin, 2 libraries (1 normal, 1 tumor) | 2,494 cells | n.a. | 98 | 455,973,339 | Submitted BAM $\rightarrow$ bamtofastq $\rightarrow$ STARsolo (GRCh38/v32); published cells as whitelist |
| cSCC cohort | cSCC scRNA-seq P6 | GSE144236 | 10x 3′ v2 | human | skin, 2 libraries (1 normal, 1 tumor) | 6,660 cells | n.a. | 98 | 506,186,696 | Submitted BAM $\rightarrow$ bamtofastq $\rightarrow$ STARsolo (GRCh38/v32); published cells as whitelist |
| cSCC cohort | cSCC scRNA-seq P7 | GSE144236 | 10x 3′ v2 | human | skin, 2 libraries (1 normal, 1 tumor) | 5,511 cells | n.a. | 98 | 325,317,051 | Submitted BAM $\rightarrow$ bamtofastq $\rightarrow$ STARsolo (GRCh38/v32); published cells as whitelist |
| cSCC cohort | cSCC scRNA-seq P8 | GSE144236 | 10x 3′ v2 | human | skin, 4 libraries (2 normal, 2 tumor) | 3,502 cells | n.a. | 98 | 417,642,571 | Submitted BAM $\rightarrow$ bamtofastq $\rightarrow$ STARsolo (GRCh38/v32); published cells as whitelist |
| cSCC cohort | cSCC scRNA-seq P9 | GSE144236 | 10x 3′ v2 | human | skin, 2 libraries (1 normal, 1 tumor) | 6,519 cells | n.a. | 98 | 517,347,025 | Submitted BAM $\rightarrow$ bamtofastq $\rightarrow$ STARsolo (GRCh38/v32); published cells as whitelist |
| cSCC cohort | cSCC scRNA-seq P10 | GSE144236 | 10x 3′ v2 | human | skin, 2 libraries (1 normal, 1 tumor) | 4,569 cells | n.a. | 98 | 369,979,272 | Submitted BAM $\rightarrow$ bamtofastq $\rightarrow$ STARsolo (GRCh38/v32); published cells as whitelist |
| cSCC cohort | cSCC Visium P4 | GSE144239; within GSE144240 | 10x Visium | human | cSCC | 1,440 spots in 2 sections | n.a. | 120 | 180,536,176 | Submitted BAM $\rightarrow$ bamtofastq $\rightarrow$ STARsolo (GRCh38/v32); in-tissue spots as whitelist; bamtofastq from Cell Ranger 9.0.0; STARsolo 2.7.11b |

| Block | Dataset | Accession | Protocol | Species | Tissue | Units | Barcode read (nt) | cDNA read (nt) | Aligned reads | Input / preprocessing |
|---|---|---|---|---|---|---|---|---|---|---|
| cSCC cohort | cSCC Visium P6 | GSE144239; within GSE144240 | 10x Visium | human | cSCC | 7,488 spots in 2 sections | n.a. | 120 | 117,951,621 | Submitted BAM → bamtofastq → STARsolo (GRCh38/v32); in-tissue spots as whitelist; bamtofastq from Cell Ranger 9.0.0; STARsolo 2.7.11b |
| cSCC cohort | cSCC ST P2 | GSE144239; within GSE144240 | ST | human | cSCC | 1,950 spots in 3 sections | n.a. | 46 | 270,292,745 | Submitted BAM → bamtofastq → STARsolo (GRCh38/v32); in-tissue spots as whitelist; bamtofastq from Cell Ranger 9.0.0; STARsolo 2.7.11b |
| cSCC cohort | cSCC ST P5 | GSE144239; within GSE144240 | ST | human | cSCC | 1,632 spots in 3 sections | n.a. | 46 | 85,047,176 | Submitted BAM → bamtofastq → STARsolo (GRCh38/v32); in-tissue spots as whitelist; bamtofastq from Cell Ranger 9.0.0; STARsolo 2.7.11b |
| cSCC cohort | cSCC ST P9 | GSE144239; within GSE144240 | ST | human | cSCC | 3,398 spots in 3 sections | n.a. | 46 | 310,791,568 | Submitted BAM → bamtofastq → STARsolo (GRCh38/v32); in-tissue spots as whitelist; bamtofastq from Cell Ranger 9.0.0; STARsolo 2.7.11b |
| cSCC cohort | cSCC ST P10 | GSE144239; within GSE144240 | ST | human | cSCC | 1,691 spots in 3 sections | n.a. | 46 | 118,053,716 | Submitted BAM → bamtofastq → STARsolo (GRCh38/v32); in-tissue spots as whitelist; bamtofastq from Cell Ranger 9.0.0; STARsolo 2.7.11b |

1 cSCC P3 normal skin (Supplementary Fig. 2 QC only) GSE144236; GSM4284230; SRR11832843. Human normal skin, 10x 3′ v2; 1,296 cells and 32,578,919 retained alignments; barcode read 26 nt (16-nt cell barcode + 10-nt UMI), cDNA read 98 nt. This library is included in the cSCC P3 cohort aggregate and is shown separately for read-structure QC; it is not a ninth single-cell benchmark dataset and must not be counted again in cohort totals.

2 Stereo-seq brain Raw R2 reads were 110 nt. The final 10 nt were mainly eight fixed sequences (88% of the first 1 M reads), consistent with an appended DNBSEQ sample-index read. Trimming retained 100 nt and reduced alignments with ≥5-nt 3′ soft clips from 95.8% to 5.3%. The fraction passing the poly(A)-tail rule increased from 0.024% to 1.7% (159,317/9,216,636 eligible primary alignments; 10,361,484 total aligned input records).

### Supplementary Table 3 | Software, versions, key parameters and run conditions

| Tool | Version | Form | Key parameters (as run) | Input | SC timing scope | SC CPU limit | SC memory | Spatial evaluation | Timing qualification |
|---|---|---|---|---|---|---|---|---|---|
| scPolyA | scpolya-paper-freeze-20261021; dependencies: samtools 1.19.2; UMI-tools 1.1.6; bedtools 2.31.1; featureCounts (Subread 2.0.1); pysam 0.22.1; pyranges 0.1.4; gffutils 0.13; UCSC bedGraphToBigWig and bedToBigBed | Python command-line workflow (main.py run) | coverage scaling expr; expression threshold dynamic max_norm_fixed 0.05; poly(A)-tail support 1 (benchmark) / 3 (cSCC); minpolyA 5; nonA 1; MAPQ 10; merge distance 10/10; misprime ratio 0.9 (intronic 0.7); terminal-aware; peak source merged; PSI min gene UMIs 1 | Prepared CB/UB BAM (whitelist cells); GENCODE GTF | Whole native invocation; tail-support threshold 1; reference setup, PAS discovery, cell matrix and enabled PSI, weighted-UTR and iPUI summaries | 16-CPU quota | Method-stage cgroup memory peak; 256-GiB allocation | Evaluated | 8 completed single-cell runs; tail-support threshold 1 |
| polyApipe + polyApiper | polyApipe source version a6dd4bb; polyApiper 0.1.0 | Python script + R package (polyApiper) | depth threshold 1; min poly(A) 5; non-A 0; MAPQ 10; single cell: native defaults, no additional mispriming prefilter; spatial: retain misprime=False | Prepared CB/UB BAM; polyApiper organism reference from the same GTF | Peak calling and annotation; summed stage runtimes and maximum stage memory peak | 16-CPU quota | Maximum method-stage cgroup memory peak; 256-GiB allocation | Evaluated | 8 completed runs; one empty output is shown but excluded from summaries |
| scAPAtrap | 0.2.0 | R package (scripted) | tails.search = ”no”; coverage cutoff 10; min cells 10; min count 10; max peak width 1000; d 100 | Prepared CB/UB BAM | Whole native scAPAtrap() call; tails.search = ”no” | 16-CPU quota | Method-stage cgroup memory peak; 256-GiB allocation | Evaluated with tails.search = ”no”; timing not reported; brain Visium did not complete | 8 completed single-cell runs; no out-of-memory failure |
| Sierra | 0.99.27 | R package (scripted) | junction count/proportion 50/ 0.05; peak count/proportion 200/0.05; coverage 500; single cell: native default CountPeaks output; no A/T-stretch filter (pA/pT flags retained as annotations); spatial sites are filtered with the pA/pT filter | Prepared CB/UB BAM; GENCODE GTF; BSgenome package | Junction detection and Sierra; summed stage runtimes and maximum stage memory peak; median 29.16 s/M aligned reads (18.52–117.30) | 16-CPU quota | Maximum method-stage cgroup memory peak; 256-GiB allocation; median 62.63 GiB (37.62–88.66) | Evaluated | 8 completed single-cell runs; no out-of-memory failure |
| SCAPTURE | Source version 8845e84 | Command-line pipeline (scapture -m PAScall / PASmerge / PASquant) | PAScall -w 400, overlap 0.5; no –polyaDB; DeepPASS-positive exonic and intronic sites | Prepared CB/UB BAM; SCAPTURE annotation from the same GTF | PAScall + PASmerge + PASquant | 16-CPU quota; 16 threads | Cgroup memory peak; 256-GiB allocation | Not evaluated | 8 completed single-cell runs; run serially; no out-of-memory failure |
| SCAPE | 1.0.0 | Python script (apamix) | apamix on the coordinate-sorted GENCODE GTF UTR BED; min/max APA per UTR 1/5; defaults otherwise; official collapse (group_pa.py; group threshold 100 nt) | Prepared CB/UB BAM; UTR BED from the sorted GTF | apamix wall time using the sorted GTF; official group_ pa.py collapse run downstream | 16-CPU quota | Cgroup memory peak; 256-GiB allocation | Not evaluated | 8 completed single-cell runs; run serially; no out-of-memory failure |
| MAAPER | 1.1.1 | R package (scripted) | PolyA_DB PAS reference; min gene reads 50; min PAS reads 25; min PAS fraction 0.05; single-cell benchmark: single-sample mode; run=all | Prepared BAM; PolyA_DB PAS and exon RDS from the GTF; brain Visium: two selected clusters from k=3 clustering, two pseudoreplicates per cluster | Native MAAPER method stage; median 58.36 s/M aligned reads (17.29–336.87); reference-based | 16-CPU quota | Method-stage cgroup memory peak; 256-GiB allocation; median 44.19 GiB (23.82–116.02) | Evaluated | 8 completed single-cell runs; no out-of-memory failure |
| scUTRquant | 0.5.0 | Snakemake workflow | target utrome_hg38_v1 (mm10_v2 for mouse); min UMIs 1; correct BUS true; output txs, genes | BAM (bam) or tagged-BAM FASTQ (Microwell-seq, Stereo-seq); UTRome target | Native workflow only; excludes BAM-to-FASTQ adaptation and configuration | 16-CPU quota | Method-stage cgroup memory peak; 256-GiB allocation | Evaluated | 8 completed single-cell runs; no out-of-memory failure |

### Supplementary Table 4 | scPolyA rerun times

| Dataset | Comparison | Tail threshold | Endpoint | Logged stage span (s) | Logged stage span (min) | Span / fresh (%) | Tasks executed / total | Full invocation (s) | Run conditions | Timing definition and run design |
|---|---|---|---|---|---|---|---|---|---|---|
| 10x PBMC | Fresh run | 1 | Complete count matrix and APA summaries | 3795 | 63.3 | 100.0 | 228 / 228 | 3799.46 | 16 CPUs; 256 GiB; no overlapping timed work; no out-of-memory failure. | Logged stage span follows the Fig. 6c stage boundaries through the count matrix and APA summaries; excludes invocation startup and finalization. Fresh baseline. |
| 10x PBMC | Unchanged parameters | 1 → 1 | Complete count matrix and APA summaries | 87 | 1.5 | 2.3 | 0 / 228 | 91.47 | 16 CPUs; 256 GiB; no overlapping timed work; no out-of-memory failure. | Logged stage span follows the Fig. 6c stage boundaries through the count matrix and APA summaries; excludes invocation startup and finalization. Independent copy of the completed fresh-run directory; moved-run path repair enabled. All stages completed. |
| 10x PBMC | Threshold change | 1 → 3 | Complete count matrix and APA summaries | 1740 | 29.0 | 45.8 | 60 / 228 | 1744.00 | 16 CPUs; 256 GiB; no overlapping timed work; no out-of-memory failure. | Logged stage span follows the Fig. 6c stage boundaries through the count matrix and APA summaries; excludes invocation startup and finalization. Independent copy of the completed fresh-run directory; moved-run path repair enabled. All stages completed. |

### Supplementary Table 5 | cSCC 3′UTR candidates

| Gene | Tier | KC Δ logit | KC Δ 3′UTR (nt) | KC patients (lower/n) | KC P | KC FDR | Tissue Δ logit | Tissue sections (tumour ≤ TME / n) | Tissue patients | Tissue P | Myeloid control Δ logit (P) | Composition control Δ logit (P) |
|---|---|---|---|---|---|---|---|---|---|---|---|---|
| ILF3 | 1 | -0.66 | -420 | 9/9 | 2.7e-06 | 0.0026 | -0.39 | 9/10 | 5 | 0.0065 | -0.37 (P 0.084) | 0.22 (P 0.21) |
| PDCD6IP | 1 | -0.88 | -180 | 9/9 | 8.4e-06 | 0.0031 | -0.76 | 9/9 | 4 | 2.0e-05 | -0.41 (P 0.16) | -0.19 (P 0.39) |
| LZIC | 1 | -0.65 | -481 | 9/9 | 2.1e-05 | 0.0038 | -0.51 | 7/8 | 4 | 0.003 | -0.31 (P 0.069) | 0.16 (P 0.093) |
| PXMP4 | 1 | -0.98 | -502 | 9/9 | 3.4e-04 | 0.013 | -0.58 | 7/7 | 3 | 0.0015 | untestable | 0.55 (P 0.0012) |
| APLP2 | 1 | -0.74 | -123 | 10/10 | 4.5e-04 | 0.014 | -0.26 | 10/10 | 5 | 3.2e-04 | -0.16 (P 0.45) | -0.08 (P 0.52) |
| FAM199X | 1 | -0.60 | -632 | 9/9 | 4.4e-04 | 0.014 | -0.66 | 7/7 | 3 | 0.0045 | -0.25 (P 0.19) | -0.20 (P 0.29) |
| PRKRIP1 | 1 | -0.65 | -182 | 8/8 | 0.0014 | 0.024 | -0.50 | 7/7 | 3 | 0.0061 | -0.33 (P 0.18) | 0.02 (P 0.88) |
| MR1 | 1 | -1.20 | -954 | 8/8 | 0.0016 | 0.026 | -0.74 | 8/8 | 4 | 0.0075 | -0.44 (P 0.18) | -0.32 (P 0.27) |
| MCL1 | 1 | -0.57 | -174 | 9/10 | 0.0017 | 0.027 | -0.23 | 8/10 | 5 | 0.0067 | -0.06 (P 0.69) | 0.16 (P 0.33) |
| TIPRL | 1 | -0.55 | -142 | 8/9 | 0.0022 | 0.031 | -0.38 | 8/9 | 4 | 0.0059 | -0.02 (P 0.94) | -0.13 (P 0.33) |
| YIPF4 | 1 | -1.01 | -539 | 9/9 | 0.0022 | 0.031 | -0.63 | 8/9 | 4 | 0.009 | -0.10 (P 0.57) | -0.51 (P 0.083) |
| IGF1R | 1 | -0.26 | -385 | 8/9 | 0.0033 | 0.038 | -0.47 | 7/8 | 4 | 0.0066 | untestable | -0.16 (P 0.46) |
| SREBF2 | 1 | -0.30 | -64 | 9/9 | 0.0038 | 0.041 | -0.34 | 9/9 | 4 | 0.0011 | -0.34 (P 0.39) | 0.10 (P 0.23) |
| PEA15 | 2 (comp.) | -0.57 | -80 | 9/9 | 2.9e-04 | 0.013 | -0.49 | 9/10 | 5 | 0.0038 | -0.14 (P 0.39) | -0.84 (P 4.7e-05) |
| SLC35F5 | 2 (comp.) | -0.87 | -1387 | 9/9 | 4.6e-04 | 0.014 | -0.45 | 7/7 | 3 | 0.0019 | 0.19 (P 0.15) | -0.42 (P 6.4e-04) |
| CIAO1 | 2 (comp.) | -0.78 | -393 | 9/9 | 0.0011 | 0.022 | -0.51 | 8/9 | 4 | 0.0014 | -0.35 (P 0.1) | -0.46 (P 0.003) |
| TIMP3 | 2 (comp.) | -0.90 | -455 | 8/8 | 0.0013 | 0.023 | -0.70 | 8/8 | 4 | 0.0019 | untestable | -0.26 (P 0.043) |
| AGFG1 | 2 (comp.) | -0.72 | -247 | 9/9 | 0.0013 | 0.023 | -0.56 | 4/4 | 2 | 0.0016 | -0.04 (P 0.93) | -0.54 (P 0.0067) |
| RRAS2 | 2 (comp.) | -0.53 | -102 | 8/9 | 0.0014 | 0.024 | -0.59 | 4/4 | 2 | 0.0054 | untestable | -0.35 (P 0.011) |
| RMDN1 | 2 (FDR) | -0.60 | -176 | 9/9 | 0.006 | 0.051 | -0.72 | 8/8 | 4 | 6.0e-04 | -0.27 (P 0.27) | 0.02 (P 0.89) |
| WDR1 | 2 (both) | -0.27 | -50 | 8/10 | 0.0068 | 0.055 | -0.19 | 10/11 | 5 | 0.0091 | -0.17 (P 0.18) | -0.15 (P 0.012) |
| NLE1 | 2 (FDR) | -0.86 | -233 | 8/9 | 0.007 | 0.056 | -0.60 | 7/7 | 3 | 0.003 | untestable | 0.79 (P 0.022) |
| SAP30L | 2 (FDR) | -0.76 | -847 | 7/8 | 0.0077 | 0.058 | -0.50 | 8/8 | 4 | 0.0036 | -0.45 (P 0.22) | 0.12 (P 0.68) |
| STAT3 | 2 (FDR) | -0.35 | -172 | 9/10 | 0.0078 | 0.058 | -0.35 | 8/8 | 4 | 0.0024 | -0.00 (P 0.98) | 0.26 (P 0.011) |
| MED13L | 2 (FDR) | -0.18 | -33 | 9/9 | 0.0083 | 0.06 | -0.33 | 7/8 | 4 | 0.0083 | 0.17 (P 0.11) | -0.06 (P 0.57) |
| RRP15 | 2 (FDR) | -0.49 | -255 | 8/9 | 0.0089 | 0.063 | -1.09 | 4/4 | 2 | 0.0091 | -0.03 (P 0.95) | 0.06 (P 0.86) |
| NCBP1 | 2 (FDR) | -0.41 | -186 | 7/8 | 0.01 | 0.068 | -0.37 | 8/8 | 4 | 0.0097 | -0.03 (P 0.9) | 0.01 (P 0.94) |
| YIPF6 | 2 (FDR) | -0.57 | -180 | 8/9 | 0.01 | 0.068 | -0.72 | 7/8 | 4 | 0.0019 | -0.25 (P 0.38) | -0.03 (P 0.92) |
| CYBRD1 | 2 (FDR) | -0.88 | -541 | 7/8 | 0.011 | 0.07 | -0.79 | 4/4 | 2 | 0.0068 | 0.02 (P 0.94) | 0.06 (P 0.77) |
| SLCO4A1 | 2 (FDR) | -0.38 | -7 | 7/8 | 0.013 | 0.078 | -1.25 | 7/7 | 4 | 0.0058 | -0.01 (P 0.98) | -0.36 (P 0.32) |
| CALU | 2 (FDR) | -0.40 | -93 | 8/9 | 0.013 | 0.078 | -0.45 | 10/10 | 5 | 8.7e-04 | 0.09 (P 0.85) | -0.06 (P 0.63) |
| CD2BP2 | 2 (FDR) | -0.53 | -161 | 8/9 | 0.015 | 0.087 | -0.46 | 8/9 | 4 | 0.0015 | -0.13 (P 0.35) | 0.28 (P 0.042) |
| NDC1 | 2 (FDR) | -0.43 | -240 | 8/9 | 0.017 | 0.093 | -0.47 | 6/7 | 3 | 0.0044 | -0.39 (P 0.096) | 0.47 (P 0.064) |
| USP15 | 2 (FDR) | -0.52 | -37 | 7/9 | 0.017 | 0.093 | -0.68 | 7/8 | 4 | 0.0053 | 0.27 (P 0.5) | -0.34 (P 0.1) |
| TBC1D22A | 2 (FDR) | -0.41 | -75 | 8/9 | 0.018 | 0.094 | -0.41 | 8/9 | 4 | 0.0097 | 0.06 (P 0.8) | 0.07 (P 0.8) |
| ARIH2 | 2 (FDR) | -0.46 | -190 | 8/9 | 0.02 | 0.1 | -0.56 | 7/7 | 3 | 7.1e-04 | -0.18 (P 0.41) | 0.02 (P 0.86) |
| STAM2 | 2 (FDR) | -0.55 | -360 | 7/9 | 0.021 | 0.1 | -0.43 | 7/7 | 3 | 0.0039 | -0.15 (P 0.54) | -0.04 (P 0.89) |
| GFPT1 | 2 (FDR) | -0.41 | -503 | 6/8 | 0.022 | 0.11 | -0.63 | 7/7 | 3 | 9.3e-04 | -0.11 (P 0.69) | 0.09 (P 0.53) |
| PLXNA1 | 2 (FDR) | -0.36 | -104 | 7/9 | 0.023 | 0.11 | -0.44 | 9/9 | 4 | 0.0046 | -0.27 (P 0.5) | 0.19 (P 0.24) |
| RANBP3 | 2 (FDR) | -0.34 | -80 | 7/9 | 0.026 | 0.12 | -0.35 | 7/8 | 4 | 0.0065 | -0.02 (P 0.85) | 0.15 (P 0.2) |
| FBXL4 | 2 (FDR) | -0.37 | -326 | 7/8 | 0.027 | 0.12 | -1.05 | 4/4 | 2 | 0.0034 | -0.19 (P 0.65) | 0.03 (P 0.84) |
| OGFRL1 | 2 (FDR) | -0.38 | -622 | 6/9 | 0.028 | 0.12 | -0.50 | 7/8 | 4 | 0.0099 | 0.11 (P 0.49) | -0.13 (P 0.33) |
| BID | 2 (FDR) | -0.37 | -28 | 8/9 | 0.028 | 0.12 | -0.52 | 7/7 | 3 | 0.0015 | -0.12 (P 0.38) | 0.04 (P 0.71) |
| RBM28 | 2 (FDR) | -0.51 | -409 | 7/8 | 0.028 | 0.12 | -0.77 | 7/7 | 3 | 0.0014 | 0.02 (P 0.96) | -0.43 (P 0.11) |
| PPP6R2 | 2 (FDR) | -0.29 | -46 | 7/9 | 0.035 | 0.14 | -0.40 | 8/8 | 4 | 0.0094 | -0.19 (P 0.39) | 0.24 (P 0.057) |
| GDE1 | 2 (FDR) | -0.22 | -89 | 8/9 | 0.039 | 0.15 | -0.29 | 7/9 | 4 | 0.0098 | -0.13 (P 0.38) | -0.06 (P 0.47) |
| SERPINB8 | 2 (both) | -0.45 | -209 | 8/9 | 0.041 | 0.15 | -0.79 | 8/9 | 5 | 0.0024 | -0.05 (P 0.78) | -0.22 (P 0.012) |
| FGFR1OP2 | 2 (FDR) | -0.14 | -24 | 8/9 | 0.044 | 0.16 | -0.92 | 4/4 | 2 | 0.0073 | -0.00 (P 0.99) | -0.22 (P 0.22) |
| PARVA | 2 (FDR) | -0.74 | -773 | 7/9 | 0.045 | 0.16 | -0.61 | 7/8 | 4 | 0.0066 | untestable | -0.40 (P 0.3) |
| TTC1 | 2 (FDR) | -0.36 | -16 | 7/10 | 0.047 | 0.16 | -0.40 | 8/8 | 4 | 0.0097 | -0.05 (P 0.72) | -0.07 (P 0.33) |

### Supplementary Table 6 | scPolyA output files

| File | Level | Contents | Mode | Columns |
|---|---|---|---|---|
| inputs.csv | run | Copy of –inputs (one row per sample BAM) | Input manifest | sample_id, source_path, is_deduped, requires_split |
| preprocess_inputs.csv | run | Preprocessing plan per sample and contig | Input manifest | pp_id, pp_path, sample_id, source_type, contig_name, region_label, dedup_status, is_deduped, requires_split, barcode_suffix |
| unit_inputs.<dedup_method>.csv | run | Deduplicated unit BAMs; use resolved_path | Input manifest | unit_id, pp_id, sample_id, source_type, contig_name, is_deduped_unit, dedup_status, barcode_suffix, resolved_path |
| peaks-mispriming-audited.terminal-aware.v1.bed | site | Audited final peaks (headerless BED) | Terminal-aware | chrom, start, end, name, max_cov, strand, total_polyA_reads, misprimed_ratio, total_reads |
| annotated.terminal-aware.v2.bed | site | Annotated peaks, 16-column BED (BigBed / GTF alongside) | Terminal-aware | chrom, start, end, peak_name, max_cov, strand, total_polyA_reads, misprimed_ratio, total_reads, gene_name, distance_to_gene, canonical_transcript, stop_codon_position, distance_to_stop_codon, feature, gene_id |
| annotated.terminal-aware.v2.gtf | site | Counting GTF; attributes listed in the next row | Terminal-aware | seqname, source, feature, start, end, score, strand, frame, attribute |
| annotated.terminal-aware.v2.gtf:attribute | site | GTF attribute keys | Terminal-aware | peak_name, gene_name, canonical_transcript, stop_codon_position, distance_to_gene, distance_to_stop_codon, misprimed_ratio, feature, gene_id |
| terminal_aware_peak_to_gene.v2.tsv.gz | site | Audited peak metadata and gene mapping | Terminal-aware | peak_name, gene_name, gene_id, chrom, start, end, peak_length, strand, feature, canonical_transcript, stop_codon_position, distance_to_gene, distance_to_stop_codon, three_prime_utr_length, max_cov, total_polyA_reads, total_reads, misprimed_ratio, general_mispriming_pass, intronic_mispriming_pass, mispriming_pass, peak_order_in_gene, n_peaks_in_gene, is_distal_peak |
| xr_to_peak.terminal-aware.v2.tsv.gz | read | Peak sidecar: read → assigned peak | Terminal-aware | query_name, flag, feature |
| counts.terminal-aware.v2.tsv.gz | site × cell | Per-sample peak counts, long format | Terminal-aware | peak_name, cell_barcode, umi_count |
| terminal_aware_gene_cell.v2.tsv.gz | gene × cell | Terminal gene-cell UMI counts, long format | Terminal-aware | gene_id, gene_name, strand, terminal_gene_unit_id, cell_barcode, gene_umi_count |
| terminal_aware_cell_gene.matrix.v2.tsv.gz | cell × gene | Dense terminal cell-gene matrix (size-limited) | Terminal-aware | cell_barcode, one column per terminal_gene_unit_id |
| terminal_aware_psi.v2.tsv.gz | site × cell | Terminal PSI | Terminal-aware | peak_name, gene_id, gene_name, strand, terminal_gene_unit_id, cell_barcode, peak_umi_count, gene_umi_count, three_prime_utr_length, psi |
| terminal_aware_weighted_three_prime_utr_length.v2.tsv.gz | gene × cell | Weighted terminal 3′UTR length | Terminal-aware | gene_id, gene_name, strand, terminal_gene_unit_id, cell_barcode, gene_umi_count, weighted_three_prime_utr_length |
| intronic_ipui_per_site.v2.tsv.gz | site × unit | Per-site intronic usage (iPUI) | Terminal-aware | gene_id, gene_name, intronic_pas_id, chromosome, coordinate, strand, quantification_unit_id, I_i, U, iPUI_i, misprimed_ratio, mispriming_pass |
| intronic_ipui_gene_total.v2.tsv.gz | gene × unit | Gene-total intronic usage (iPUI) | Terminal-aware | gene_id, gene_name, strand, quantification_unit_id, retained_intronic_pas_count, I_total, U, total_iPUI |
| rna_gene_cell.tsv.gz | gene × cell | RNA gene-cell counts, long format | Optional RNA | gene_name, cell_barcode, gene_umi_count |
| rna_cell_gene.matrix.tsv.gz | cell × gene | Dense RNA cell-gene matrix (size-limited) | Optional RNA | cell_barcode, one column per gene_name |
